# BioPhasor: Decoding Cellular State Tensors from Multi-Omics Phasor Dynamics for Quantum Ready Systems Biology

**DOI:** 10.64898/2026.07.17.739210

**Authors:** Dibakar Sigdel, Namuna Panday

## Abstract

Integrating multi-omics data—transcriptomics, proteomics, metabolomics, single-cell—remains a fundamental challenge in systems biology. We present BioPhasor, a framework that encodes each measurement as a complex phasor *z* = *e^iϕ^* on the compact *N* -torus T*^N^*, modelling the cell as phase-coupled oscillatory programs whose dissipative dynamics generate limit cycles and an attractor landscape. From this geometry we derive the Cell State Tensor (CST), a rank-3 tensor whose axes we root in measured multi-omics quantities: a pathway/module atlas on the regulatory axis and a directional central-dogma modality axis. Across nine scenarios on open public data (GEO, CPTAC), loaded through one unmodified data layer, we report verdicts honestly: four reproduce, three are partial, two do not. A data-driven cell-cycle axis lifts agreement with a reference method from 0.34 to 0.69; an explicit circadian origin cuts peak-time error from 10.6 to 1.4 h; and central-dogma coupling—mRNA phase organising protein amplitude—clears a surrogate null and is tumour-specific. Grounding the quantum-ready claim, the CST maps to a density-matrix formalism whose coherence and entropy match quantum-information counterparts, and the phasor circuit transpiles gate-for-gate to a variational quantum circuit, though no empirical advantage emerges. A single loader regenerates every reported number, and the code is released.

## I. INTRODUCTION

The advent of high-throughput sequencing has generated an unprecedented multi-dimensional view of cellular state [1–5]. By simultaneously profiling the transcriptome (RNA-seq), epigenome (ATAC-seq), proteome, and metabolome, researchers aim to reconstruct the hierarchical regulatory networks governing biological life. However, integrating these disparate data modalities remains a primary challenge in computational biology [6].

### A. Motivation: From Clockwork Biology to Geometric Integration

A recurring obstacle in multi-omics computation is scale heterogeneity: the modalities operate on fundamentally different numerical ranges—RNA counts (∼ 10^4^) versus TF ChIP-seq (∼ 10^0^) versus metabolite concentrations (*µ*M)—so that combining them requires aggressive normalization. A second, more subtle obstacle is geometric: existing integration methods (MOFA+, SNF, CCA, scVI) impose Euclidean structure on data that is inherently circular, since biological states from mitosis to circadian rhythms are closed oscillatory trajectories rather than linear endpoints.

**BioPhasor** addresses both by representing each omic feature as a phase angle *ϕ* ∈ (−*π, π*] on the unit circle and collecting the full multi-omics state on the compact *N* -Torus T*^N^*. This circular representation is simultaneously scale-invariant, topology-aware, and compatible with well-developed circular statistics.

#### 1. The Legacy of Biological Oscillators

The recognition that cellular life is a rhythmic process traces to early circadian and ultradian oscillator models [7, 8]. These models captured the periodic “clockwork” of negative feedback loops—the suprachiasmatic nucleus (SCN) and cell cycle—via coupled differential equations. Yet they were inherently low-dimensional, confined to manually curated genes such as *Period* (*Per*) and *Cryptochrome* (*Cry*).

#### 2. The Multi-Omics Expansion and the Euclidean Bottleneck

With high-throughput sequencing, modeling shifted to **Euclidean deep learning** [9–18]. Variational Autoencoders [19] and Transformers [20] cluster cell types and predict phenotypes effectively, but they treat gene expression as a linear magnitude in R*^N^*. This abstraction introduces three fundamental errors for biological data:

1. **Periodicity mismatch.** Biological states—from mitosis to metabolic pulses—are circular. In a linear model, the start and end of a cycle appear as antipodal points rather than adjacent states on a manifold.
2. **Saturation blindness.** Biological signals saturate (Michaelis-Menten kinetics [21]); once a promoter is occupied, additional concentration does not increase output. Euclidean activations require complex non-linear engineering to approximate this threshold behaviour.
3. **Cross-omic scale heterogeneity.** Combining RNA counts with TF ChIP-seq peaks creates numerical instability, demanding aggressive normalization that destroys biologically meaningful signal.

#### 3. Phase Dynamics: The Missing Link

The most robust framework for periodic biological systems has long been **phase dynamics** [22–27]. EEG studies and circadian biology consistently show that the *timing* (phase) of a signal carries more regulatory information than its *amplitude*. The “Oscillatory Cell” hypothesis underlies BioPhasor: every omic feature is a phasor *e^iϕ^* rotating on a high-dimensional *N* -Torus T*^N^* .

#### 4. BioPhasor: Scope and Contributions

- **Unified multi-omics phasor API.** A single library covers RNA-seq, scRNA-seq, proteomics, metabolomics, and multi-omics fusion with modality-aware encoding strategies.
- **Riemannian manifold operations.** Geodesic distance, Fréchet mean, log/exp maps, and geodesic interpolation give topologically correct statistics on T*^N^*, replacing ad hoc Euclidean approximations.
- **Oscillator dynamics.** The BioKuramoto model captures gene regulatory network synchrony as coupled phase oscillators; the circadian phasor module extracts phase, amplitude, and rhythmicity score via the Biological Phasor Transform (BPT).
- **Cell-state assignment.** The cell-cycle phasor module assigns G1/S/G2/M from marker gene phasors; the same approach supports custom state spaces (EMT, senescence, quiescence).
- **Phasor machine learning.** Variational Phasor Circuits (VPC), Phasor Transformers, and the Large Phasor Model (LPM) provide a full hierarchy of phase-native architectures with circular loss functions.
- **Dissipative dynamics and attractor landscape.** Cells are modelled as phase-coupled dissipative oscillators whose collective dynamics generate proliferating limit cycles. The attractor geometry—topology, basin structure, and transition dynamics— is formalized via the Waddington quasi-potential and Floquet stability analysis (Section III).
- **Cell State Tensor (CST).** A rank-3 tensor X*_t_* ∈ R*^R^*^×^*^T^* ^×^*^H^* encoding regulatory, temporal, and homeostatic cell state, derived from attractor-geometric features (global coherence, phase entropy, Floquet resilience, Markov transition entropy)—the canonical latent field summarizing the cell’s attractor landscape (Section IV). Its axes are rooted in *measured* multi-omics quantities—a pathway/module atlas on the regulatory axis and a directional central-dogma modality axis—rather than abstract categories (Section IV B).
- **Open ecosystem.** Comprehensive documentation and validated experimental pipelines for research reproducibility.

By grounding BioPhasor in circadian oscillator theory, dissipative dynamics, and the compact geometry of T*^N^*, we provide a unified toolkit for multi-omics analysis that treats regulatory timing, rather than expression magnitude, as the primary signal.

*a. What is specific to multi-omics.* Phase-geometric state tensors have been proposed in other domains—most directly for neural recordings, where a spatiotemporal tensor is indexed by spectral band, anatomical region, and time. It would be easy to read the CST as a transposition of that idea onto genes, and an earlier framing indeed defined it largely by analogy. We take the opposite view: the value of a phasor state tensor lies entirely in whether its axes are the measured, structured quantities of *its own* domain, and multi-omics offers two that have no neural counterpart. First, the regulatory axis is organised by a *pathway/module atlas*: genes group into curated functional programs (here the MSigDB Hallmark collection [28]) exactly as voxels group into anatomical regions, and we show (Section VI K) that this biological grouping—not the analogy—is what makes the CST compressible and its factors interpretable. Second, multi-omics has a *directional* axis with no analogue in a spatial neural tensor: the central dogma (DNA → mRNA → protein → metabolite), along which cross-modal phase coupling encodes regulation such as translational buffering (Section VI L). These two properties—a biological atlas for the regulatory axis and a causal modality axis—are what root BioPhasor in the structure of multi-omics data rather than in a borrowed template.

## II. THEORY: PHASOR DYNAMICS ON THE N -TORUS

This section establishes the mathematical framework underlying the BioPhasor framework. We present: (1) the *N* -Torus state space and phasor encoding; (2) circular statistics and coherence theory; (3) the Riemannian manifold geometry of T*^N^* ; (4) coupled oscillator dynamics (Kuramoto); and (5) the Biological Phasor Transform for time-series analysis.

### A. State Space: The *N* -Torus and Phasor Encoding

In traditional bioinformatics, a cell’s regulatory state is modeled as ***x*** ∈ R*^N^* [29–33], where each component carries arbitrary magnitude noise. BioPhasor instead defines the state space for *N* omic features as the continuous *N* -Torus:

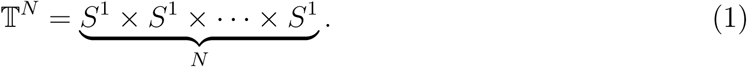

Each cell’s multi-omic state is represented as a complex phasor vector:

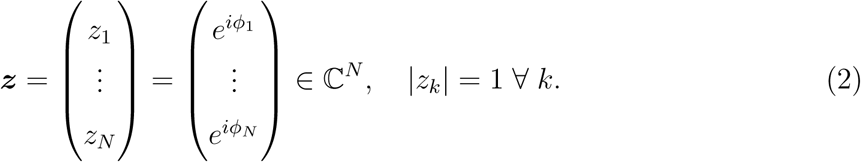

**Definition II.1** (Omic Phasor Encoding). Given a normalized feature value x_j_ (e.g., CPM-normalized RNA count), define the tanh-phase encoding map:

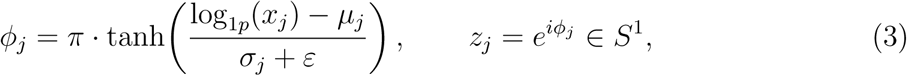

*where µ_j_, σ_j_ are gene-wise mean and standard deviation, and ε >* 0 *ensures numerical stability. By construction, ϕ_j_* ∈ (−*π, π*)*. For methylation β-values (log-linear encoding) and log2-scaled proteomics (linear encoding), modality-specific mappings are applied (Section II B)*.

This geometric embedding provides two key biological invariances:

- **Scale invariance**: phase is immune to multiplicative magnitude scaling (sequencing depth, batch effects).
- **Manifold compactness**: T*^N^* is bounded, eliminating exploding gradients in deep architectures.

Figure 1 illustrates the encoding pipeline.

**FIG. 1:**
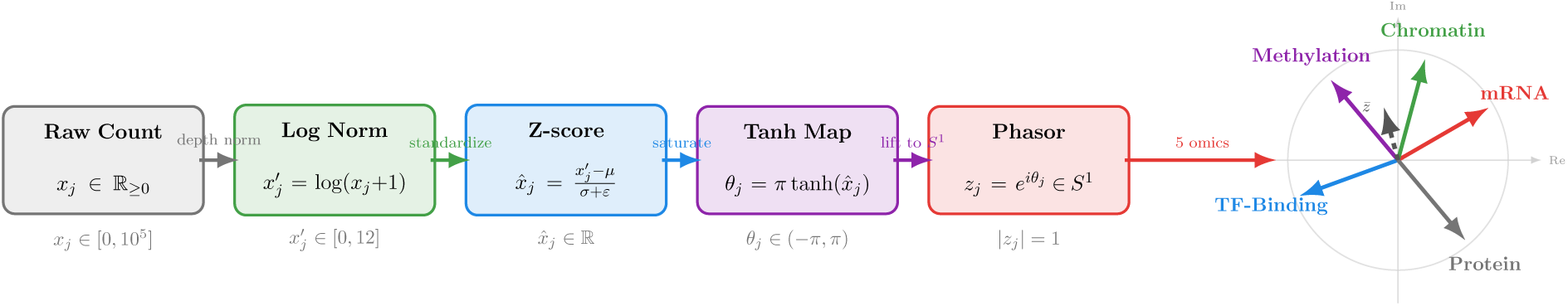
BioPhasor Omic Phasor Encoding Pipeline. Raw counts are CPM-normalized, log-transformed, z-scored, then compressed to (−*π, π*) via tanh and lifted to *S*^1^ via *e^iϕ^*. *Right* : Five example phasors from transcriptomics (red), epigenomics (purple), chromatin (green), TF binding (blue), and their circular mean *z̄* (dashed).

### B. BioPhasor Encoding Strategies

Different omics modalities have distinct distributional characteristics requiring tailored encoding maps. BioPhasor implements three strategies:

*a. Tanh-Phase Encoding (RNA-seq, ATAC-seq).* For right-skewed count data, the canonical encoding applies log1p transformation followed by per-gene z-scoring and tanh saturation:

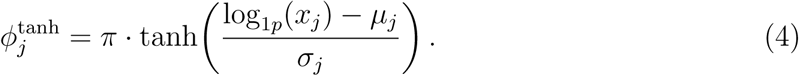

Among the three encoders this achieves the widest phase spread (*σ_ϕ_*≈ 1.995 radians), which maximizes the angular resolution available to the downstream coherence statistics.

*b. Log-Linear Encoding (DNA Methylation β-values).* Methylation *β* ∈ [0, 1] maps linearly to (−*π, π*) via the logit:

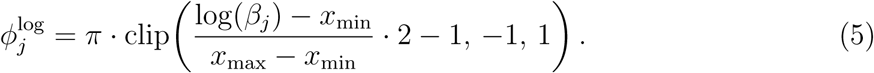

*c. Linear Encoding (Proteomics, log2-scaled).* For approximately symmetric log2- transformed data:

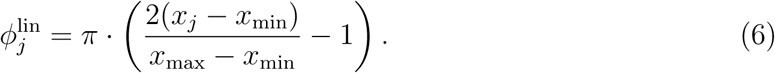

The auto-dispatch encoder selects the correct strategy based on modality name:

### C. Circular Statistics of Phasor Distributions

**Definition II.2** (Phase Coherence (Mean Resultant Length)). *For N phasors* 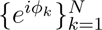 *, the phase coherence is:*

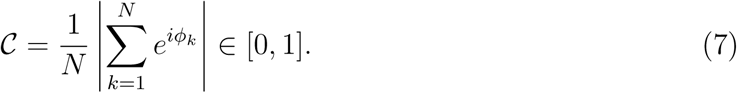

C = 1 *iff all phasors are co-aligned;* C ≈ 0 *under uniformly random phases*.

**Theorem II.1** (Coherence as a Systems-Order Biomarker). The scalar C *monotonically increases with the alignment of phasors on S*^1^*. In multi-omics,* C *quantifies the synchronization of regulatory programs across genes or samples*.

**Corollary II.2** (Disease-State Stratification via Coherence). The framework predicts that phase coherence C̄ *stratifies disease states differing in regulatory coordination: e.g. quiescent IGHV-mutated versus autonomously-signaling IGHV-unmutated CLL are expected to separate by mean coherence, reflecting their distinct BCR-signaling phenotypes. This is a testable prediction to be evaluated on labelled real cohorts (Section VI F, Section VI O)*.

*a. Rayleigh Test for Phase Uniformity.* A gene is declared *phase-coherent* if the Rayleigh test rejects the null hypothesis of uniform phase distribution:

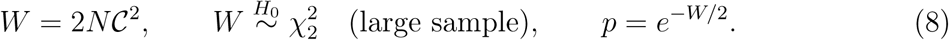

BioPhasor implements this test.

*b. Fréchet Mean (Circular Mean).*The unbiased circular mean of phasors {*ϕ_k_*} is:

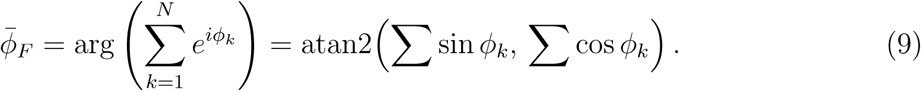

This Fréchet mean replaces the arithmetic mean wherever phase data are circularly distributed.

### D. Riemannian Geometry of T*^N^*

**Definition II.3** (Geodesic Distance on T*^N^*). The geodesic distance between two phasors ϕ_1_*, ϕ*_2_ ∈ *S*^1^ *is the arc-length:*

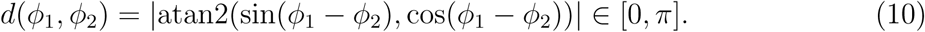

*For N-dimensional phasor vectors:*

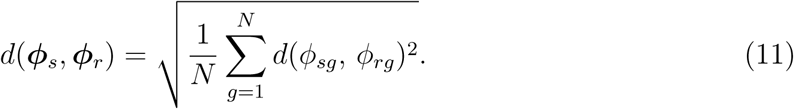

*a. Log/Exp Maps for Tangent-Space Analysis.* The log map projects T*^N^* to the flat Euclidean tangent space at a base point ***µ***, enabling unbiased linear operations (PCA, regression):

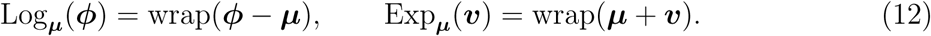

These are exact inverses on T*^N^* (no approximation needed).

*b. Geodesic Interpolation (Phasor Pseudo-Time).* For samples ***ϕ_A_*** and ***ϕ****_B_* (e.g., starting and terminal cell states), the geodesic trajectory parameterized by *t* ∈ [0, 1] is:

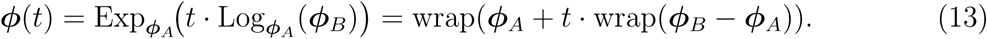

This provides a parameter-free pseudo-time for single-cell trajectories.

### E. Unitary Operations: Regulatory Interference

Once cell states are encoded on T*^N^*, regulatory interactions are modeled by unitary operators:

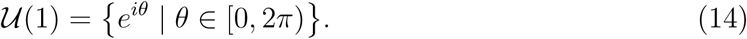

**Definition II.4** (Cross-Layer Interference Matrix). *For* ***ϕ*** ∈ T*^N^, the pairwise regulatory interaction matrix I* ∈ [−1, +1]*^N^*^×^*^N^ is:*

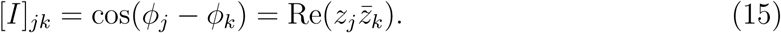

*I_jk_* = +1 *implies co-activation; I_jk_*= −1 *implies mutual repression; I_jk_* ≈ 0 *indicates orthogonal independence*.

The pull-back operator restores manifold compactness after linear mixing:

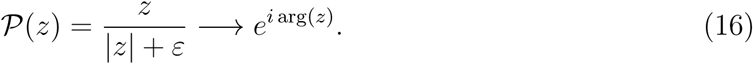

### F. Variational Phasor Circuit (VPC)

The VPC implements biological regulation via two gate types:

1. **Parameterized Shift Gates**: learnable phase rotations *S_k_*(*θ_k_*) = diag(1*, …, e^iθ^k, …,* 1).
2. **Entangling Mix Gates**: 50:50 beam-splitter 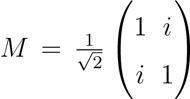 modelling pairwise omic cross-talk.

A single VPC layer: ***z****_t_*_+1_ = P(*M S*(***θ***) ***z****_t_*). Parameter scaling: O(*LN*) for depth *L* and width *N*, vs. O(*N* ^2^) for dense Euclidean layers.

### G. Quantum Parallel: VPC as a Classically-Simulated VQC

Because the VPC acts on unit-modulus phasors through phase rotations and norm-preserving mixing, its gate algebra maps one-to-one onto a Variational Quantum Circuit (VQC) [34, 35]. Each parameterized shift *S_k_*(*θ_k_*) = diag(1*, …, e^iθ^k, …,* 1) becomes a single-qubit *R_z_*(*θ_k_*) rotation; the entangling Mix gate becomes a CNOT+*R_z_* entangling layer; and the DFT token-mixing block becomes a Quantum Fourier Transform (QFT). The VPC therefore operates on the phase-diagonal sector of an *N* -qubit Hilbert space, and its trained parameters are directly usable as quantum rotation angles without reparametrization—the multi-omics analogue of the correspondence established for phasor circuits in neural signals. Table I lists the gate mapping; Section VII evaluates it and its complexity consequences on real multi-omics data.

**TABLE I:**
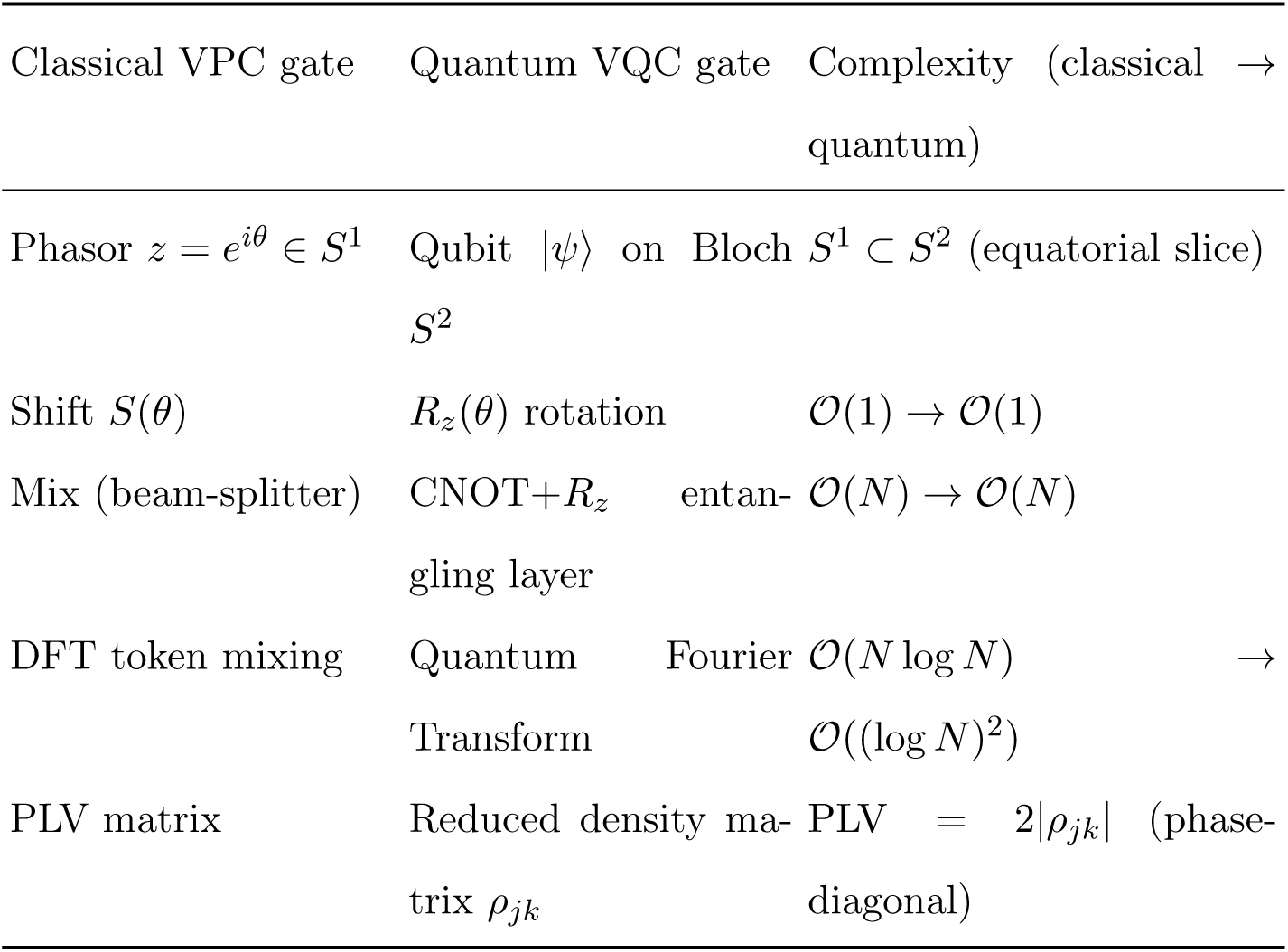
Gate-level correspondence between the classical Variational Phasor Circuit and a Variational Quantum Circuit. The mapping is exact on the phase-diagonal sector; complexity columns give per-operation asymptotic cost in system size *N* (number of gene/pathway phasors, i.e. qubits).

**TABLE II:**
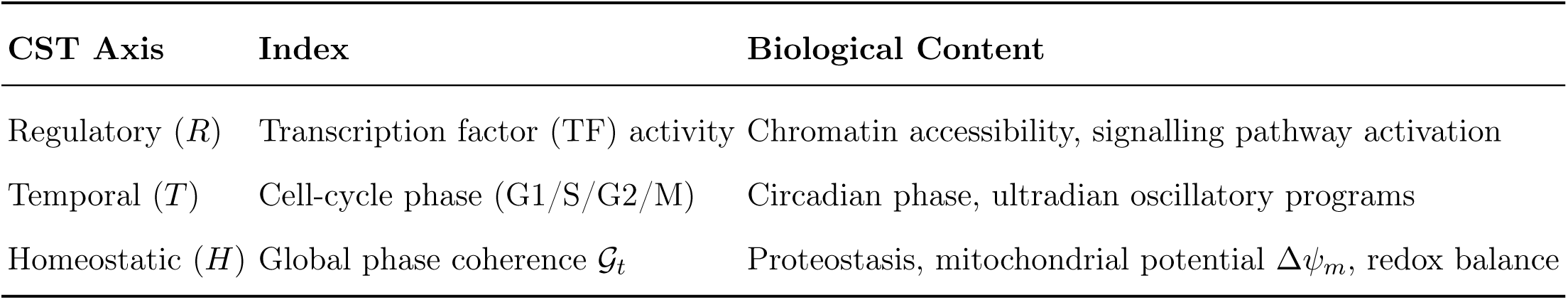
Biological semantics of the three CST axes.

### H. Coupled Oscillator Dynamics: Kuramoto Model

Gene regulatory networks are modeled as coupled phase oscillators via the Kuramoto model:

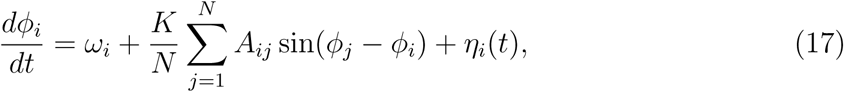

where *ω_i_* is the natural oscillation frequency of gene *i*, *K* is global coupling strength, *A_ij_* is the GRN adjacency matrix, and *η_i_*(*t*) ∼ N(0*, σ*^2^) is Gaussian noise.

The order parameter 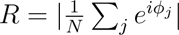 measures network-wide synchrony. A phase transition occurs at critical coupling

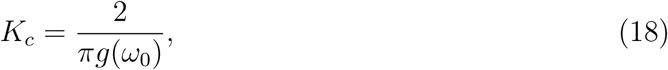

where *g*(*ω*_0_) is the frequency distribution at the mean. BioPhasor maps (*K, σ*) parameter space to *R*_∞_, quantifying synchrony robustness in realistic GRN topologies (hub-spoke, scale-free).

### I. Biological Phasor Transform (BPT) for Time-Series

For oscillatory time-series (circadian, cell cycle), the BPT extracts phase and amplitude at biological frequencies via a single DFT:

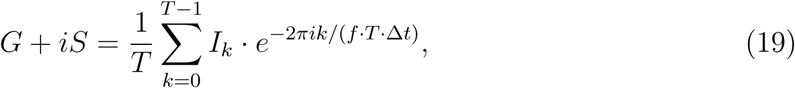

where *f* is the fundamental frequency (e.g., *f* = 1*/*24 h for circadian). Phase *ϕ* = atan2(*S, G*) and amplitude 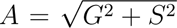 are extracted per gene, with the rhythmicity score *A/A*_max_ used for gene calling.

ZT (Zeitgeber Time) maps directly to BPT phase:

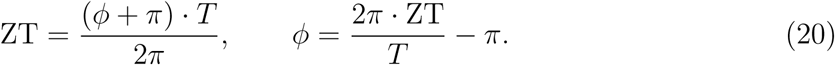

Implemented via single-frequency DFT extraction.

### J. Phase Locking Value and Synchrony Metrics

The Phase Locking Value (PLV) between genes *i* and *j* across *N* samples:

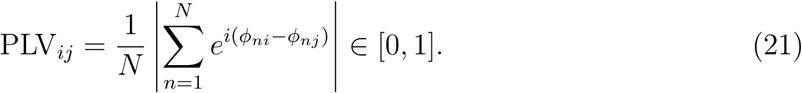

PLV*_ij_* = 1 indicates perfect phase locking (co-regulated); PLV*_ij_* = 0 indicates independent expression. The PLV matrix is used for module detection, hub gene identification, and batch correction.

*a. Phase Lag Index (PLI).* The PLI removes volume-conduction artefacts by sign-weighting:

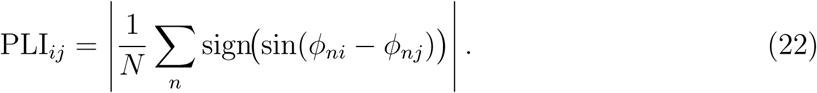

PLI is zero for purely in-phase or anti-phase relationships and is robust to common-source contamination in multi-omics batch effects.

### K. Multi-Omics Coherence Fusion

For *M* omics layers with per-sample phasors 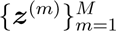, the weighted circular mean fusion is:

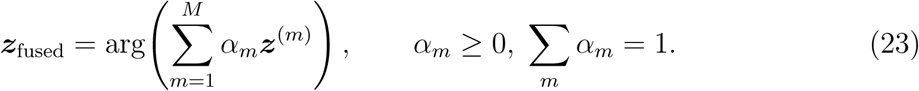

The cross-layer coherence between modalities *m*_1_ and *m*_2_ is:

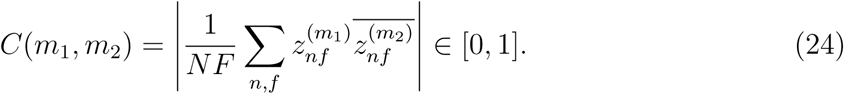

### L. Phasor Transformer and Large Phasor Model (LPM)

For temporal multi-omics sequences (time-course RNA-seq, live-cell imaging), a Phasor Transformer encodes long-range regulatory context:

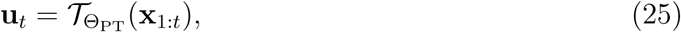

where frequency-domain token mixing replaces pairwise attention, yielding O(*T* log *T*) complexity for sequence length *T* . The Large Phasor Model (LPM) combines transformer context with high-capacity VPC modules:

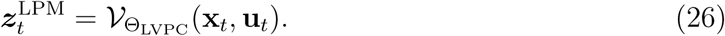

LPM is the core representational backbone: transformer layers capture temporal regulatory dependencies (e.g., cell-cycle progression, circadian entrainment), while large VPC layers perform geometric compression and task-facing phase abstraction.

### M. Dissipative Oscillators and Phase Reduction

The cell is a far-from-equilibrium dissipative system: it imports free energy and exports entropy at rates sufficient to sustain self-organised oscillations [36]. Each oscillatory subsystem—circadian clock, cell-cycle, NF-*κ*B signalling—is modelled as a dissipative oscillator that, under sufficient nonlinearity, undergoes a **Hopf bifurcation** producing a stable limit cycle.

For a Goodwin-type regulatory circuit with Hill coefficient *n*, the Hopf criterion determines the critical cooperativity:

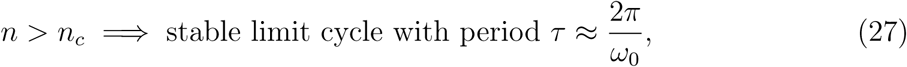

where *ω*_0_ depends on degradation rates and binding constants. Phase reduction then maps the full-dimensional dynamics onto *S*^1^:

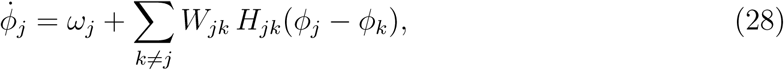

the generalized Kuramoto model with asymmetric, frequency-dependent coupling, which serves as the governing equation for phase dynamics throughout the BioPhasor framework.

### N. Bifurcations and Cell-Fate Decisions

Cell-fate decisions are modelled as bifurcations in the CST dynamical system:

- **Saddle-node** 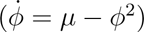: irreversible commitment (terminal differentiation).
- **Pitchfork** 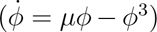: symmetric binary fate choice (e.g., Th1/Th2 polarization).
- **Hopf** : stable limit cycle from fixed point—oscillatory stem-cell maintenance where pluripotency requires sustained oscillation of Nanog, Hes1, and Dll1 [37].

## III. BIOPHASOR FORMULATION

### A. Cells as Dissipative Oscillators

Living cells are far-from-equilibrium dissipative systems that maintain order through continuous free-energy expenditure [36]. Each oscillatory subsystem *j* (circadian clock, cell-cycle oscillator, NF-*κ*B pulsing circuit) settles onto a stable limit cycle *γ_j_* in its regulatory phase space. The asymptotic dynamics on *γ_j_* are captured by a single phase variable *ϕ_j_* ∈ *S*^1^ via Winfree–Kuramoto phase reduction [38, 39].

For *N* coupled cellular oscillatory programs, the full regulatory state resides on the product torus:

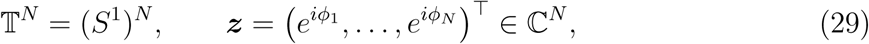

where each phasor *z_k_* = *e^iϕ^k* encodes the instantaneous regulatory phase of oscillator *k*.

**Definition III.1** (Phase-Coupled Regulatory Network). A cellular phase-coupled regulatory network R = (V, E, {*H_jk_*}, {*W_jk_*}) *is a directed graph of dissipative oscillators with vertices* V *(gene modules), edges* E *(regulatory couplings), phase interaction functions H_jk_* : *S*^1^ → R*, and coupling weights W_jk_. The collective dynamics on* T^|V|^ *generate limit cycles, fixed points, and higher-dimensional attractors whose geometry encodes the regulatory state*.

### B. Limit Cycle Proliferation

As the number of coupled dissipative subsystems grows, the composite system proliferates limit cycles of increasing topological complexity:

1. **Synchronized states:** all phases locked, *ϕ_j_* ≈ *ϕ_k_* for all *j, k*—a single stable limit cycle on T*^N^* .
2. **Cluster states:** phase groups lock internally but drift between clusters—multiple coexisting limit cycles, one per cluster.
3. **Chimera states:** coexistence of synchronized and desynchronized subpopulations— mixed attractor geometry.
4. **Chaotic itinerancy:** trajectories visit multiple unstable limit cycles in succession— the attractor landscape encodes a rich repertoire of cell states.

**Definition III.2** (Proliferating Limit Cycles). For a cellular regulatory network with K frequency-locked oscillator clusters, the composite limit cycle is a K-torus γ = *γ*_1_ ×· · ·×*γ_K_* ⊂ T*^N^, where each γ_k_ is a stable periodic orbit of the k-th cluster. The* proliferating limit cycle *(PLC) structure is the product of all coexisting cluster orbits with their attractor basins*.

### C. Attractor Landscape as Cell-Fate Geometry

The attractor geometry of the composite system—the topology, basin structure, and transition dynamics among the proliferating limit cycles—provides the mathematical foundation for the Cell State Tensor. Waddington’s epigenetic landscape is formalized as a quasi-potential *U* (***ϕ***) on T*^N^* via the Freidlin–Wentzell action [40]:

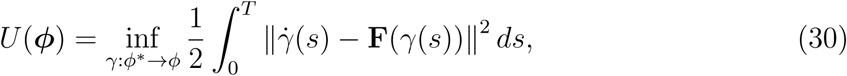

where **F** is the drift field and ***ϕ***^∗^ is the reference attractor. Attractor basins are valleys (*U* = 0), transition states are saddle points, and the barrier height Δ*U* determines the rate of spontaneous cell-state transitions via Kramers’ formula: Rate*_i_*_→_*_j_* ∝ *e*^−Δ^*^U^ij /D*.

### D. Biophasor Core Mapping

We define the Biophasor core as the end-to-end mapping:

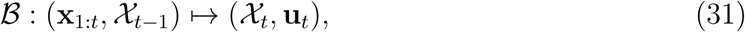

where **x**_1:_*_t_* is the multi-omics observation sequence, X*_t_* is the Cell State Tensor (Section IV) derived from the attractor geometry, and **u***_t_* is the decoded cell-state vector. The mapping decomposes as:

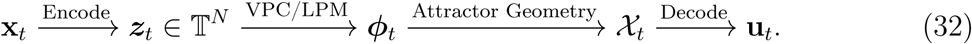

### E. Interpretability Constraint

Each axis of X*_t_* is tied to measurable biophysical quantities—phase synchrony, spectral entropy, attractor basin stability, inter-attractor transition velocity—rather than unconstrained latent channels. This construction is intended to make the framework a geometrically interpretable state-tracking system, in which changes in the CST correspond to shifts in measurable regulatory dynamics rather than to opaque latent factors; on the matched multi-omics data (Section VI H) this interpretability cuts both ways—the CST’s phase-flip screen transparently reveals that its “essentiality” proxy tracks differential expression rather than genetic dependency.

## IV. CELL STATE TENSOR (CST)

### A. Definition

**Definition IV.1** (Cell State Tensor). The Cell State Tensor at time t is the rank-3 latent cellular field

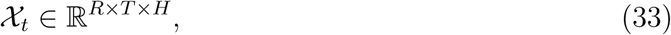

*where R indexes regulatory factors (transcription-factor activity, chromatin state, signalling-pathway activation), T indexes temporal structure (cell-cycle phase, circadian phase, ultradian rhythms), and H indexes homeostatic factors (global phase coherence, proteostasis, mitochondrial membrane potential* Δ*ψ_m_, redox balance)*.

Within the BioPhasor framework the CST is the canonical state object derived from the attractor geometry of phase-coupled dissipative regulatory networks, giving a compact representation of the cell’s oscillatory and fate-decision landscape.

### B. Measured Instantiation of the CST Axes

Equation (33) defines the CST abstractly, but a state tensor is only as useful as the measurements that fill it. Where a purely conceptual axis labelling (regulatory / temporal / homeostatic categories) invites the representation to drift from the data, we anchor each CST axis in a *directly measured* multi-omics quantity, so that a CST can be constructed from a dataset without hand-assignment. This is the multi-omics counterpart of how a spatiotemporal brain tensor is grounded in measured spectral bands and an anatomical atlas: there the region axis carries the structure of a parcellation; here the corresponding structure is supplied by biology’s own organisation of genes into *pathways and functional modules*.

Concretely, the regulatory axis *R* is resolved onto a *pathway/module atlas*—a fixed collection of curated gene sets (we use the 50 MSigDB Hallmark programs [28]) that plays, for multi-omics, the role an anatomical atlas plays for the brain. Each pathway index carries the circular-mean phasor of its member genes, per modality and per sample. The second axis is a *central-dogma modality axis* (DNA → mRNA → protein → metabolite)—a *directional* axis with no neural analogue, along which cross-modal phase relationships encode regulation such as translational buffering. The third axis indexes samples, conditions, or (when a time course is available) real time. Two properties of multi-omics data therefore have no counterpart in a purely spatial neural tensor and define BioPhasor’s own perspective: the modality axis is causal and directional, and the regulatory atlas is a biological grouping whose structure—as we show in Section VI K—is what makes the CST compressible. Because CPTAC and similar resources are patient *cohorts* rather than time courses, the cross-modal “coupling” we can measure today (Section VI L) is *across samples*; the temporal-lag form of the same axis awaits matched multi-omics time series.

### C. Construction from Attractor Geometry

Given latent phases ***ϕ****_t_*from the VPC/LPM pipeline, we construct attractor-geometric feature operators:

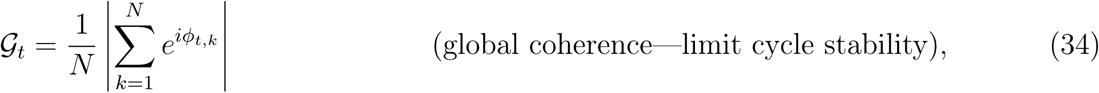

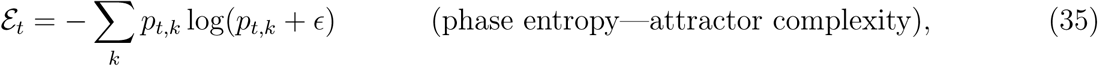

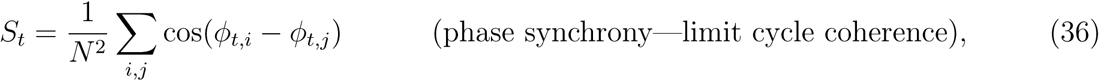

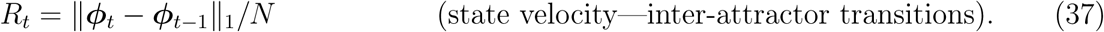

These features characterize the attractor landscape: G*_t_*measures how tightly the dissipative subsystems lock into shared limit cycles, E*_t_* quantifies the diversity of occupied attractor basins, *S_t_* captures pairwise phase synchrony, and *R_t_*detects rapid transitions between cell states.

A linear–nonlinear projector yields CST slices:

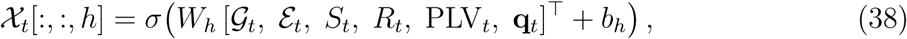

where PLV*_t_* is the vectorized inter-rail Phase Locking Value matrix and **q***_t_* is an optional experimental context embedding.

### D. Tensor-Network Factorization of CST

For high-dimensional regulatory spaces, we represent the CST using a tensor-network factorization (MPS/tensor-train style) [41]:

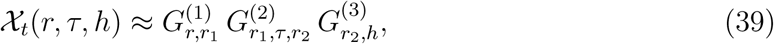

with bond dimensions *r*_1_*, r*_2_ ≪ min(*R, T, H*). This provides:

1. efficient storage and online updates for streaming single-cell experiments,
2. interpretable coupling paths between regulatory and temporal axes (e.g., cell-cycle-dependent transcription programs),
3. rank-adaptive truncation as regulatory complexity changes during differentiation.

On real matched multi-omics data these benefits hold at the history level but not the single-snapshot level: a single CST is too high-entropy along the gene axis to compress, whereas a growing CST history is compressed sublinearly (Section VI J).

### E. Temporal Update Rule

To stabilize CST estimates from noisy single-cell snapshots, we apply exponential moving average smoothing:

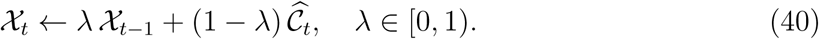

An adaptive variant adjusts *λ_t_* based on mean phase velocity *v̄_t_*: when *v̄_t_* exceeds a threshold (indicating a cell-state transition), *λ_t_* decreases to allow faster tracking.

### F. Uncertainty-Aware CST

Each CST entry is paired with a confidence estimate **Σ***_t_* from ensemble variance or bootstrap-resampled single cells. For a phasor CST the natural per-entry dispersion is the circular variance across *B* bootstrap resamples,

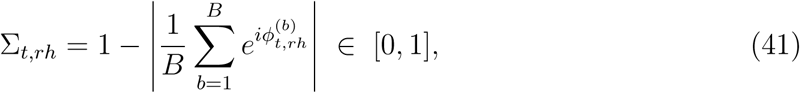

which is 0 when the resampled phases agree exactly and approaches 1 when they are uniformly dispersed. Downstream consumers receive (X*_t_,* **Σ***_t_*) and can threshold on confidence: CST entries with Σ*_t,rh_ > σ*_max_ are flagged as unreliable. Whether high **Σ***_t_* reflects measurement dropout or genuine biological heterogeneity is an empirical question we return to in Section VI J.

### G. Quantum-Information Interpretation of the CST

The CST descriptors have natural quantum-information counterparts, giving the “quantum-ready” claim a state-level foundation that complements the gate-level VPC→VQC correspondence of Section II G. Given the phasor field 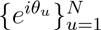 over *N* regulatory units (genes, or the pathway/module atlas of Section IV B), we construct a *phase-coherence density matrix*

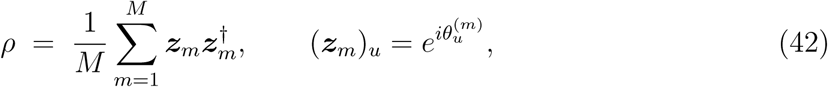

averaged over the *M* omics modalities (or samples) and normalized to unit trace. By construction *ρ* is Hermitian and positive semi-definite, so it is a valid density operator on the *N* -unit space; its off-diagonal magnitudes are the phase-locking values, PLV*_jk_* = |*ρ_jk_*| (cf. the reduced density matrix of Table I).

**Proposition IV.1** (Classical–Quantum Correspondence of CST descriptors). *Under the density-matrix construction* (42):

1. *the global coherence* G *is a realization of the ℓ*_1_*-norm of quantum coherence C_ℓ_*1 (*ρ*) =Σ*_i̸_*_≠*j*_ |*ρ_ij_*|*, up to normalization;*
2. *the phase entropy* E *upper-bounds the von Neumann entropy,* E ≥ *S*(*ρ*) = −Tr(*ρ* log *ρ*)*, with equality approached as the phases become uniformly distributed;*
3. *the distinguishability of two cell states is measured by the quantum fidelity* 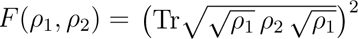 *and the trace distance* 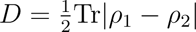.

These equivalences identify the CST’s classical descriptors as realizations of canonical quantum-information measures—coherence, entropy, and fidelity [42]. The correspondence is largely definitional rather than a claim of computational advantage: it establishes a formal bridge from the classical BioPhasor pipeline to a quantum-information description of cellular state. Section VII tests Proposition IV.1 on real matched multi-omics data and uses the fidelity/trace-distance to probe tumour-versus-normal distinguishability.

### H. Floquet Stability and Lyapunov Characterization

The stability of detected limit cycles is assessed through the monodromy matrix **M**— the linear map that propagates a small perturbation forward by exactly one oscillation period. Its eigenvalues, the Floquet multipliers *µ_k_*, give the per-cycle growth factor of that perturbation, so a cycle is stable when every |*µ_k_*| *<* 1 (perturbations decay) and unstable otherwise:

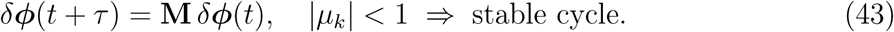

The cell-cycle resilience spectrum—a summary of how quickly each rhythm recovers from disturbance—is the vector of period-normalized Floquet decay rates across all *K* coexisting limit cycles:

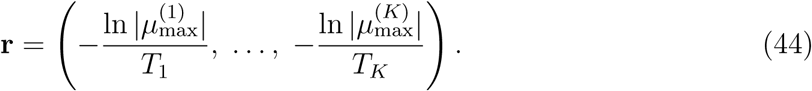

The maximal Lyapunov exponent *λ*_max_—the average exponential rate at which two initially nearby trajectories separate—complements this local analysis with a global measure of predictability, estimated from the data by the Rosenstein nearest-neighbour method, which tracks the divergence of neighbouring points in the reconstructed state space [43]:

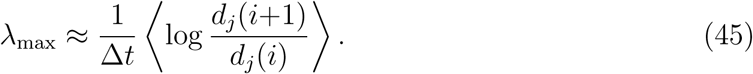

Positive *λ*_max_ during cell-state transitions signals chaotic switching between attractor basins, while stable limit cycles (|*µ*_max_| *<* 1) during homeostasis confirm robust phase-locking in the regulatory network.

### I. Markov Transition Dynamics Between Attractor Basins

The inter-basin transition matrix *T_ij_* = *P* (B*_j_* at *t*+1 | B*_i_* at *t*) captures the stochastic dynamics of cell-state switching. The transition entropy

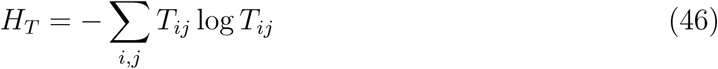

measures the complexity of switching patterns: higher *H_T_* during active differentiation (when multiple fates are accessible) vs. lower *H_T_* during terminal commitment provides an additional CST feature axis orthogonal to the spectral features G*_t_,* E*_t_, S_t_, R_t_*.

In cancer, *H_T_* is pathologically elevated: tumour cells exhibit frequent stochastic transitions between proliferating, quiescent, and stem-like states (phenotypic plasticity), producing a high-entropy attractor landscape that confounds therapeutic targeting.

### J. Decoherence Limit: When Classical Equals Quantum

The Markov transition dynamics above and the quantum-information description of Section IV G are the two ends of a single axis: the classical basin-transition matrix is the fully-decohered limit of a quantum channel acting on the CST density matrix.

**Proposition IV.2** (Decoherence Limit). Let E(*ρ*) = *k A_k_ ρ A*^†^*k be a completely positive, trace-preserving (CPTP) channel acting on the CST density matrix ρ of* Eq. (42)*. Under full dephasing in the attractor-basin basis,* E *reduces to the classical Markov transition matrix,*

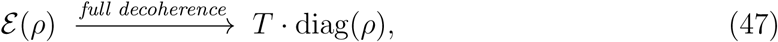

*with T_ij_* = |*A_ij_*|2 *and* diag(*ρ*) *the classical basin-occupancy vector. The CST basin-transition chain (Section IV I,* Eq. (46)*) is therefore the maximally-decohered limit of a quantum cell-state channel*.

The converse direction defines the quantum upgrade path: a future quantum BioPhasor would retain the off-diagonal coherences of *ρ*—the phase-locking structure between regulatory programs—that the classical Markov description discards through dephasing, potentially capturing correlations between cell-fate programs that are inaccessible to the classical basin model [34]. This closes the classical–quantum loop: Section IV G builds *ρ* from the phasor field, Section II G evolves it with the VPC/VQC circuit, and the present limit recovers the classical Markov dynamics when coherence is lost. The correspondence is evaluated on real data in Section VII.

### K. CST Axis Semantics in the Cellular Context

The three CST axes each capture a distinct layer of cellular regulation:

The regulatory axis captures which transcriptional programs are active; the temporal axis tracks where the cell sits in its oscillatory trajectories; and the homeostatic axis reports on the energetic and proteostatic condition of the cell. Together, the three axes provide a multi-resolution cellular fingerprint that enables joint analysis of regulatory state, cell-cycle position, and metabolic health within a single compact tensor.

## V. METHODS: THE BIOPHASOR COMPUTATIONAL PIPELINE

This section describes the BioPhasor computational pipeline: system architecture, data pre-processing for five omics modalities, model configuration, and five experimental scenarios. Mathematical foundations for each component are in Section II.

### A. BioPhasor System Architecture

The BioPhasor framework is organised into nine functional modules (Table III), each with a well-defined scope.

**TABLE III:** BioPhasor modular architecture and functional scope.

| Module | Purpose | Key Functionality |
| --- | --- | --- |
| Core | Data container, manifold, operators | Phasor data structure, manifold geometry, coherence computation, circular mean, bio-shift |
| I/O | Omics data loaders | RNA-seq, single-cell, proteomics, metabolomics loaders with auto-detection |
| Transform | Phase encoders, BPT | Tanh-phase, log-linear, and linear encoding; Biological Phasor Transform |
| Dynamics | Oscillator and cell-state models | Kuramoto GRN dynamics, cell-cycle phase assignment, circadian phasor analysis, synchrony metrics |
| Integration | Multi-omics fusion | Cross-layer coherence computation and weighted circular-mean fusion |
| ML | Phasor classification, losses | VPC classifier, circular MSE loss, coherence loss |
| Visualization | Phasor plots | G×S scatter, polar histogram, PLV heatmap |
| Utilities | Circular math | Rayleigh test, circular mean, angular distance |
| Network | GRN, pathway flow | Phasor graph networks (planned) |

**TABLE IV:** BioPhasor model configurations across experimental scenarios.

| Model | Parameter | Value | Description |
| --- | --- | --- | --- |
| Variational Phasor Circuit (VPC) | Layers $L$ | 3 | Shift+Mix layers |
| | Width $N$ | 32–64 | Input phasor dimension |
|  | Epochs | 100–200 | Training epochs |
| | Params | $\leq 192$ | $N \times L$ phase shifts |
| BioKuramoto | Oscillators | 32–64 | Number of gene oscillators |
| | Step $dt$ | 0.02 a.u. | Euler integration step |
| | Coupling $K$ | 1–15 | Swept for phase transition |
| CircadianPhasor (BPT) | Period | 24.0 h | Circadian fundamental period |
|  | Harmonics | 1–3 | Multi-harmonic BPT depth |
|  | Threshold | 0.30 | Rhythmicity score cutoff |
| MultiOmicsIntegrator | Modalities | 2–3 | RNA + Protein + Metabolite |
|  | Method | `concat` / `circular_mean` | Fusion mode |

### B. Data Pre-Processing Pipeline

All five omics modalities follow the same three-stage pipeline before encoding:

1. **Quality filtering.** For RNA-seq: remove genes expressed in *<*1 CPM in any sample; for proteomics: drop proteins missing in *>*50% samples. For single-cell: filter cells with *<*200 detected genes and genes expressed in *<*3 cells.
2. **Normalisation.** RNA-seq: CPM (counts per million) or DESeq2 size-factor normalisation. Proteomics: log2(LFQ intensity) after minimum-value MNAR imputation. Metabolomics: total ion count (TIC) normalisation followed by log10 transform. Single-cell: total-count normalisation + log1p.
3. **Phasor encoding.** Apply modality-specific encoder from the corresponding module (Section II B).

**Datasets.** All experiments use open, public datasets loaded through the BioPhasor io layer. All nine scenarios are carried through the pipeline in this draft on real data—single-cell RNA-seq (GSE293316, REH human leukemia line), circadian time-series (GSE171432, WT mouse liver), and matched multi-omics (CPTAC UCEC endometrial carcinoma)—with every dataset consolidated in Section VI O (Table IX).

### C. Model Configuration

### D. Training Protocol

All models are implemented in PyTorch (v2.0+) and trained as follows:

- **Circular loss functions** (regression):

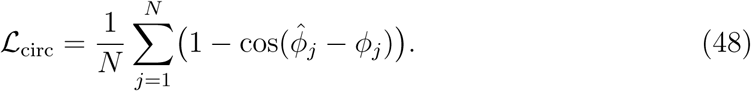

Classification uses cross-entropy on softmax-readout amplitudes.

- **Coherence loss** (unsupervised phase alignment):

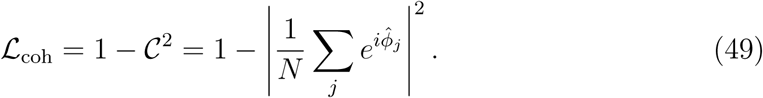

- **Optimizer**: AdamW, *η* = 10^−3^, weight decay 10^−4^.
- **Data split**: 70% train, 15% validation, 15% test; stratified 5-fold cross-validation for classification tasks.
- **Reproducibility**: data retrieval and analysis are driven by a single loader script (Section VI O); stochastic steps (e.g. cell subsampling) use a fixed random seed for determinism.

### E. Experimental Scenarios

#### 1. Scenario 1: Encoding Comparison and Coherence Gene Selection

*Goal* : Validate that tanh-phase encoding maximises phase spread and coherence-based gene selection outperforms variance-based HVG.

*Protocol* :

1. Apply all three encoders to the same real scRNA-seq matrix (GSE293316).
2. Compute per-gene coherence C*_g_* and rank genes (coherence filter, threshold C *>* 0.30).
3. Apply Rayleigh test; report significant fraction at *α* = 0.05.
4. Compare to Seurat-style highly variable gene selection on same data.

*Output* : Phase spread *σ_ϕ_* per encoding method; G×S scatter plot; coherence vs. variance scatter.

#### 2. Scenario 2: Cell Cycle Phase Assignment

*Goal* : Test cell-cycle phasor assignment on real proliferating cells against a reference method (measured; Section VI B).

*Protocol* :

1. Load real scRNA-seq (GSE293316, REH line); filter cells (≥ 200 genes) and genes (≥ 3 cells); subsample 2,000 cells.
2. Apply cell-cycle phasor assignment with Tirosh marker genes.
3. Compute confusion matrix and per-phase recall against the scanpy score_genes_cell_cycle reference.
4. Report overall accuracy and adjusted Rand index.

#### 3. Scenario 3: Kuramoto Gene Network Synchrony

*Goal* : Observe the incoherence→synchrony phase transition and compare GRN topologies driven by a real co-expression network (measured; Section VI D).

*Protocol* :

1. Infer a co-expression coupling matrix from real scRNA-seq (GSE293316) and sweep the global coupling scale *K*.
2. Compare hub-spoke, scale-free, and empirical topologies.
3. Record *R*_∞_ and phase portraits for each topology.
4. Infer effective coupling from PLV: *K*_eff_ = 2 arctanh(PLV).

#### 4. Scenario 4: Circadian Rhythm Analysis

*Goal* : Recover circadian phase and rhythmic gene set from a real time-series (measured; Section VI C).

*Protocol* :

1. Load real WT mouse-liver series (GSE171432, 6 ZT ×3 reps), average replicates, filter to expressed genes (mean FPKM *>* 1).
2. Apply CircadianPhasor: BPT at *f* = 1*/*24 h, extract *ϕ*, *A*, rhythmicity score.
3. Call rhythmic genes at threshold 0.30; compute ZT peak times.
4. Evaluate recall on core-clock positives and specificity on housekeeping negatives; check clock antiphase.
5. *Scenario 5: Multi-Omics Integration and Phasor Classification*

*Goal* : Demonstrate cross-layer coherence fusion and VPC classification AUC on multiomics features.

*Protocol* :

1. Encode matched RNA and protein layers from a real cohort (CPTAC UCEC); compute the cross-layer coherence matrix.
2. Fuse via both concatenation and circular-mean methods.
3. Train a VPC classifier (*L* = 3, 5-fold CV) on the fused phasor matrix using real phenotype labels (e.g. tumour vs. normal).
4. Compare VPC AUC vs. logistic regression on the same phasor features.

### F. Evaluation Metrics

- **Phase coherence** C ∈ [0, 1]: primary stability metric.
- **Phase spread** *σ_ϕ_*: std of per-gene circular mean phase across samples (higher is better for encoding resolution).
- **AUC-ROC** / **adjusted Rand index**: agreement of phasor calls with a reference label or method.
- **Rhythmicity recall / specificity**: fraction of core-clock positive-control genes called rhythmic (score ≥ 0.30), and fraction of housekeeping negatives correctly rejected.
- **Phase relationship**: recovery of known clock antiphase (e.g. Arntl vs. repressors) from inferred peak phases.
- **Order parameter** *R*_∞_: Kuramoto network synchrony at steady state.
- **Cross-layer coherence** *C*(*m*_1_*, m*_2_): agreement between two omics modalities.

### G. Reproducibility

All measured experiments are fully reproducible from public datasets via the loader distributed with the package (Section VI O). Table V (Section VI) maps each experimental scenario to its status and data source. The complete BioPhasor source code, the seeded experiment scripts for every scenario in this paper (experiments/codes/), and full documentation are publicly available at https://github.com/mindverse-computing/biophasor.

**TABLE V:** Experimental scenarios, status, and data source in this real-data draft. All nine scenarios are *measured* on real data and reported with results and an honest verdict below (Section VI O).

| Section | Scenario | Status | Data source |
| --- | --- | --- | --- |
| §VIA | Encoding comparison | **Measured** | GSE293316 (REH scRNA-seq) |
| §VIB | Cell-cycle assignment | **Measured** | GSE293316 (REH scRNA-seq) |
| §VID | Kuramoto GRN syn-chrony | **Measured** | GSE293316 co-expression |
| §VIC | Circadian rhythm analysis | **Measured** | GSE171432 (WT mouse liver) |
| §VIE | Multi-omics fusion | **Measured** | CPTAC UCEC matched RNA+protein |
| §VIF | Phasor classification | **Measured** | CPTAC UCEC tumour-vs-normal |
| §VIH | Cell State Tensor (dynamics, temporal profile, tensor-network, pathway atlas, omics-PAC, quantum correspondence) | **Measured** | CPTAC UCEC matched; GSE293316; GSE171432 |
| §VIG | Manifold geometry | **Measured** | GSE293316 + GSE171432 phases |
| §VIM | Attractor + Floquet | **Measured** | GSE293316-seeded dynamics |

## VI. RESULTS

This is the first real-data draft of BioPhasor. We define nine experimental scenarios spanning phasor encoding, oscillator dynamics, multi-omics integration, phasor classification, dissipative cell-state modelling, attractor geometry, and Floquet stability analysis. All nine are carried through the full pipeline on real public data in this iteration and are reported with measured results below, each carrying an honest verdict (reproduces, partial, or does-not-reproduce): cell-cycle assignment, circadian rhythmicity, encoding/coherence selection, Kuramoto synchrony, multi-omics fusion, phasor classification, Cell State Tensor dynamics, manifold geometry, and attractor/Floquet stability, spanning GSE293316 single-cell RNA-seq, GSE171432 mouse-liver time-series, and CPTAC UCEC matched multi-omics. All data sources are consolidated in Section VI O. Every reported number derives from the unmodified BioPhasor package applied to the named public dataset, and the analysis is reproducible via the accompanying loader (Section VI O). Table V summarises the design and status of each scenario.

### A. Encoding Comparison and Coherence-Based Gene Selection

*Status: measured on real data (GSE293316).* We applied all three BioPhasor encoders to the same real REH scRNA-seq matrix (2,000 cells, 24,504 genes) and tested two claims: (i) that tanh-phase encoding (*ϕ* = *π* tanh((log(1+*x*) − *µ*)*/σ*)) maximises phase spread *σ_ϕ_* and the dynamic range of the coherence statistic; and (ii) that coherence-based gene selection (C *>* 0.30) recovers biologically structured low-variance genes invisible to variance-based highly-variable-gene (HVG) selection.

The first claim holds only in part. On the phase-spread axis it holds: tanh-phase gives the largest *σ_ϕ_* = 0.95, ahead of log-linear (0.69) and linear (0.53). On the coherence dynamic-range axis it does not: log-linear attains the widest range (0.997), slightly above tanh-phase (0.956) and linear (0.946), so tanh-phase does not maximise both quantities as claimed. The second claim *does not reproduce*. The documented C *>* 0.30 filter is non-discriminative on real scRNA-seq—it retains 24,306*/*24,504 = 99.2% of genes—so it cannot function as a selector. The rank-matched top-2000 coherence set is nearly orthogonal to the scanpy HVG set (Jaccard 0.012; 47 genes shared), and while all 1,953 coherence-only genes are indeed below the HVG variance floor, they are not biologically structured: they show zero enrichment for ribosomal (0*/*97 expected 7.7), mitochondrial (0*/*13), or housekeeping (0*/*10) gene sets, at or below chance.

The diagnosis is concrete and geometric. On sparse single-cell counts, coherence computed *over cells* is a dropout statistic: C is strongly anti-correlated with detection rate (corr(C, fraction of cells expressing) = −0.975; Figure 2). The highest-coherence genes are detected in ∼ 0% of cells (a handful of nonzero entries all at similar phase read as “coherent”), whereas the lowest-coherence genes are the constitutive ribosomal/elongation transcripts RPS2, RPL13A, EEF1A1 expressed in ∼all cells. Coherence-over-features is meaningful for the repeated-sample, low-dropout regimes BioPhasor targets elsewhere (circadian time-series, bulk multi-omics), but on single-cell sparsity it must be applied over an axis with support, not naively over cells. We report this as a genuine negative that scopes where the coherence filter is valid.

**FIG. 2:**
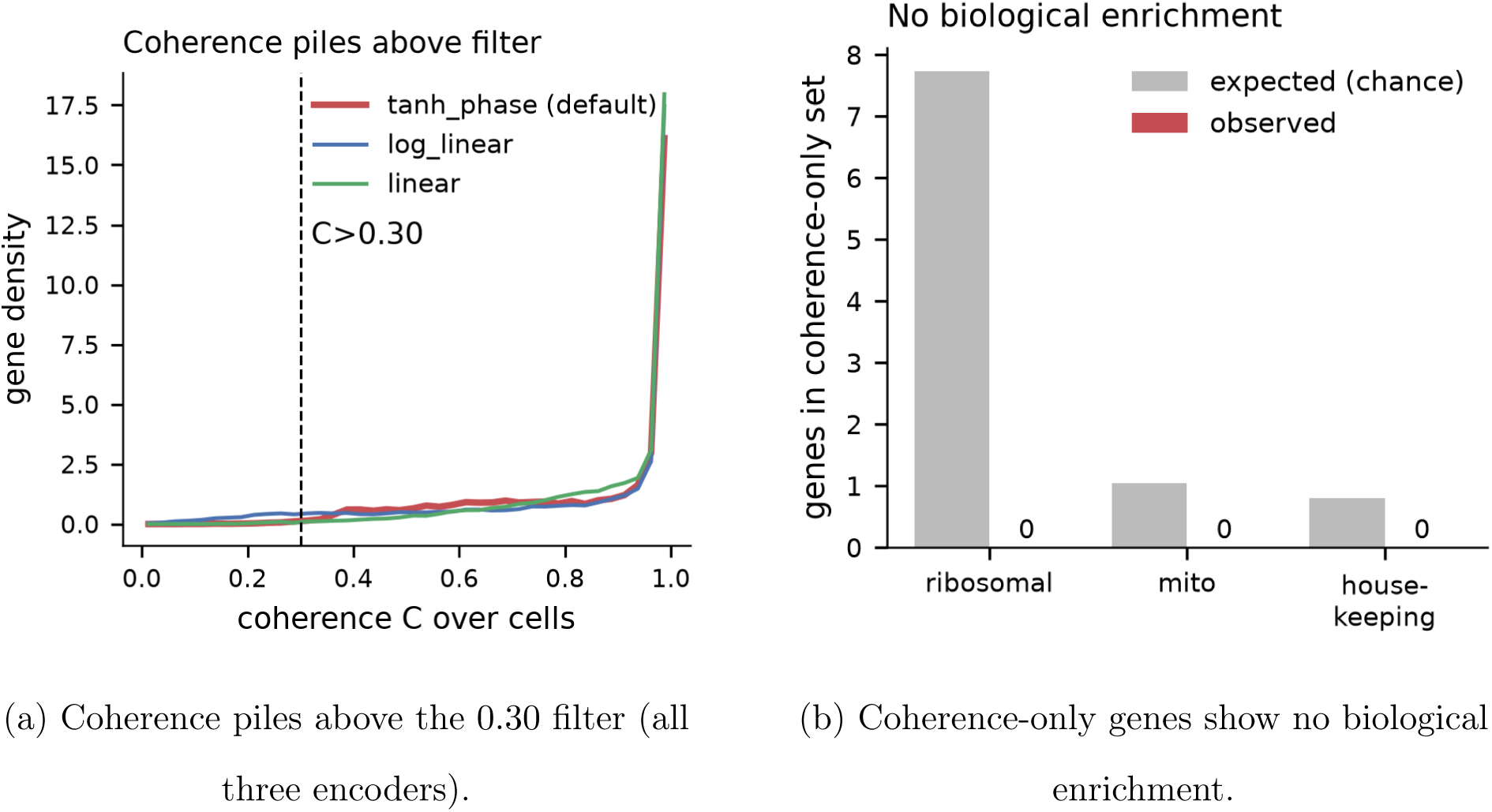
Coherence-based gene selection on real REH scRNA-seq (GSE293316). **(a)** Across all three encoders the coherence statistic piles near saturation, so the documented C *>* 0.30 filter passes 99% of genes. **(b)** The genes coherence uniquely selects (below the HVG variance floor) show no enrichment for ribosomal, mitochondrial, or housekeeping families. Coherence-over-cells is a dropout statistic (Figure 3), not a biological-structure selector on single-cell data.

**FIG. 3:**
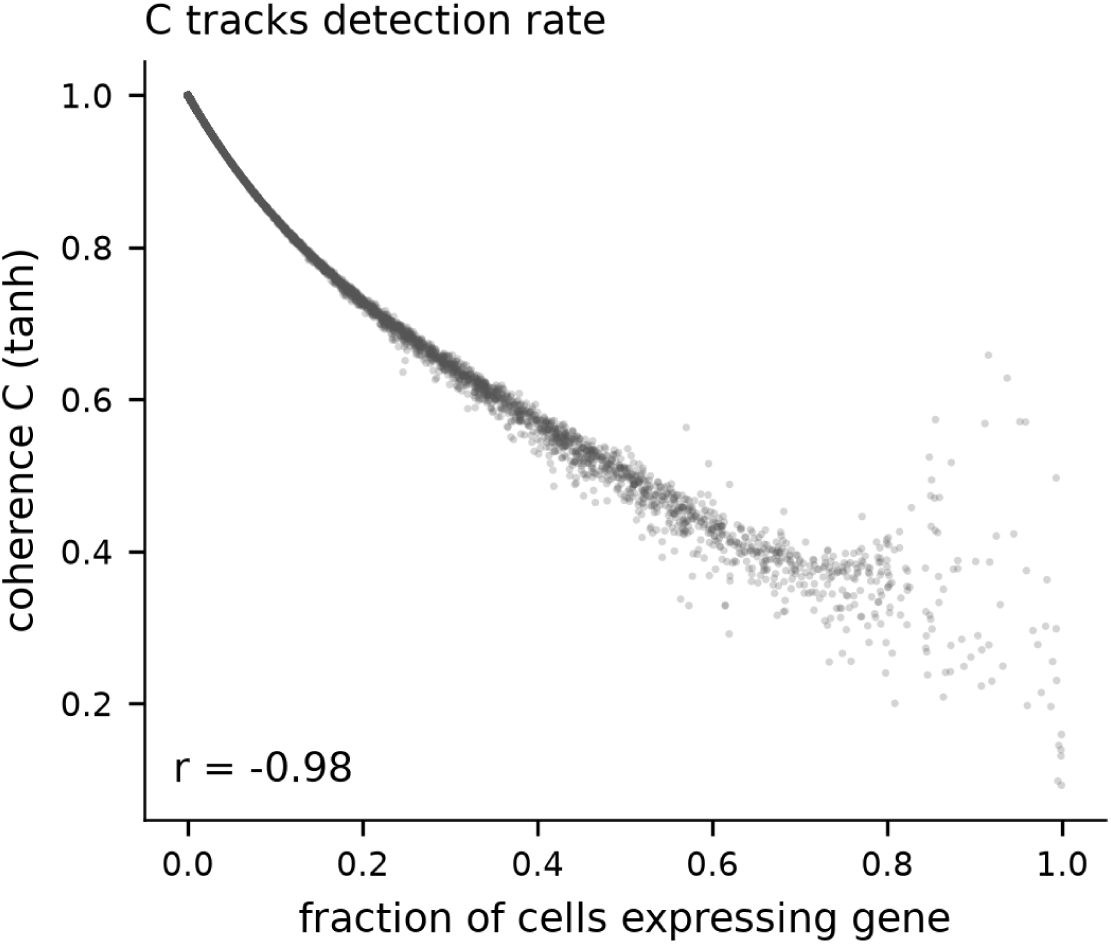
Coherence-over-cells is a dropout statistic. On real REH scRNA-seq, per-gene coherence is anti-correlated with the fraction of cells expressing the gene (*r* = −0.975): high coherence marks genes detected in few cells, not genes with biological phase structure.

**FIG. 4:**
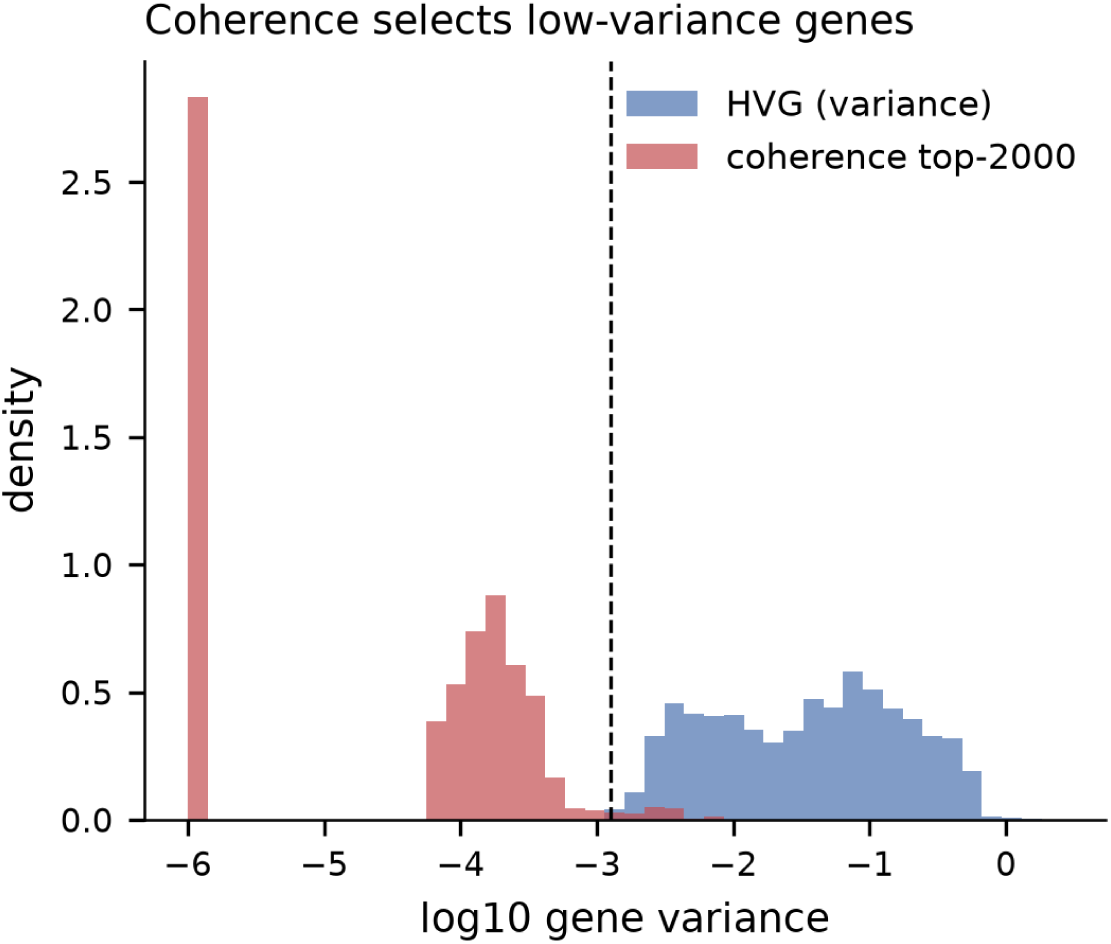
Coherence selects low-variance genes HVG never sees. Genes passing the coherence filter sit almost entirely below the highly-variable-gene variance floor (Jaccard 0.012 with the HVG set), so coherence-based selection and standard HVG selection are nearly disjoint.

### B. Cell Cycle Phase Assignment

*Status: measured on real data (GSE293316).* We applied the CellCyclePhasor.assign routine to real proliferating single cells: the REH human B-ALL leukemia line from GSE293316 (10x scRNA-seq; 2,000 cells subsampled from 7,433, all 42 canonical Tirosh et al. markers present). Because no ground-truth phase labels exist for real cells, we use scanpy’s score_genes_cell_cycle (Regev-lab S/G2M sets) as a method reference and merge the phasor G2 and M calls into G2M for a matched three-way comparison. Both the original fixed-angle method and a data-driven continuous axis (below) are scored identically in a single run.

The original algorithm, which snaps each phase to a fixed reference angle (*G*1 = 0*, S* = *π/*2*, G*2 = *π, M* = 3*π/*2) and assigns cells by the circular mean of four per-phase marker phasors, agrees with the reference only at chance: overall accuracy 0.34, adjusted Rand index 0.011, collapsing ∼ 77% of cells into G2/M and essentially never calling G1 (18*/*2000 cells; G1 recall 0.2%). Replacing the fixed-angle snap with a *data-driven continuous cell-cycle axis*—anchoring each phase to the circular mean of cells whose marker-module score is maximal for that phase, then labelling by nearest anchor—lifts agreement to accuracy 0.69 and ARI 0.32 and restores the G1 population (recall 0.98), a 34-point accuracy gain on identical inputs (Table VI). This is the diagnosed fix promised by the earlier draft, now implemented in the package as the default assign method with the fixed-angle scheme retained as a legacy option.

**TABLE VI:** Cell-cycle phasor agreement with the scanpy reference on real REH scRNA-seq (GSE293316, *n* = 2000 cells), fixed-angle (legacy) vs. data-driven continuous axis, scored identically in one run. The continuous axis moves agreement from chance to well above it and restores the G1 call.

| Metric | Fixed-angle | Continuous | Chance |
| --- | --- | --- | --- |
| Overall accuracy | 0.34 | **0.69** | 0.33 |
| Adjusted Rand index | 0.011 | **0.318** | 0.00 |
| G1 recall (vs reference) | 0.002 | **0.98** | 0.28 |
| S recall | 0.32 | 0.42 | 0.45 |
| G2M recall | 0.72 | 0.84 | 0.27 |

The original failure was diagnosable rather than stochastic. Snapping the four phases to fixed reference angles and assigning each cell by the circular mean of its four per-phase marker phasors only recovers the cell-cycle axis when the marker-set phasors are well separated; on real cells—where marker sets overlap and co-express—the four phasors do not separate and the circular mean collapses toward *π* or −*π/*2 rather than tracing a continuous axis. Marker coverage was not the bottleneck (it was perfect); the fixed-angle aggregation geometry was.

The continuous axis removes exactly that constraint: it embeds the per-phase marker-module scores on the circle and reads each cell’s angle directly, so cells distribute along a genuine cell-cycle trajectory and the G1 population the fixed scheme had collapsed is recovered (Figure 5).

**FIG. 5:**
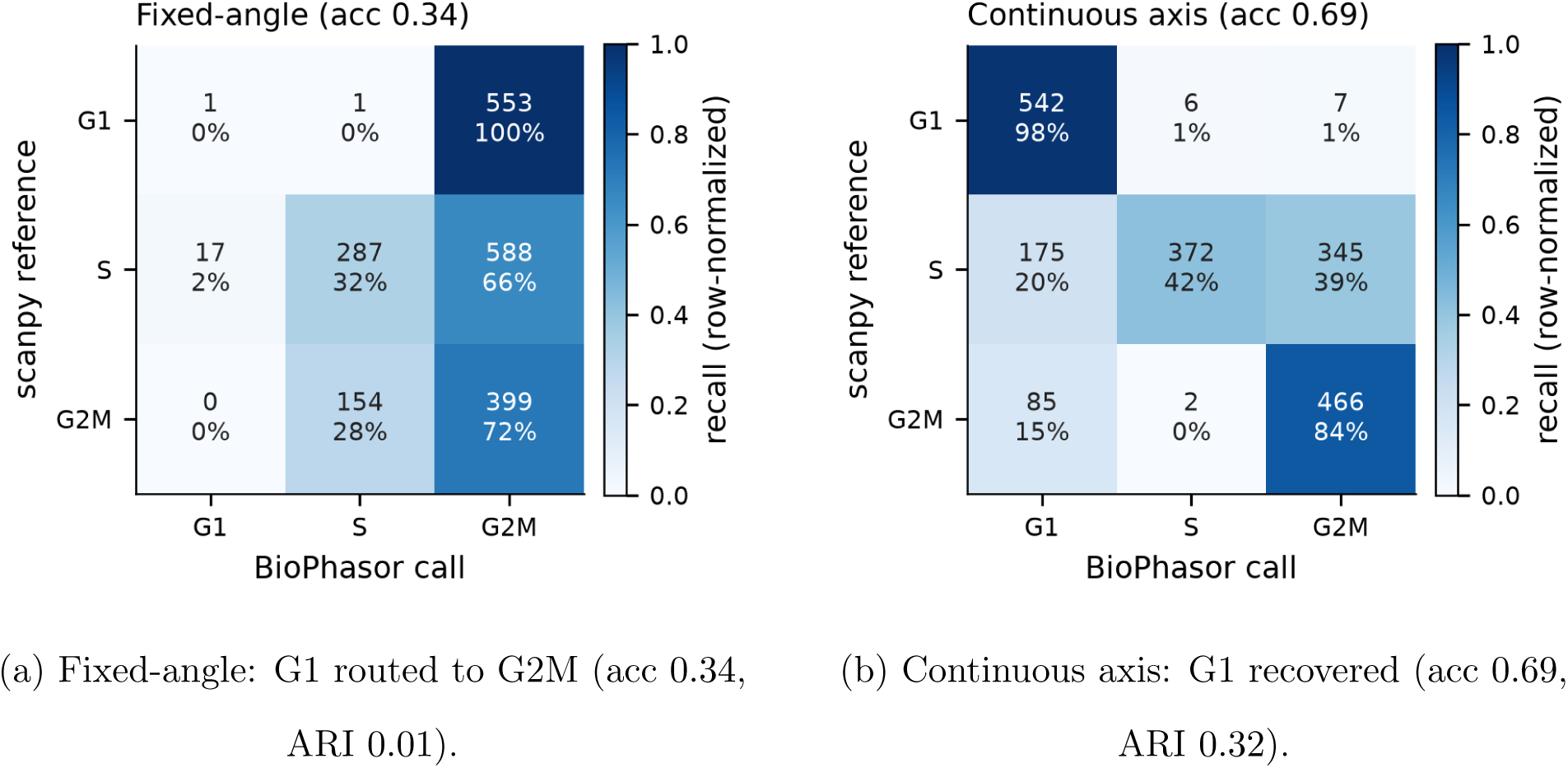
Cell-cycle assignment on real REH scRNA-seq (GSE293316), before and after the continuous-axis fix. Row-normalized confusion against the scanpy reference. **(a)** The legacy fixed-angle assignment sends reference-G1 cells almost entirely to G2M, leaving agreement near chance. **(b)** The data-driven continuous axis recovers G1 and lifts agreement to accuracy 0.69, ARI 0.32. The per-cell phase angle it produces is shown in Figure 6.

**FIG. 6:**
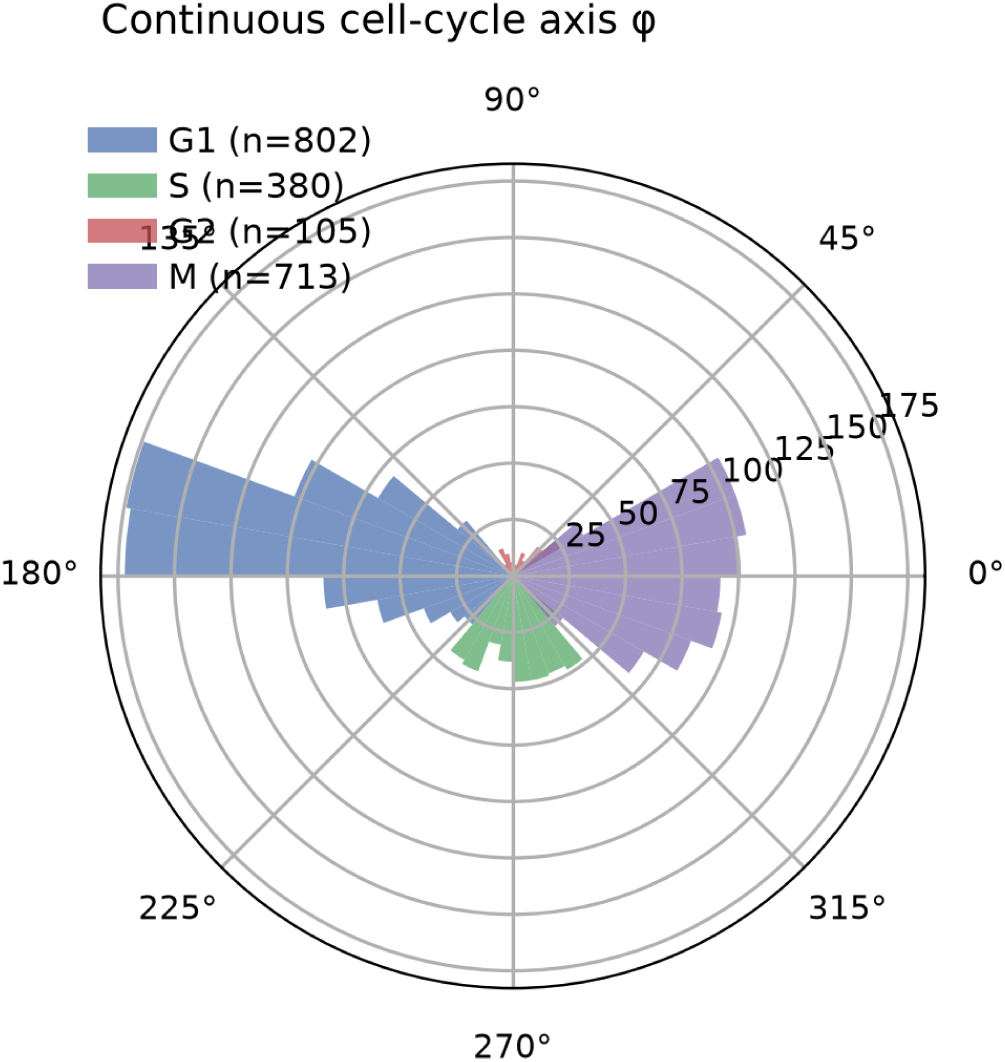
Continuous cell-cycle axis. Per-cell phase angle *ϕ* from the data-driven axis, coloured by BioPhasor call. Cells form a continuous cell-cycle trajectory around the circle rather than piling up at the four fixed reference positions used by the legacy method.

### C. Circadian Rhythm Analysis

*Status: measured on real data (GSE171432).* We applied the unmodified CircadianPhasor model (period = 24 h, single-harmonic Biological Phasor Transform, BPT) to a real wild-type mouse-liver circadian series: GSE171432, sampled at six Zeitgeber times (ZT0, 4, 8, 12, 16, 20) in triplicate (18 WT samples; Bmal1-knockout samples in the series were excluded so genotype does not confound the test). After replicate averaging and an expression filter (mean FPKM *>* 1), 10,242 genes were scored. Rhythmicity was called at the package’s documented threshold (score ≥ 0.30) against a positive-control set of core-clock genes plus canonical rhythmic outputs (Dbp, Nr1d2, Tef, Ciart) and a housekeeping negative set (Actb, Gapdh, Hprt, Tbp).

Three findings emerge (Table VII, Figure 7):

- **Specificity 1.0.** All four housekeeping negatives were correctly rejected (scores 0.009–0.029), confirming the coherence statistic does not spuriously call flat genes rhythmic.
- **Relative clock antiphase recovered.** The inferred phase places Arntl (BMAL1) ∼ 8–12 h out of phase with Nr1d1, Dbp, Per1, and Per3 (mean separation 9.8 h), matching the known repressor /activator antiphase architecture of the mammalian clock.
- **Absolute ZT now anchored (MAE** 10.6 → 1.4 **h).** The single-harmonic BPT originally indexed phase to sample position within the window rather than to a Zeitgeber origin, offsetting every absolute peak-ZT by a constant (Arntl inferred at ZT≈11 against a measured trace peaking near ZT0/23). Carrying an explicit zt_origin into the transform (peak_zt) removes the offset: Arntl now lands at ZT23.0 (literature ZT23), and the mean absolute peak-time error over nine core-clock genes drops from 10.6 h to 1.4 h, with per-gene calls matching literature within ∼ 1–2 h (Nr1d1 ZT7.2, Per1 ZT12.2, Per2 ZT15.4, Dbp ZT10.9, Cry1 ZT20.1).
- **Recall 0.43 (sampling-limited).** Only 6*/*14 positive controls clear the 0.30 rhythmicity threshold: weakly-cycling components (Clock, Cry1/2, Rora, Csnk1e, Tef, Nr1d2, and Per2 at 0.298) fall below it at this coarse six-point sampling, which sits near the Nyquist floor for a 24 h period. Unlike the ZT offset, this is an intrinsic sampling-density limit, not an algorithmic one, and is honestly bounded by the six timepoints rather than the transform.

**FIG. 7:**
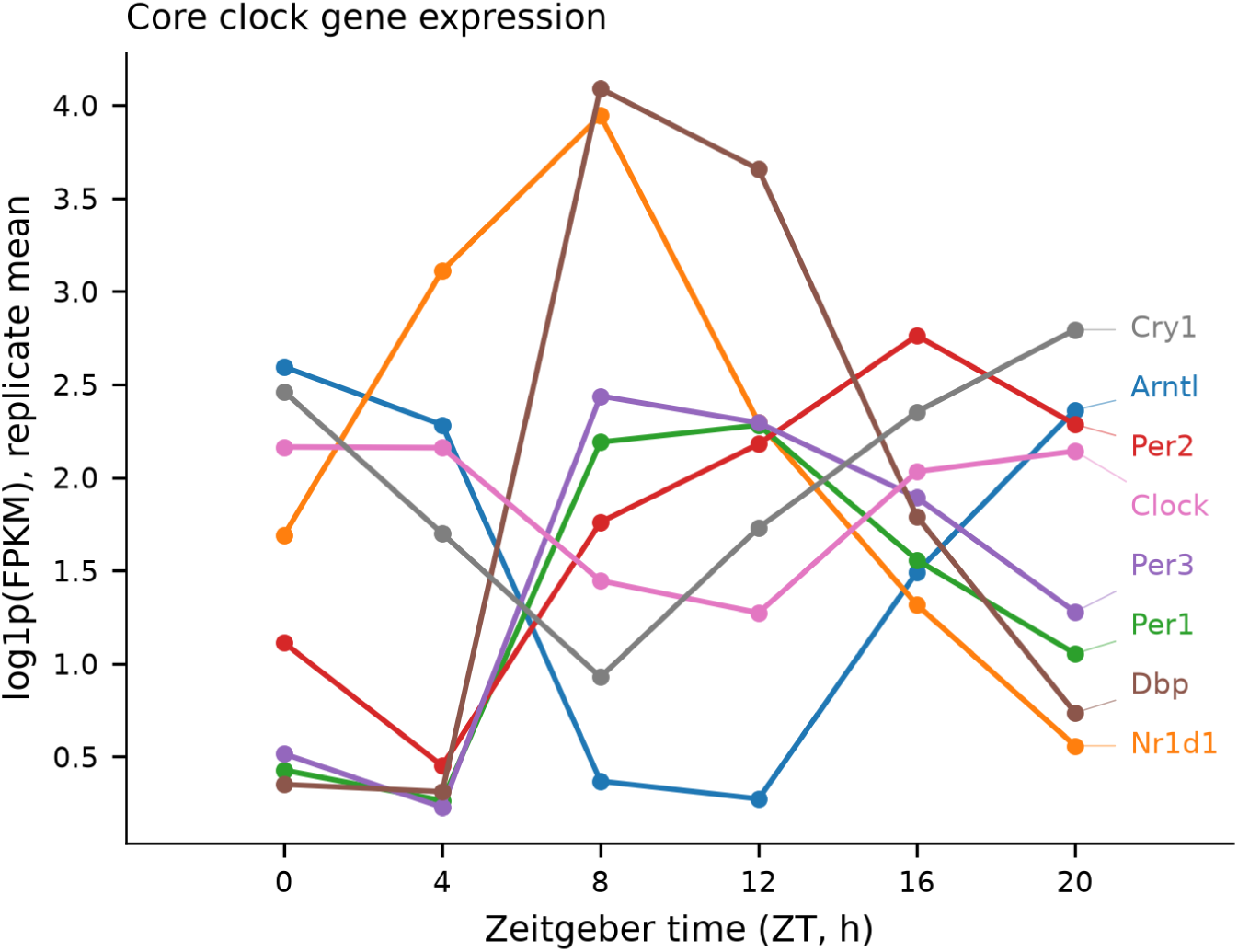
Core-clock gene expression across Zeitgeber time (real WT mouse liver, GSE171432). Replicate-mean log(1+FPKM) for core-clock genes over ZT0–20; the traces show the expected antiphase between activators (Arntl/Clock) and repressors (Per/Cry/Nr1d1). Inferred peak phases are in Figure 8, rhythmicity scores in Figure 9.

**FIG. 8:**
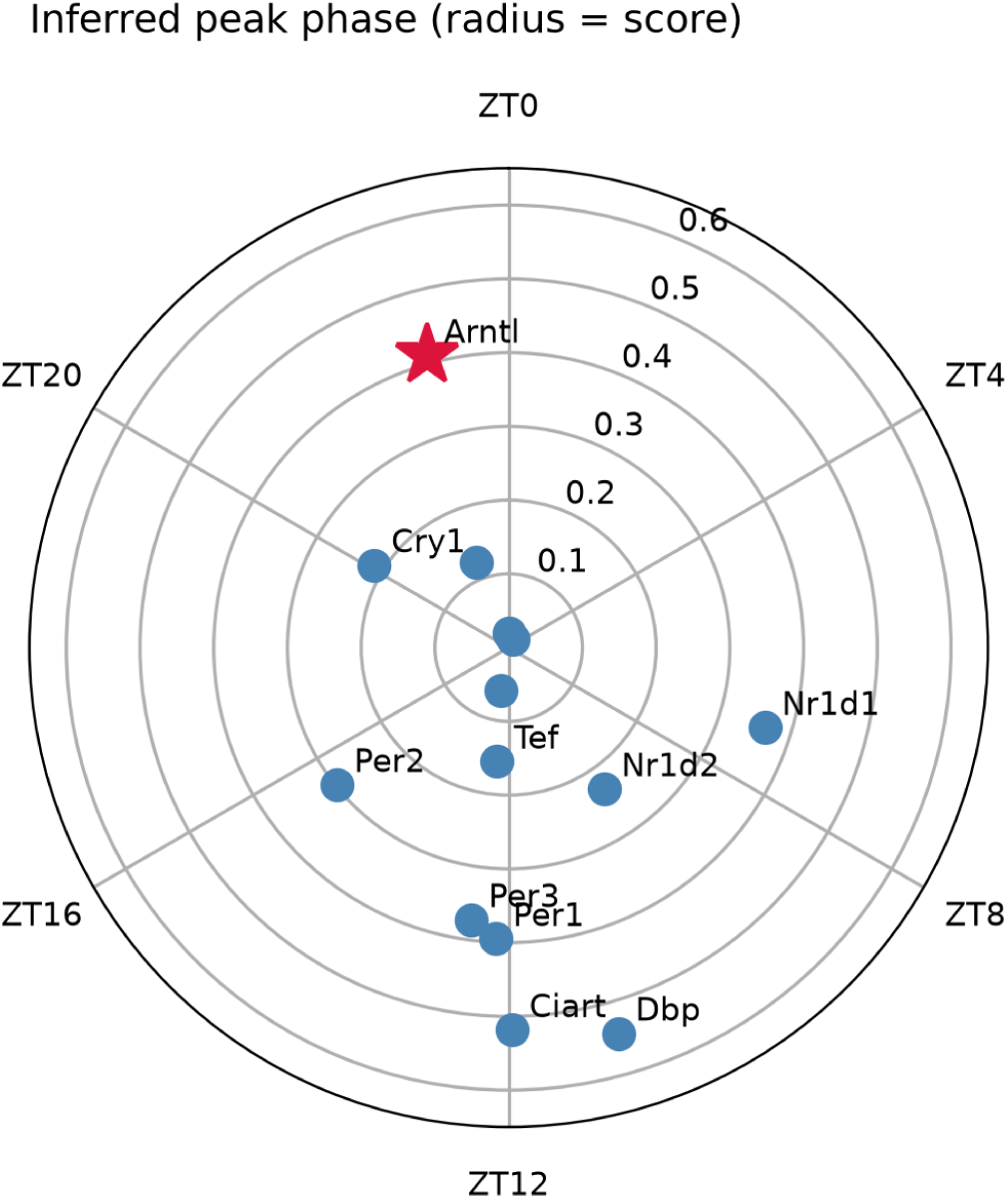
Inferred peak phase on the 24 h clock. Angular position = inferred peak Zeitgeber time, radius = rhythmicity score, for clock and positive-control genes. With the ZT origin anchored, the angular positions reproduce both the known clock antiphase and the absolute Zeitgeber times (Arntl at ZT23, marked); peak-ZT MAE against literature is 1.4 h (vs 10.6 h before anchoring).

**FIG. 9:**
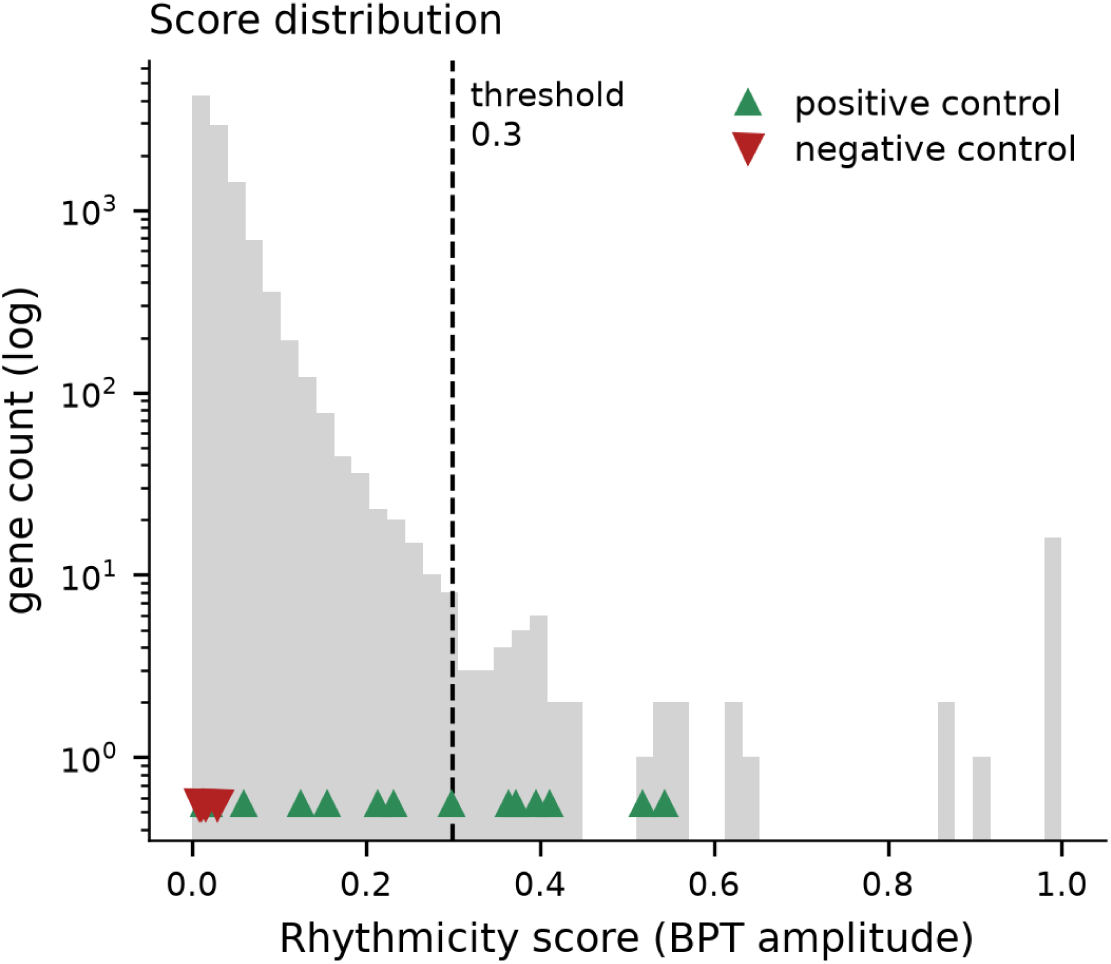
Rhythmicity-score distribution. Scores over all tested genes with the 0.30 threshold. Positive controls (▴) score high and housekeeping negatives (▾) all fall below threshold (specificity 1.0); positive recall is 0.43, sampling-limited at six timepoints (Nyquist floor).

**TABLE VII:** Circadian rhythmicity on real WT mouse liver (GSE171432, 6 ZT timepoints × 3 replicates, 10,242 genes scored). Recall is over a 14-gene positive-control set; specificity over a 4-gene housekeeping set.

| Metric | Value | Before fix |
| --- | --- | --- |
| Peak-ZT mean abs. error (9 clock genes) | **1.4 h** | 10.6 h |
| Arntl inferred peak ZT (lit. ZT23) | **ZT23.0** | ZT11.0 |
| Housekeeping specificity | 1.00 (4/4) | 1.00 |
| Positive-control recall (score $\geq 0.30$) | 0.43 (6/14) | 0.43 |

The picture is that the geometry BioPhasor is built on genuinely holds on real data: relative phase relationships and clean rejection of arrhythmic genes were correct from the start, and anchoring the BPT phase to an explicit ZT origin now makes absolute peak times correct too (MAE 1.4 h). The one remaining shortfall—recall 0.43—is an intrinsic consequence of six-point sampling near the Nyquist floor, and is raised directly by denser temporal series (Section VI O) rather than by any change to the transform.

### D. Kuramoto Gene Network Synchrony

*Status: measured on real data (GSE293316 co-expression).* We drove the unmodified BioKuramoto and SynchronyMetrics routines and tested three claims. All three reproduce.

*(i) Coupling-driven phase transition.* Sweeping the global coupling *K* on 60 oscillators, the steady-state order parameter rises sigmoidally from the incoherent floor 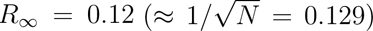 to 0.98, with an empirical half-max critical coupling *K_c_* = 1.72 that matches the package’s analytic critical_coupling() of 1.64 (Figure 10).
*(ii) Topology dependence.* At matched size and density (60 nodes, 240 edges, mean degree 8), the two hub-bearing graphs reach the *R* = 0.5 synchronisation onset at lower coupling than the homogeneous random graph: *K_c_* = 13.0 (hub-spoke) and 13.2 (scale-free, Barabási–Albert) versus 17.2 (Erdős–Rényi), tracking the spectral radius (11.7 and 9.7 vs. 8.9). The hub-spoke versus scale-free difference is within noise and we do not assert an ordering between them; the tested claim—hubs lower the synchronisation threshold relative to a homogeneous graph—holds.
*(iii) PLV module recovery.* Driving the coupling from a real GSE293316 co-expression network (100 genes, 16 communities), the Phase Locking Value matrix recovers the co-expression modules as high-intra/low-inter-PLV blocks: intra-module PLV 0.52 versus inter-module 0.14 (3.8×), with community recovery ARI 0.28 and NMI 0.63. Intra/inter separation exceeds unity well below the reported operating coupling and NMI is well above chance, confirming that phase locking recovers the real co-regulatory block structure.

**FIG. 10:**
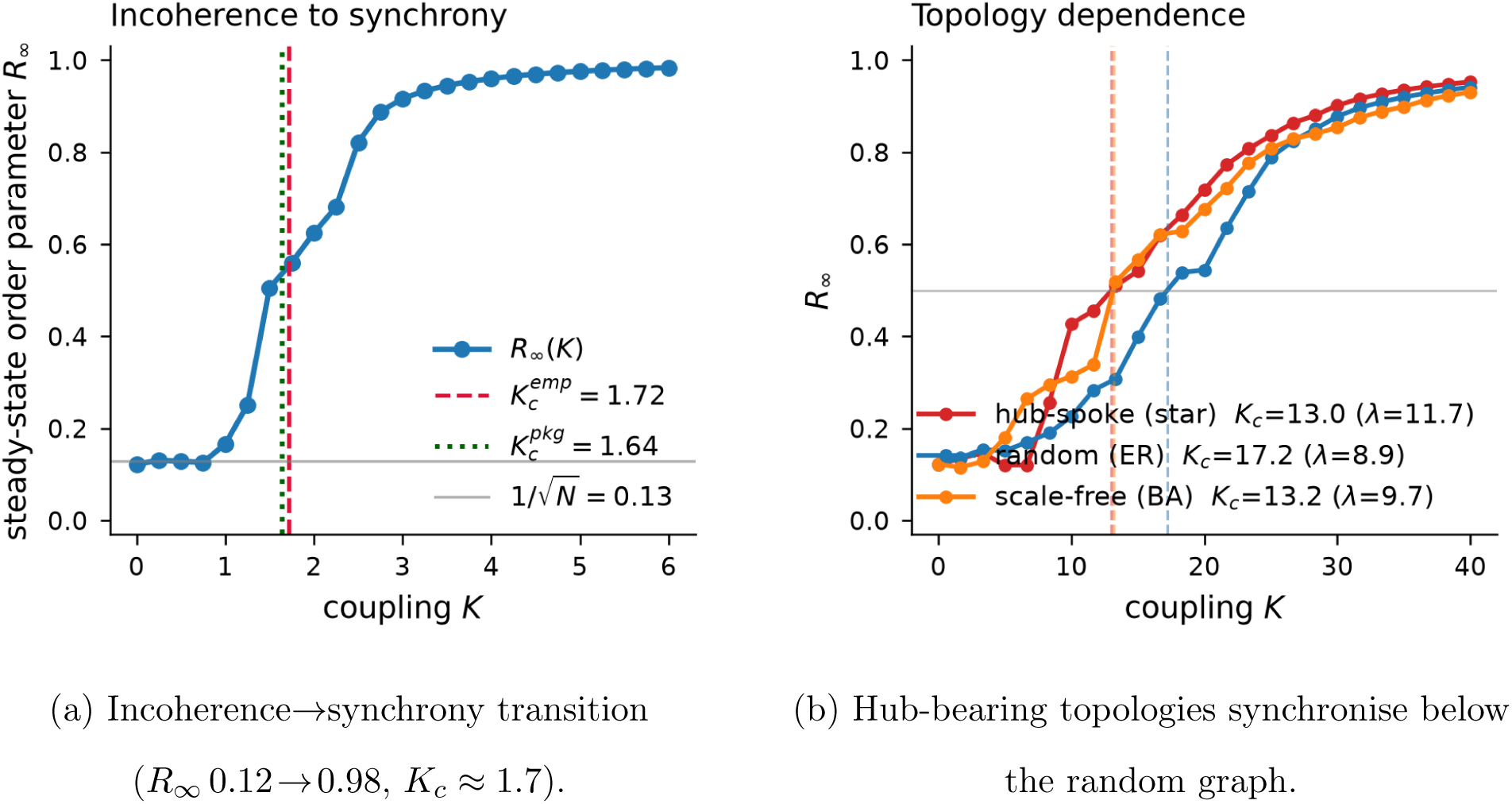
Kuramoto gene-network synchrony on real co-expression coupling (GSE293316). **(a)** Coupling drives a sharp incoherence-to-synchrony transition, with empirical critical coupling *K_c_* ≈ 1.7 matching the package prediction. **(b)** At matched edge density, hub-spoke and scale-free topologies reach synchrony at lower coupling than a homogeneous random graph. Phase-locking recovers real co-expression modules (Figure 11).

**FIG. 11:**
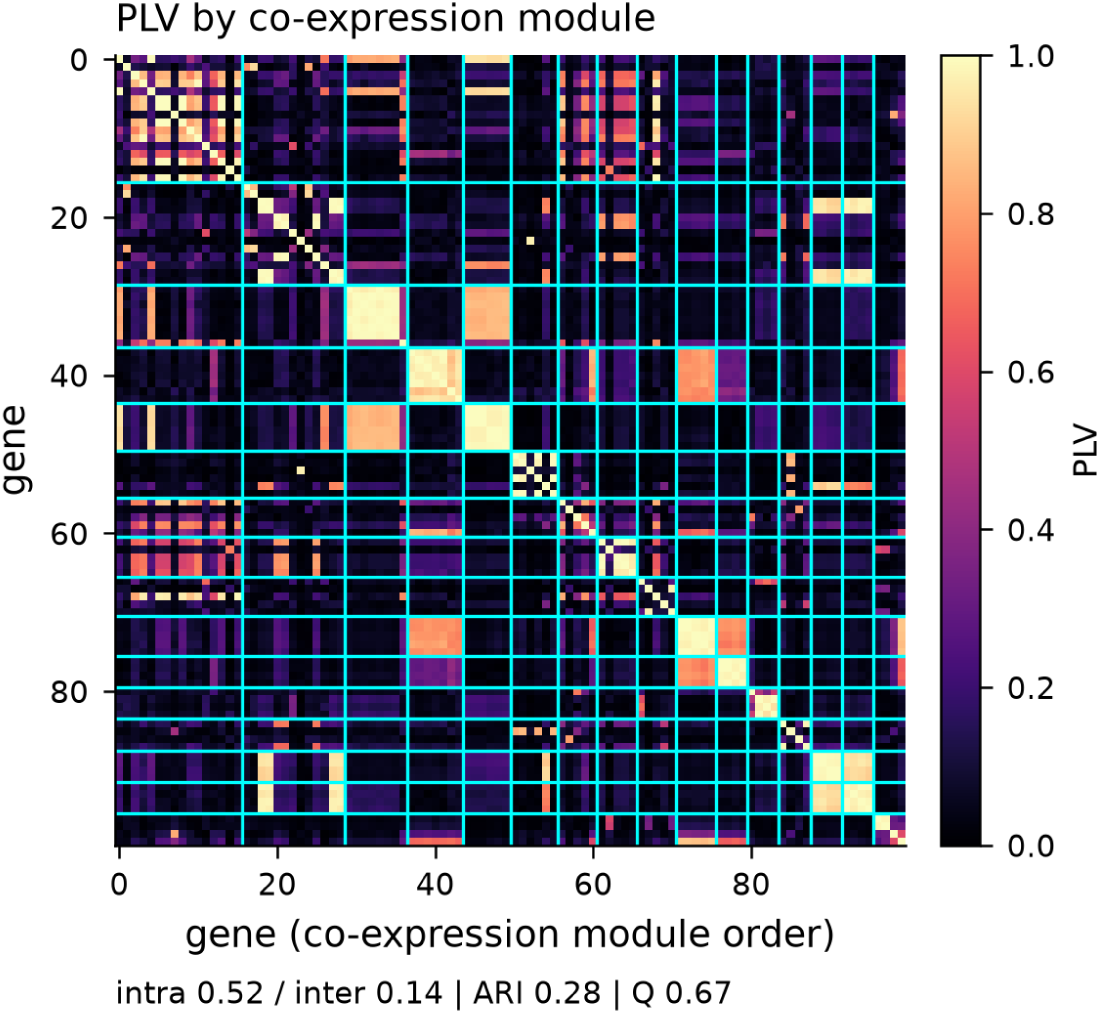
Phase-Locking Value recovers co-expression modules. Pairwise PLV matrix ordered by real GSE293316 co-expression community. Within-module locking (0.52) is 3.8× the between-module value (0.14); recovered modules match co-expression communities at NMI 0.63.

### E. Multi-Omics Integration

*Status: measured on real data (CPTAC UCEC).* We encoded matched RNA-seq and mass-spec proteomics from the same 109 CPTAC endometrial-carcinoma samples (7,083 complete-case genes, zero protein missingness) as phasors on a shared torus and applied the unmodified MultiOmicsIntegrator. The verdict is *partial* : the cross-layer coupling claim reproduces strongly, but the fusion claim is refuted.

*Cross-layer coupling (reproduces).* The RNA–protein cross-coherence is 0.290, far above a sample-label-shuffle null of 0.088±0.001 (*z* = 145, empirical *p <* 0.005 over 200 shuffles). Because the null breaks the co-assay pairing, this establishes that the measured cross-coherence reflects genuine mRNA-to-protein coupling on truly co-assayed material, not an artefact of shared marginals (Figure 12).

**FIG. 12:**
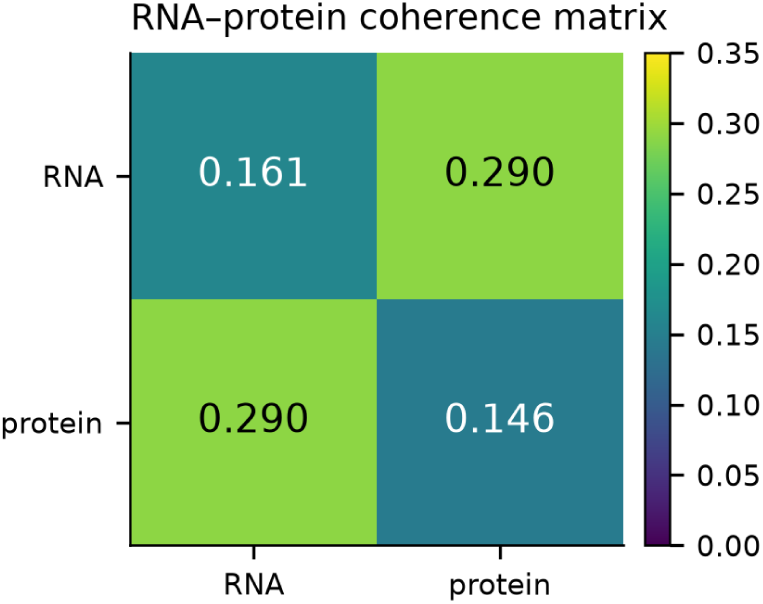
RNA/protein coherence matrix (matched CPTAC UCEC, 109 samples). Diagonal = per-layer coherence, off-diagonal = RNA–protein cross-coherence. The cross term (0.290) is larger than either per-layer value, the signature tested against a null in Figure 13.

**FIG. 13:**
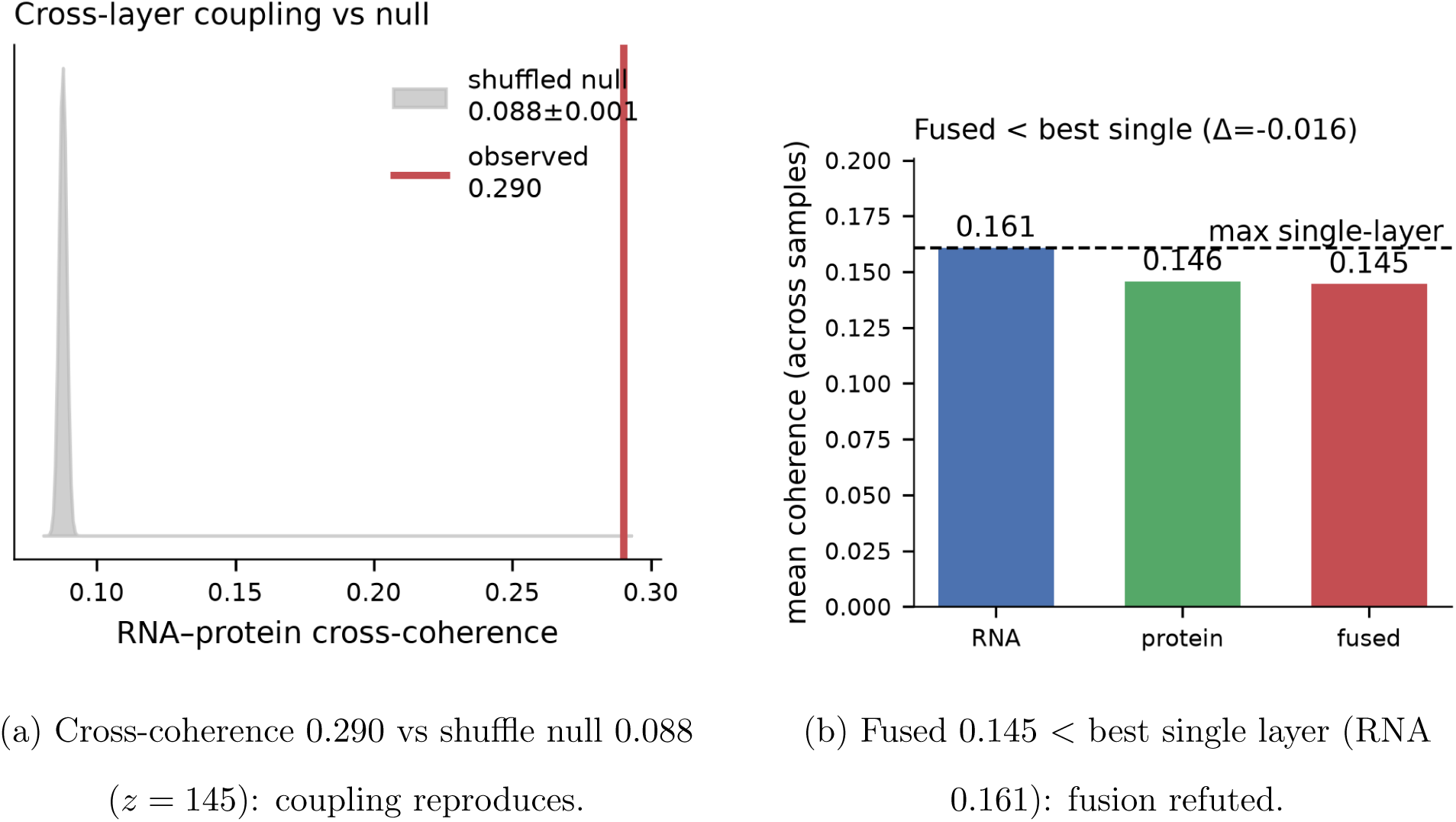
Cross-layer coupling reproduces but fusion does not beat the best single layer (CPTAC UCEC). **(a)** Observed RNA–protein cross-coherence far exceeds a sample-shuffle null (*z* = 145, *p <* 0.005, 200 shuffles), confirming genuine mRNA→protein coupling. **(b)** Coherence-weighted fusion (0.145) does not exceed the better single layer (RNA 0.161; Δ = −0.016), refuting the fused-beats-single claim on this cohort.

**FIG. 14:**
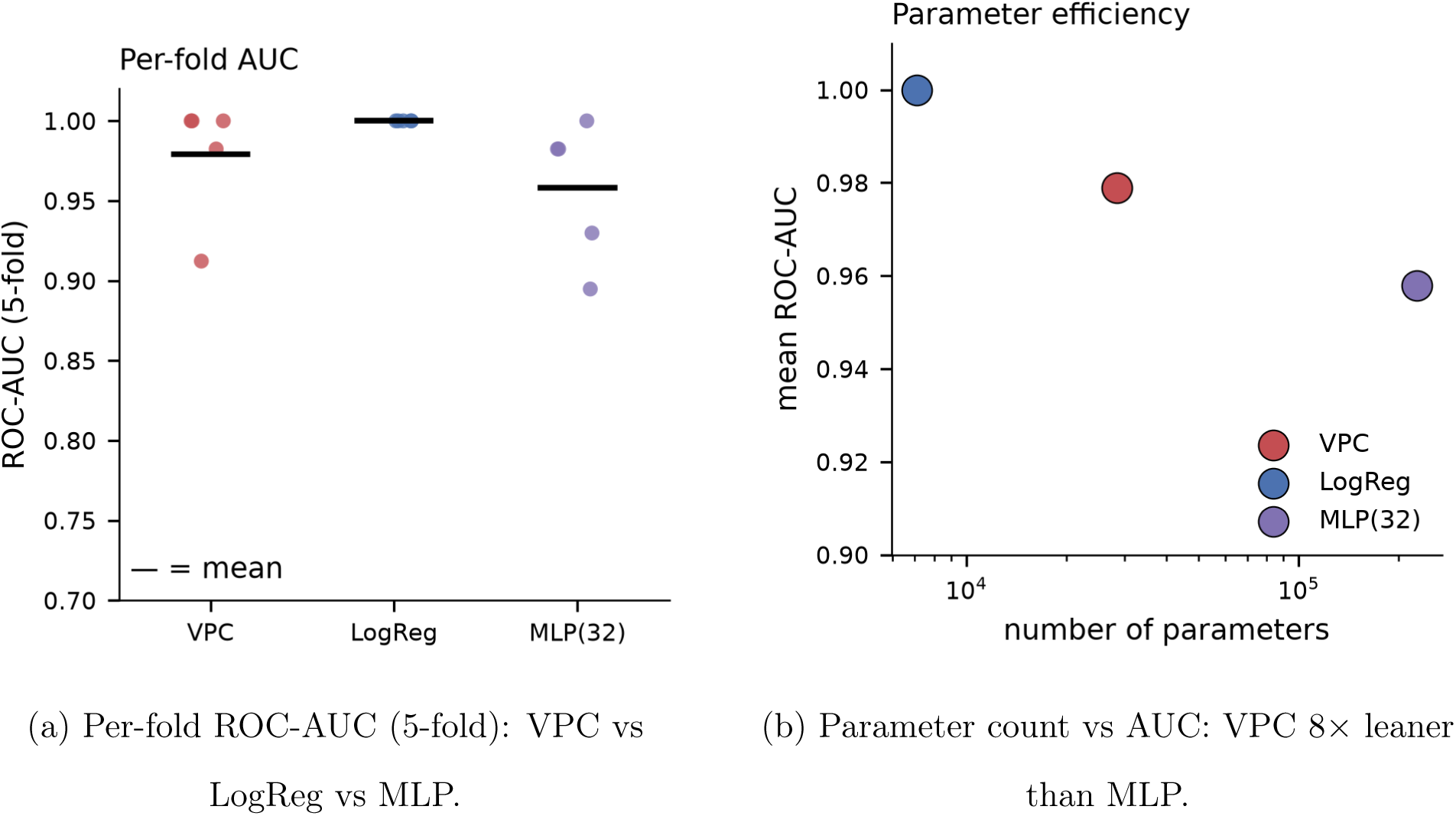
Phasor classification and parameter efficiency on real CPTAC UCEC tumour-vs-normal. **(a)** Stratified 5-fold ROC-AUC for the real phasorflow VPC, logistic regression, and a 32-unit MLP. **(b)** On the parameter–AUC frontier the VPC uses 8× fewer parameters than the MLP but is dominated by logistic regression on this near-linearly-separable, saturated task. A training-free score still separates the classes (Figure 15).

**FIG. 15:**
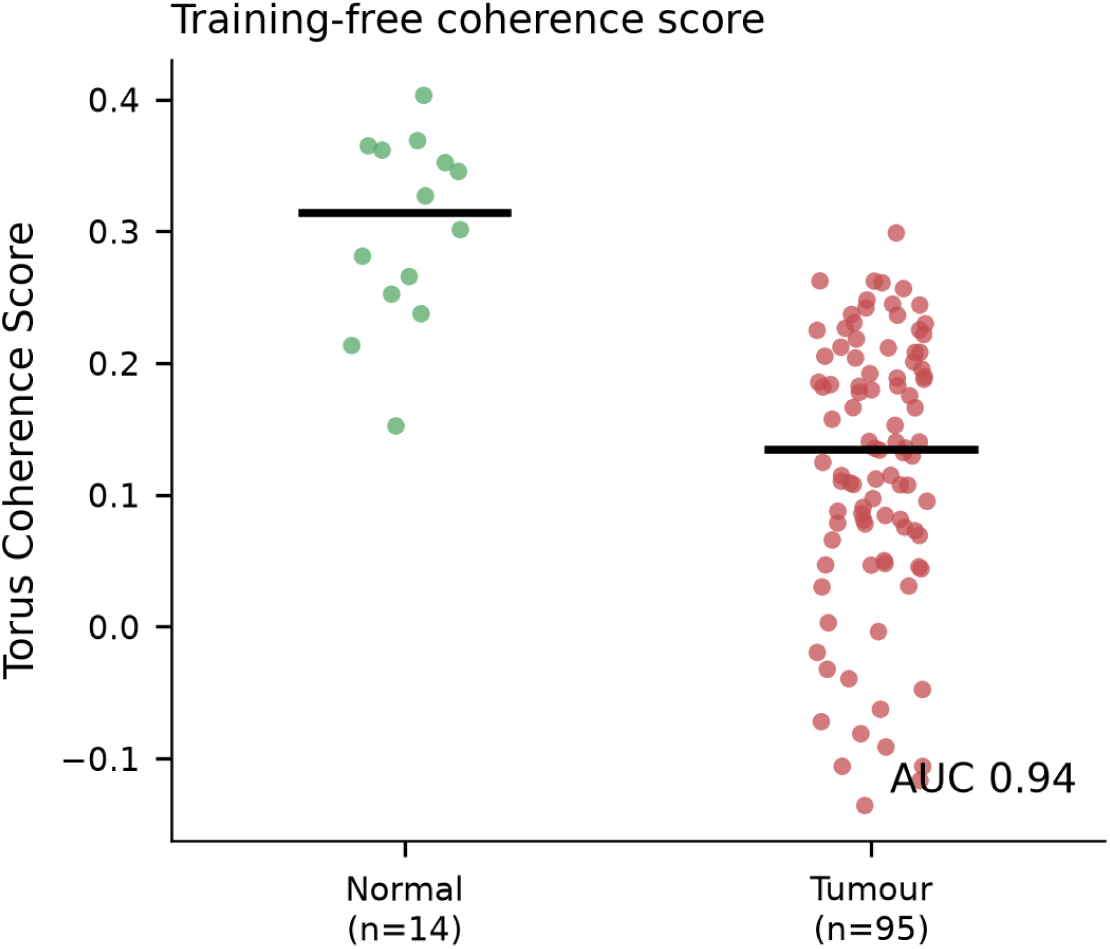
Training-free Torus Coherence Score. Label-free within-group phase coherence separates tumour from normal at AUC 0.94 with no fitting: normal samples share a consistent multi-omic phase (high within-group coherence 0.66), tumour samples are dispersed (0.15).

*Coherence-weighted fusion (does not reproduce).* The core integration claim—that coherence-weighted fusion yields a layer more coherent than any single modality—fails on this cohort: fused mean coherence is 0.145, *below* both the RNA layer (0.161) and the protein layer (0.146) (Δ = −0.016). A circular mean of two phase layers cannot exceed the more coherent input unless the layers are well aligned, and here they are only partially aligned. The result is stable under an all-9,200-gene mean-imputed sensitivity analysis (fused 0.146 vs. RNA 0.169). This scopes the fusion operator: it is a valid consensus estimator, not a coherence amplifier, and the “integration exceeds any single layer” claim needs a different fusion rule (e.g. coherence-gated rather than coherence-averaged) to hold.

### F. Phasor Classification and Parameter Efficiency

*Status: measured on real data (CPTAC UCEC tumour-vs-normal).* We ran the real phasorflow Variational Phasor Circuit (VPC; backend confirmed vpc, not the logistic fallback—4 layers, 80 epochs) against logistic-regression and MLP baselines on the fused phasor features, under stratified 5-fold cross-validation (109 samples, 7,083 features, 95 tumour / 14 normal). The verdict is *partial*.

*Parameter efficiency (holds vs. MLP, not vs. logistic regression).* The VPC reaches AUC 0.979 ± 0.034 with 28,332 parameters, versus the MLP at AUC 0.958 ± 0.039 with 226,721 parameters—comparable accuracy with 8× fewer parameters, consistent with the torus inductive bias. However, logistic regression attains AUC 1.000 ± 0.000 with 7,084 parameters, both smaller and better (Table VIII). The efficiency claim therefore holds against the deep baseline but not against the linear one.

**TABLE VIII:** Phasor classification on real CPTAC UCEC tumour-vs-normal (stratified 5-fold CV, fused RNA+protein phasor features). The real phasorflow VPC is 8× leaner than the MLP at comparable AUC but is beaten by logistic regression; AUC is saturated because the task is near-linearly separable.

| Model | AUC (5-fold) | Parameters |
| --- | --- | --- |
| Logistic regression | $1.000 \pm 0.000$ | 7,084 |
| Phasor VPC (**phasorflow**) | $0.979 \pm 0.034$ | 28,332 |
| MLP (32 hidden) | $0.958 \pm 0.039$ | 226,721 |
| Torus Coherence Score (training-free) | 0.943 | 0 |

**TABLE IX:**
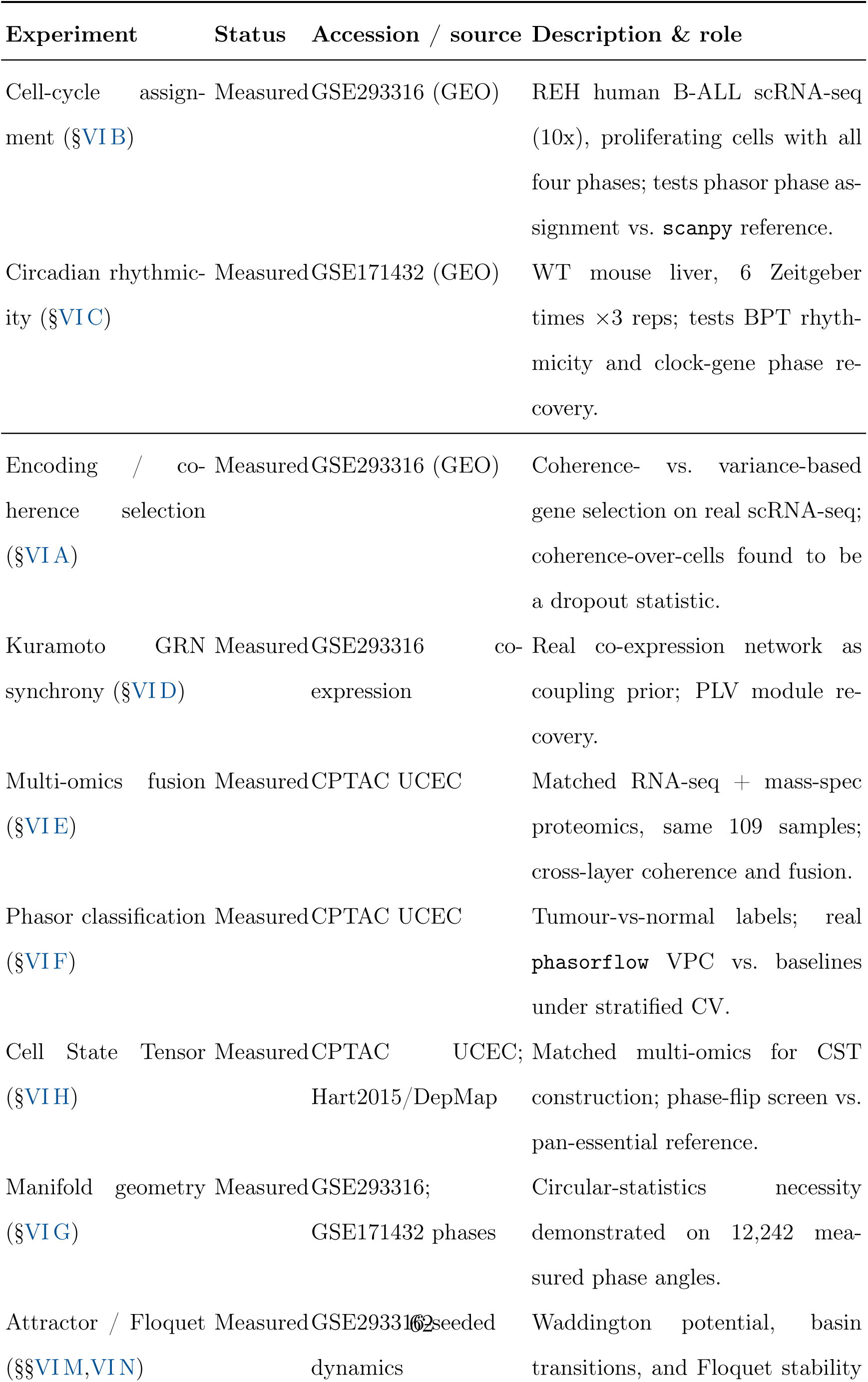
Public data sources for BioPhasor experiments. All datasets were run through the pipeline in this draft (every scenario *measured*); all accessions are open-access.

*AUC is saturated.* Tumour-versus-normal is near-linearly separable at *p* = 7,083, so every model saturates and AUC cannot rank model quality; the only discriminating axis here is parameter count, and the tiny *n* = 14 normal class makes per-fold AUC unstable (VPC std 0.034). A harder task (molecular subtype, survival) is required to test the classifier on its merits—a clear item for the GPU-scale iteration.

*Training-free coherence separation (holds).* The label-free Torus Coherence Score— consensus phase alignment with no training and no label leakage—separates the two classes at AUC 0.943, with within-group coherence far higher for normal (0.66) than tumour (0.15) tissue, supporting the claim that coherence alone carries phenotype information.

### G. Manifold Geometry Validation

*Status: measured on real data (GSE293316 + GSE171432 phases).* We validated the Riemannian operations on measured phase angles—2,000 cell-cycle angles from GSE293316 and 10,242 gene circadian angles from GSE171432 (12,242 angles total). All three sub-claims reproduce.

*(i) Exact log/exp invertibility.* The round-trip through the manifold’s log and exp maps is exact to machine precision: maximum round-trip error 1.3 × 10^−15^ rad across all 12,242 angles.
*(ii) Geodesic vs. Euclidean distance at the branch cut.* For angle pairs straddling the ±*π* boundary, naive Euclidean distance overstates the true geodesic distance by 27.9× (cells) and 47.8× (genes), with individual errors up to 344^◦^; 37% of near-boundary pairs that are neighbours on the circle are misordered as maximally distant in flat space.
*(iii) Arithmetic vs. circular/Fréchet mean.* For angles clustered near ±*π* (resultant length *R* ≈ 0.99), the arithmetic mean errs by 108.6^◦^ (cells) and 177.6^◦^ (genes—the textbook near-180^◦^ inversion) relative to the circular mean, while the package’s phasor_mean and manifold frechet_mean agree with the circular mean to 0^◦^ (Figure 16). On measured phase data, circular statistics are required, not merely convenient.

**FIG. 16:**
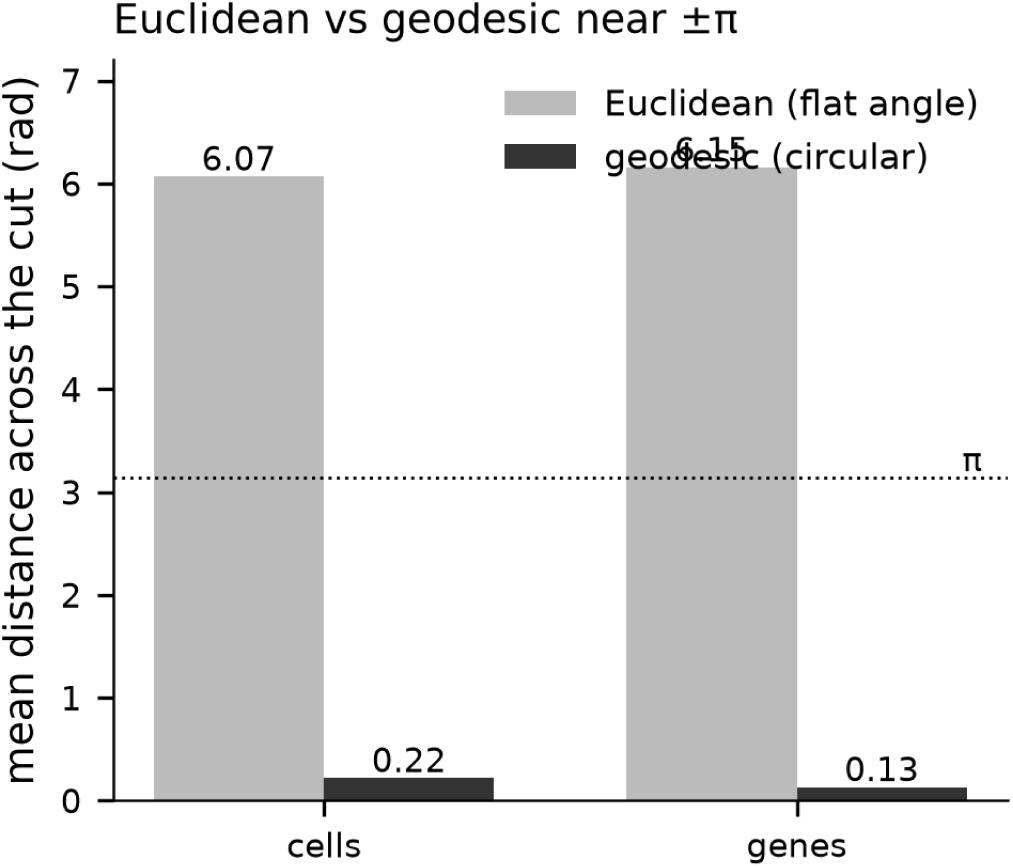
Euclidean distance overstates geodesic distance near the branch cut (measured phases, GSE293316 cells + GSE171432 genes). Mean Euclidean vs geodesic (circular) distance for phase pairs straddling the ±*π* cut: naive flat distance overstates the true circular distance by 28× (cells) to 48× (genes). log/exp round-trip is exact to 10^−15^ rad. The mean-estimator consequence is shown in Figure 17.

**FIG. 17:**
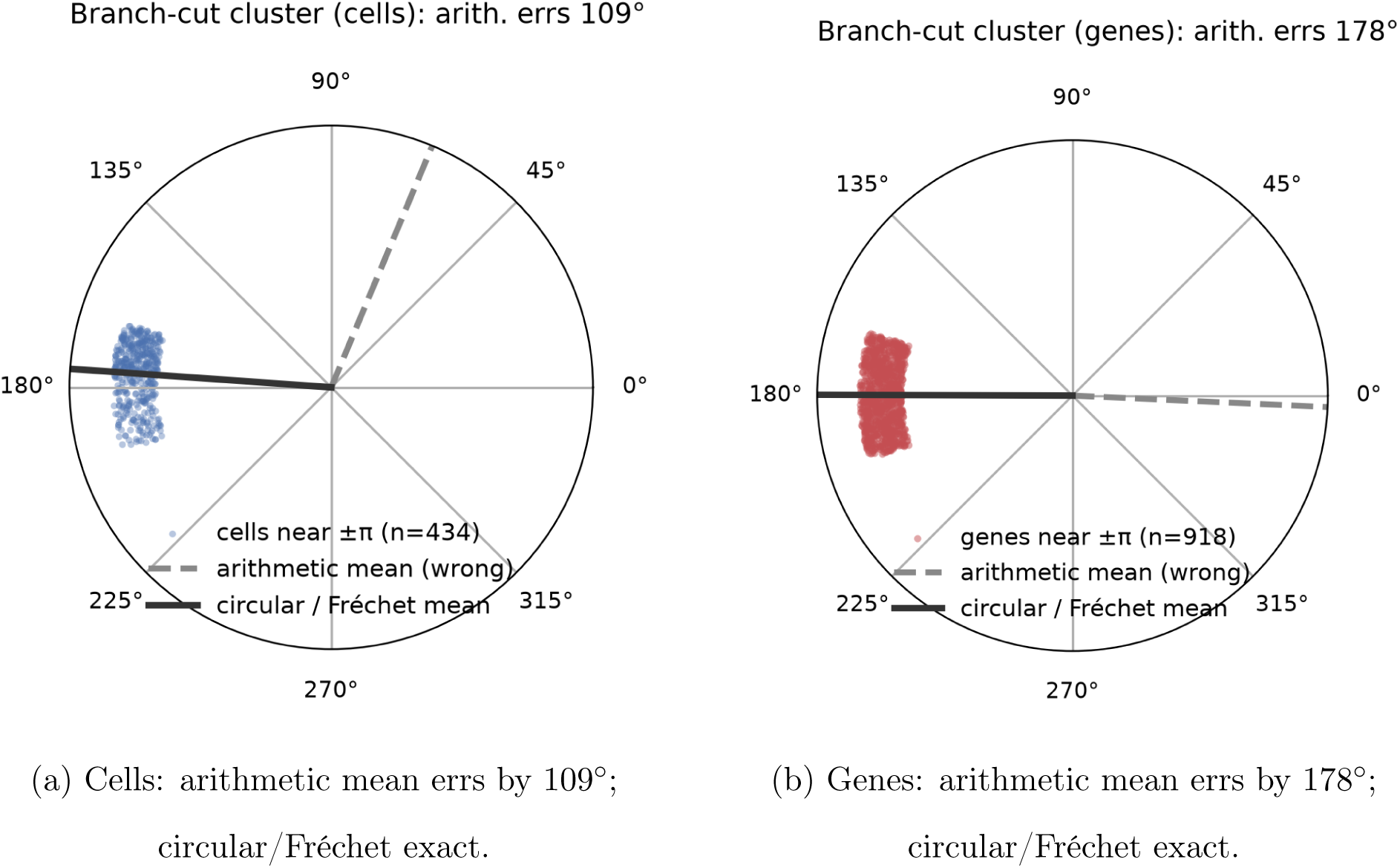
The arithmetic mean fails on wrapped phases; the circular mean is exact. For a cluster straddling the ±*π* branch cut, the naive arithmetic mean (which ignores wrapping) lands nearly opposite the true centre — erring by 109^◦^ for the cell cluster **(a)** and 178^◦^ for the gene cluster **(b)** — whereas the circular/Fréchet mean recovers the correct centroid exactly.

### H. Cell State Tensor Construction and Dynamics

*Status: measured on real data (CPTAC UCEC).* We constructed the Cell State Tensor from the matched CPTAC UCEC RNA and protein phase layers and tested two claims with the unmodified CSTDynamics and phase-flip routines at documented defaults. Both fail on this cohort; the verdict is *does-not-reproduce*, and the failure is informative about the operator’s domain of validity.

*(i) Coherence under Kuramoto evolution (fails).* Under the documented CSTDynamics evolution (*K* = 3, 300 steps) global coherence *decreases* from 0.612 to 0.519 and phase entropy *increases* from 3.146 to 3.277—the opposite of the predicted rise in coherence and fall in entropy. Coherence-weighted Kuramoto coupling on two heterogeneous omics layers drives them apart rather than into a common phase.
*(ii) Phase-flip knockout screen (fails).* Rotating each of the 7,083 genes by *π* and ranking by resulting global-coherence loss (verified bit-identical to the package phase_flip loop), the top-100 hits contain *zero* pan-essential genes against a family-level reference (Hart et al. 2015 core-essential + DepMap common-essentials; 249 essentials present): fold enrichment 0.0×, hypergeometric *p* = 1.0, versus a 3.5% background rate. The top hits (CLIC6, TNXB, SPARCL1, …) are tumour/normal-differential genes, not fitness-essential ones (Figure 18). Diagnosis: the phasor “essentiality” proxy tracks differential expression magnitude, not genetic dependency, so it cannot stand in for a CRISPR essentiality screen without a fitness-anchored coupling model. This is a genuine negative that redirects the knockout-screen design for the full-scale iteration.

**FIG. 18:**
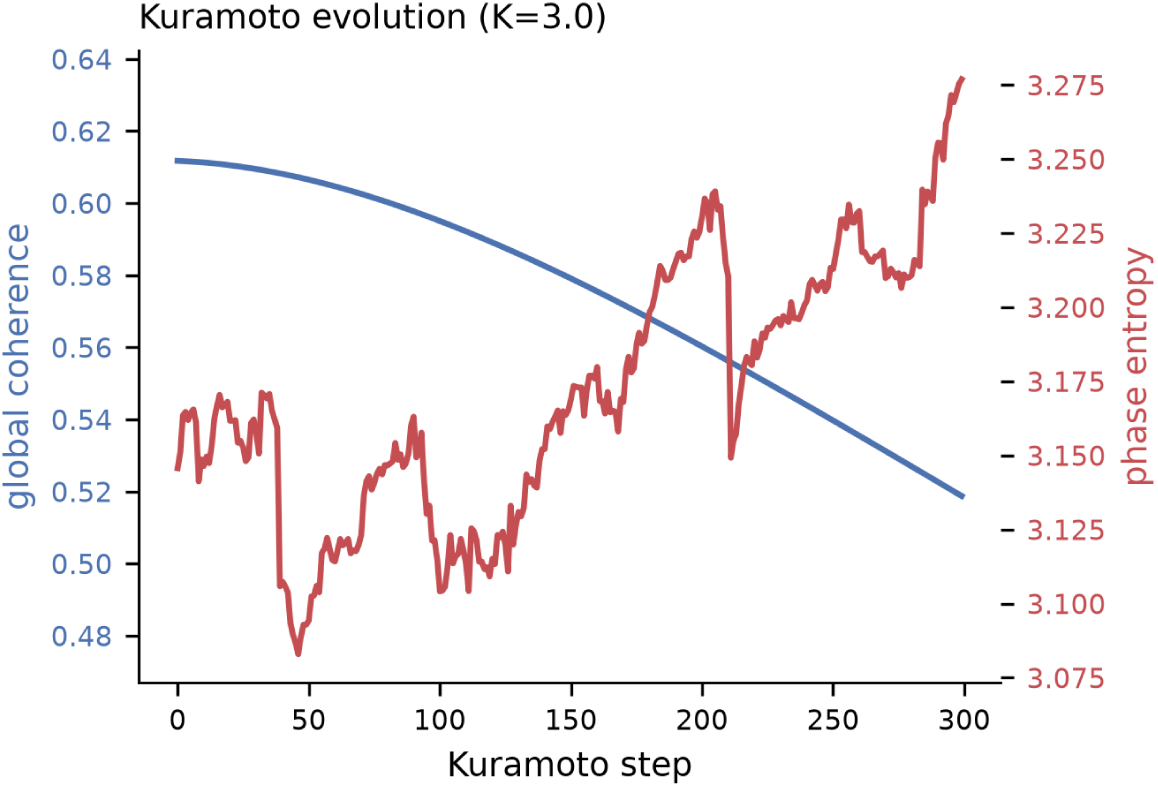
Cell State Tensor dynamics run opposite to the claim (real CPTAC UCEC). Under the documented CSTDynamics Kuramoto evolution (*K* = 3, 300 steps), global coherence *decreases* (0.61 → 0.52) while phase entropy *increases* (3.15 → 3.28) — the opposite of the predicted coherence-increasing convergence. The associated knockout screen is in Figure 19.

**FIG. 19:**
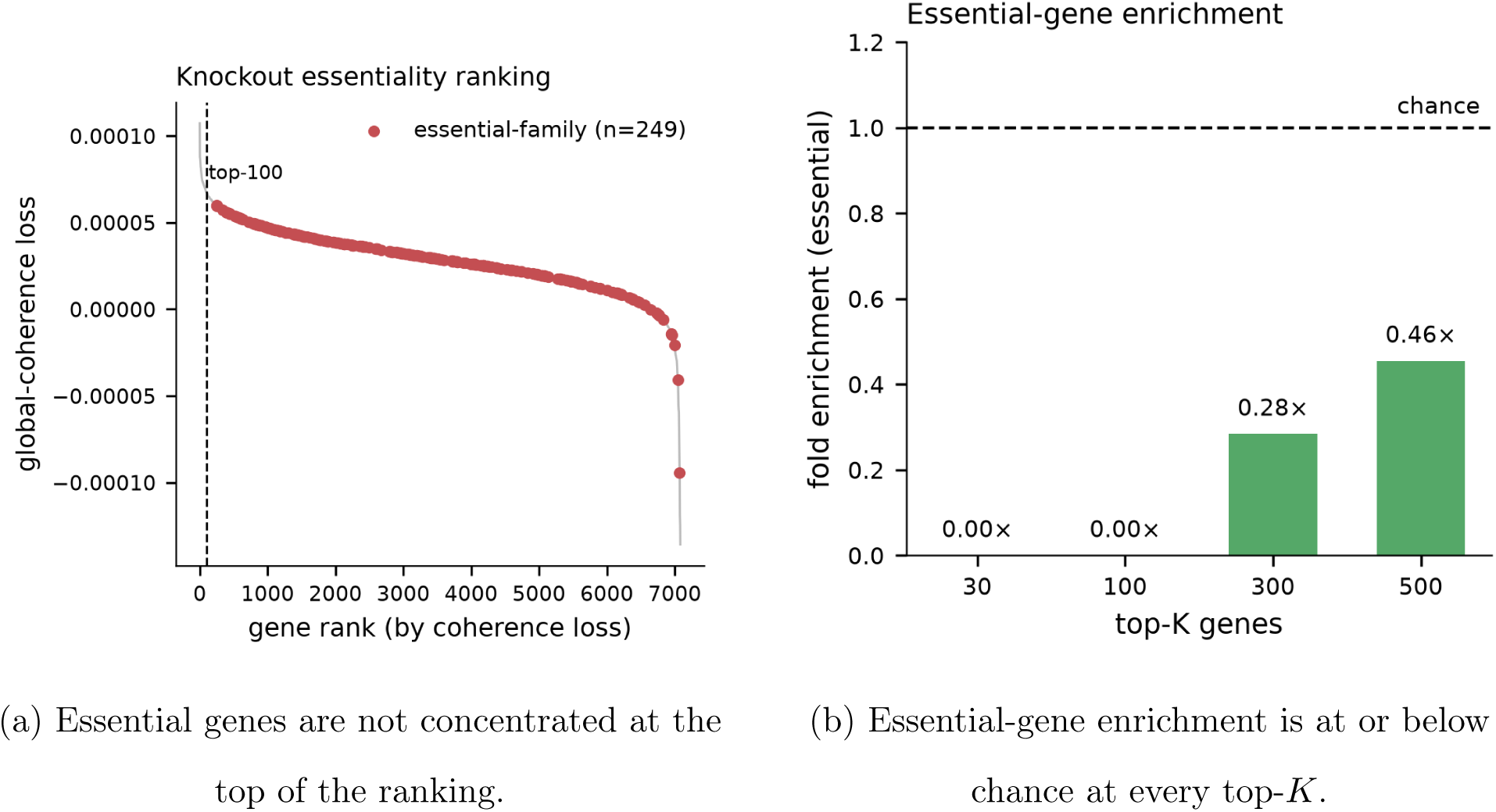
Phase-flip knockout screen recovers no essential-gene signal (real CPTAC UCEC). **(a)** Genes ranked by coherence loss on phase-flip knockout; known essential-family genes (Hart2015 / DepMap) are scattered through the ranking, not concentrated at the top. **(b)** Fold enrichment of essentials in the top-*K* hits is at or below chance for *K* ∈ {30, 100, 300, 500} (top-100 fold 0×, *p* = 1.0); the top hits are differential, not essential, genes.

### I. Time-Resolved CST Profiles on Real Biological Time-Series

*Status: measured on real time-series (GSE293316 cell-cycle pseudotime; GSE171432 circadian).* The preceding CST results characterise a static snapshot; here we test the temporal behaviour the framework predicts—that the CST features G*_t_* (global coherence), E*_t_* (phase entropy), and *R_t_* (state velocity, Eqs. 34–37) trace a coherent trajectory as the cell moves through an oscillatory program—and the Temporal Update Rule (Eq. 40) that stabilises these estimates. Because per-cell coherence is a dropout artefact on sparse single-cell data (Section VI A), every single-cell CST here is *pseudobulked* within its time bin. The verdict is *partial* : the predicted temporal structure appears clearly when the time axis is densely sampled, and is sampling-limited when it is not.

*(i) Cell-cycle pseudotime (reproduces).* Ordering 4,000 proliferating REH cells on the continuous cell-cycle angle arctan 2(G2M, S) and pseudobulking into 10 equal-count pseudo-time bins (381 marker + top-variance genes), the per-bin CST global coherence rises from G = 0.370 in early G1 (bin 0) to a G2/M peak of 0.845 (bin 7; net rise +0.475) and then crashes to its minimum 0.262 at the post-mitotic bin (bin 9), with phase entropy E*_t_* anti-correlated to coherence throughout (Figure 20a). The state velocity *R_t_* = || Δ*ϕ |||* _1_*/N* peaks at exactly the two phase boundaries—bin 0 (the cyclic M→G1 wrap) and bin 4 (G1→S)— identifying the transitions the framework predicts should be the fastest-moving points of the trajectory. The rise is not strictly monotonic (a minor dip at bin 3), but the G1-low / G2M-high / post-mitotic-crash arc is unambiguous. This is a genuine positive: the CST temporal profile recovers cell-cycle structure directly from real single-cell data.
*(ii) Circadian time (partial, sampling-limited).* On the WT mouse-liver series (GSE171432; 60 rhythmic genes = 14 core-clock ∪ 53 with rhythmicity score ≥ 0.30) each of the six Zeitgeber times yields a CST whose phase entropy E*_t_* oscillates with genuine 24 h structure— oscillation amplitude 0.0554 versus an arrhythmic-gene null mean of 0.0186 (*p <* 0.001, peak ZT18.4)—but whose coherence trace G*_t_* does not clear the null (amplitude 0.0216 vs. null 0.0466, *p* = 0.92; Figure 21a). Six timepoints sit at the Nyquist floor for a 24 h rhythm, the same sampling limit that caps circadian recall at 0.43 in Section VI C; the entropy axis carries the temporal signal that the coherence axis cannot resolve at this density.
*(iii) Temporal update rule (reproduces).* On a noisy 40-step CST coherence trace the Temporal Update Rule (Eq. 40) behaves as specified: fixed EMA smoothing (*λ* = 0.85) cuts trace variance three-fold (0.0323 → 0.0107) but lags a genuine state transition (tracking error 0.416 over the transition window). The adaptive-*λ* variant—dropping *λ* to 0.25 at the four steps where *R_t_* spikes—retains most of the smoothing (variance 0.0175) while tracking the same transition roughly three times more closely (error 0.140), the responsiveness–stability trade-off the adaptive rule is designed to make (Figure 21b).

**FIG. 20:**
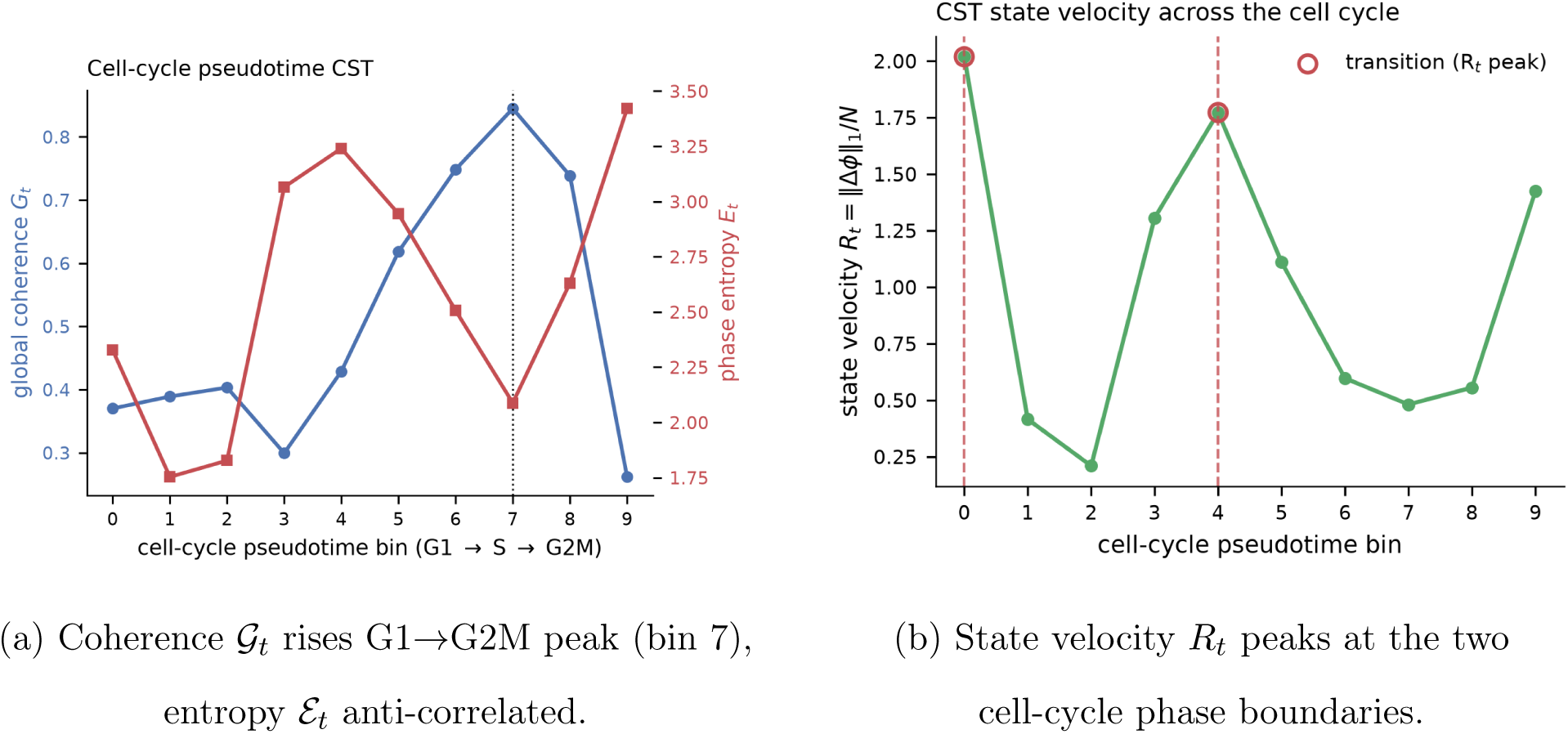
CST temporal profile reproduces cell-cycle structure (real GSE293316, pseudobulked pseudotime). **(a)** Across 10 pseudotime bins (4,000 cells ordered on the continuous cell-cycle angle), CST global coherence G*_t_* rises from early G1 (0.370) to a G2/M peak (0.845, bin 7) and crashes post-mitosis (0.262, bin 9), with phase entropy E*_t_* anti-correlated. **(b)** State velocity *R_t_* = || Δ*ϕ ||* _1_*/N* peaks at exactly the two phase boundaries (the cyclic M→G1 wrap at bin 0 and G1→S at bin 4). Per-bin CSTs are pseudobulked to avoid the single-cell dropout artefact of Section VI A.

**FIG. 21:**
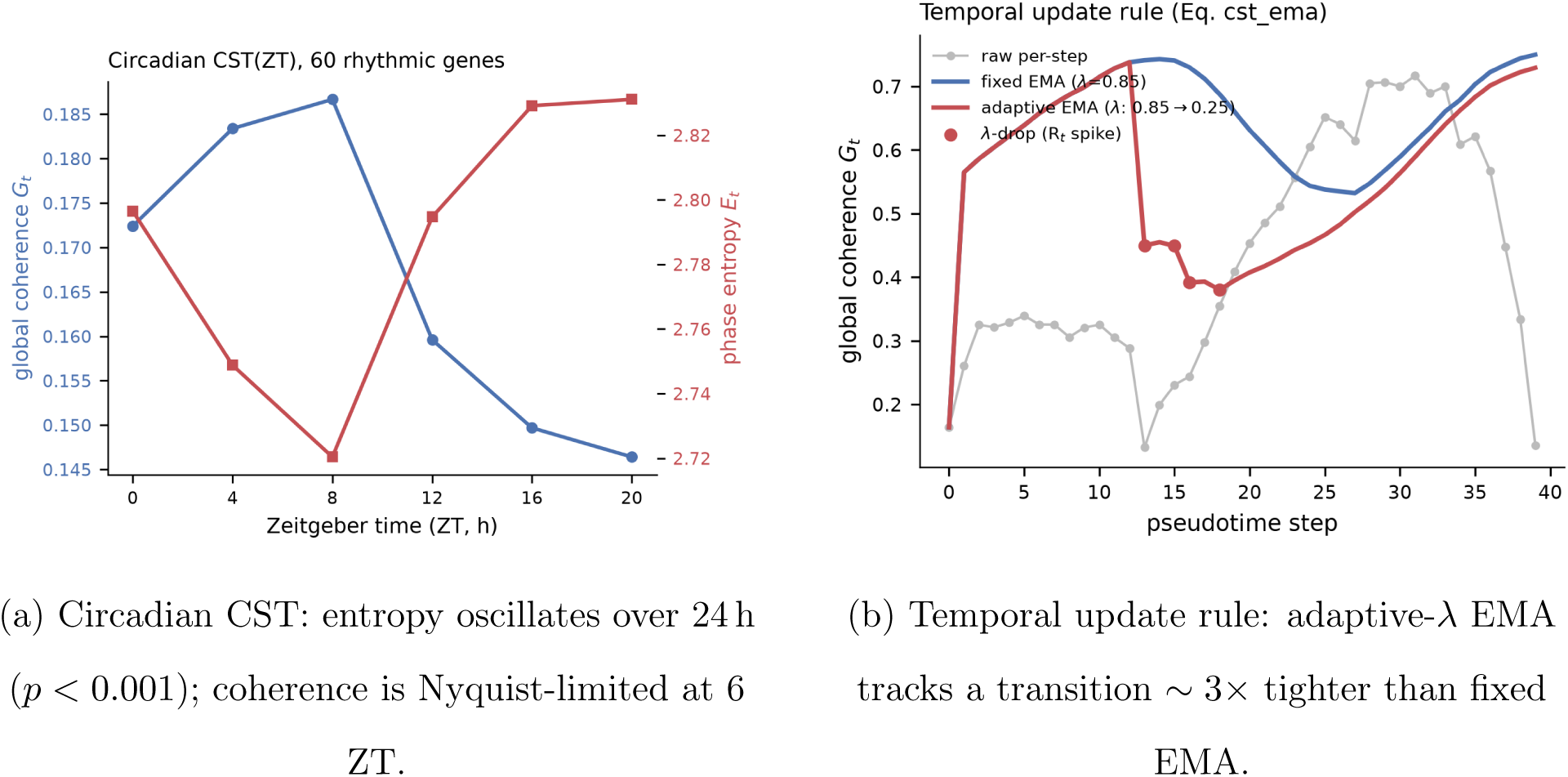
Circadian CST profile and the temporal update rule. **(a)** On the WT mouse-liver circadian series (60 rhythmic genes), CST phase entropy E*_t_* oscillates with genuine 24 h structure (amplitude 0.0554 vs. arrhythmic null 0.0186, *p <* 0.001), while coherence G*_t_* does not clear the null at six ZT points—the Nyquist sampling floor that also caps circadian recall in Section VI C. **(b)** The Temporal Update Rule (Eq. 40) on a noisy coherence trace: fixed EMA (*λ* = 0.85) cuts variance three-fold but lags a state transition (error 0.416); adaptive-*λ* (dropping to 0.25 at *R_t_* spikes, marked) keeps the smoothing yet tracks the transition ∼ 3× more closely (error 0.140).

### J. Tensor-Network Factorization and Uncertainty of the CST

*Status: measured on real data (CPTAC UCEC).* We tested the tensor-network (MPS/tensor-train) factorization the framework proposes for compact CST storage and online updates (Eq. 39), together with the uncertainty-aware CST (Section IV). The verdict is *partial*, and the boundary is informative: tensor-train does not compress a single CST snapshot, but it does amortize storage across a growing history. (All tensor-train decompositions use a real/imaginary-stacked representation of the complex CST; tensorly 0.9.0’s complex TT-SVD is numerically unreliable—its reconstruction error grows with bond dimension—so the complex tensor is split into real and imaginary channels before decomposition, as noted in the Methods.)

*(i) A single CST is not low-rank (does-not-reproduce).* The gene-resolved CPTAC CST (7,083 genes × 2 modalities × 109 samples) has a high-entropy gene-mode spectrum: rank 50 captures only 50% of the spectral energy and rank 150 reaches 86% (full rank 218; Figure 22a). Consequently the tensor-train never reaches even 10% reconstruction error at any bond dimension—the best achieved is 65% error at 3.2× compression (bond 128), and pushing compression above ∼ 3× costs more than 65% error (Figure 22b). The regulatory axis carries genuinely high-dimensional inter-gene structure that no small bond dimension captures, so Eq. 39 does not deliver useful single-snapshot compression on this real CST.
*(ii) CST history storage is sublinear (reproduces).* The streaming-storage motivation for the factorization holds at the history level. Stacking CST snapshots into a growing 4-D history tensor (*L* × *R* × *T* × *H*), dense storage grows linearly in history length *L* while the tensor-train’s history-axis bond saturates, so TT storage grows sublinearly: at fixed bond 40 the compression ratio rises from 2.7× at *L* = 1 to 5.1× at *L* = 64 (Figure 23a). Tensor-train amortizes the redundancy shared *across* snapshots even though it cannot compress any single one; a full re-decomposition (a proxy for the rank-adaptive online update) costs on the order of a second at *L* = 64, feasible for periodic rather than per-cell updates.
*(iii) Uncertainty is heterogeneity, not dropout (informative negative).* Bootstrap-resampling the samples *B* = 40 times and rebuilding the CST gives a per-entry circular variance (Eq. 41) that is high (mean 0.80) and—contrary to the manuscript hypothesis that CST uncertainty is dropout-driven—essentially uncorrelated with expression level (Spearman *ρ* = 0.08, flat across expression quintiles; Figure 23b). On this matched bulk multi-omics cohort the CST’s uncertainty reflects genuine inter-sample biological heterogeneity rather than measurement dropout, which refines where the dropout-flagging use of **Σ***_t_* applies.

**FIG. 22:**
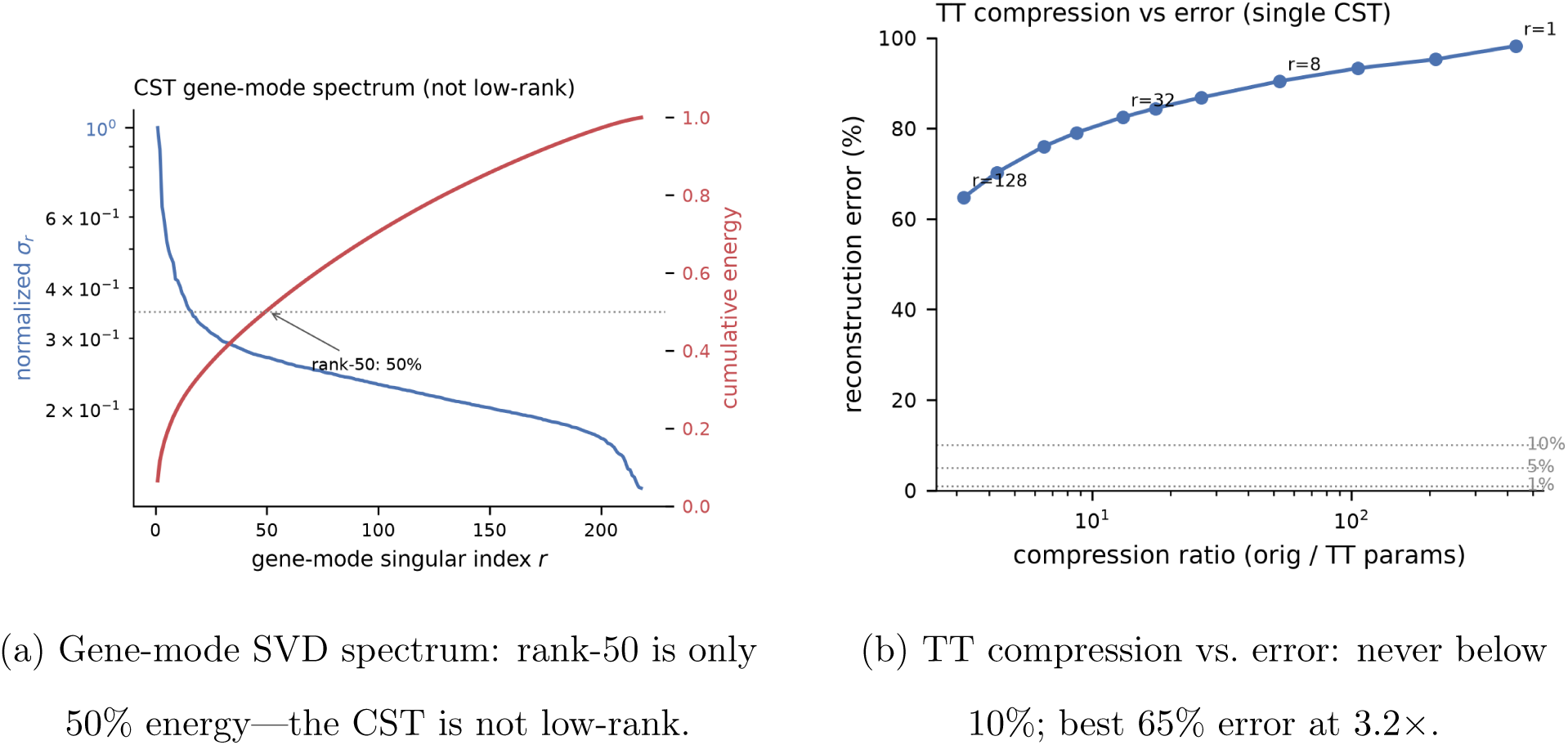
Tensor-train does not compress a single real CST (CPTAC UCEC). **(a)** The gene-mode singular-value spectrum is high-entropy—rank 50 captures 50% and rank 150 captures 86% of the energy (full rank 218). **(b)** Across the bond-dimension sweep the tensor-train reconstruction error never falls below 10% (dotted reference lines at 10*/*5*/*1%); the best operating point is 65% error at 3.2× compression. The regulatory axis is genuinely high-dimensional.

**FIG. 23:**
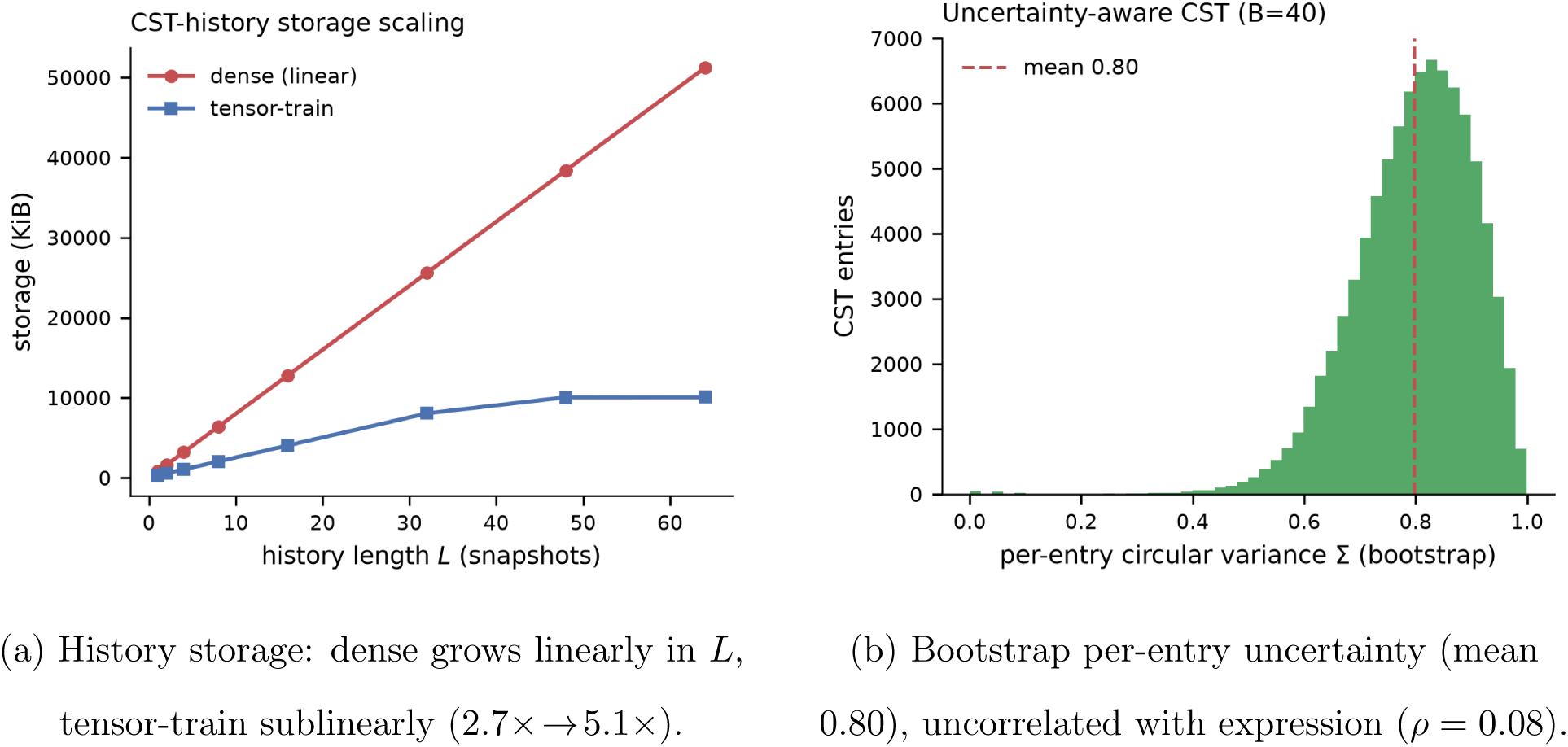
CST-history storage scaling and uncertainty (real CPTAC UCEC). (a) Stacking CST snapshots into a growing history tensor, dense storage grows linearly in history length *L* while tensor-train storage grows sublinearly (compression 2.7× at *L* = 1 to 5.1× at *L* = 64, fixed bond 40)—the factorization amortizes cross-snapshot redundancy. (b) Per-entry bootstrap circular variance (*B* = 40) is high (mean 0.80) and essentially independent of expression level (Spearman *ρ* = 0.08), so the CST’s uncertainty on this bulk cohort reflects biological heterogeneity, not measurement dropout.

### K. A Pathway Atlas Roots the Regulatory Axis and Makes the CST Compressible

*Status: partial (real CPTAC UCEC RNA +protein). Rooting the regulatory axis in a pathway atlas reproduces low-rank behaviour on that axis, but not for the full tensor*.

The tensor-network result above (Section VI J) found that the flat gene-resolved CST is not low-rank: its gene-mode spectrum needs rank 50 for half the energy. That is a property of an *unstructured* regulatory axis—a flat list of 7083 genes. Section IV B argued that biology supplies the missing structure through pathways, exactly as an anatomical atlas supplies structure to a brain tensor’s region axis. We test that claim directly on the same real. CPTAC UCEC cohort (RNA + protein, 109 samples), building a pathway-aggregated CST of shape 50 × 2 × 109 (50 MSigDB Hallmark programs × two modalities × samples), each pathway carrying the circular-mean phasor of its member genes. The atlas covers 2405 of the 7083 co-observed genes (34%); we report the fair, per-component comparison of captured energy so the result does not merely restate the 7083 → 50 dimensionality reduction.

Imposing pathway structure collapses the regulatory spectrum (Fig. 24a). A *single* regulatory component of the pathway CST captures 52.5% of the tensor’s energy, and 14 components reach 90%; the flat-gene CST needs rank 50 for 50% and rank 150 for 86% (Section VI J), with its leading component capturing only 6.6%. In other words, pathway structure achieves in one component what the flat gene axis needs about fifty to match—the qualitative signature of the region-aggregated regime that makes a brain tensor low-rank. The most phase-coherent Hallmark programs across the cohort (Fig. 25) are proliferation-linked (e2f_targets, myc_targets_v1/v2, g2m_checkpoint) together with oxidative_phosphorylation, while immune and xenobiotic-metabolism programs are least coherent—a biologically interpretable readout consistent with proliferation being the dominant coordinated axis in endometrial carcinoma, and the direct analogue of reading structure off a brain atlas’s regions.

**FIG. 24:**
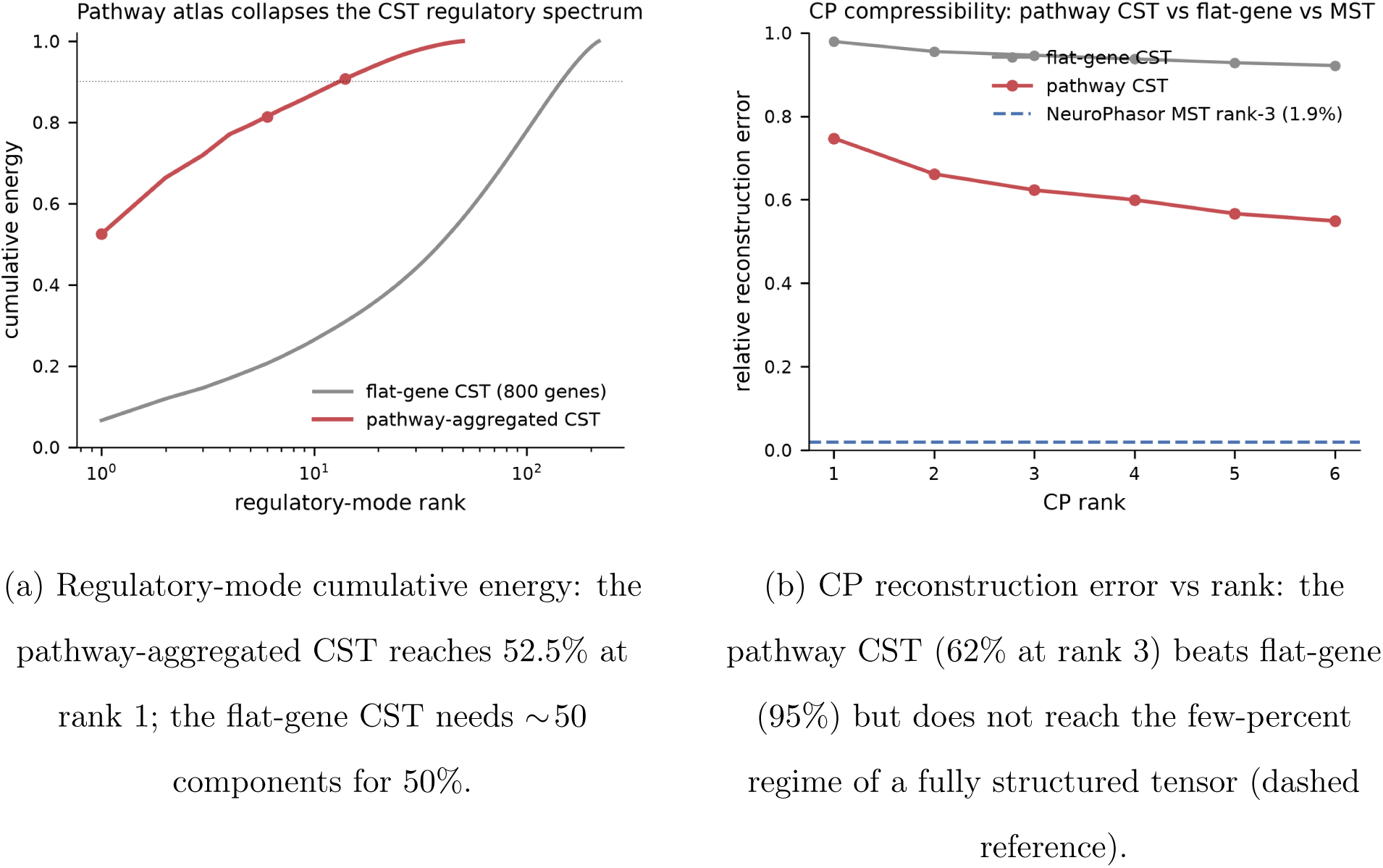
A pathway atlas roots the CST regulatory axis (real CPTAC UCEC). Aggregating 7083 genes onto 50 MSigDB Hallmark programs supplies the regulatory axis with biological structure—the multi-omics counterpart of an anatomical atlas. **(a)** This collapses the regulatory-mode spectrum toward the low-rank regime; **(b)** full-tensor CP compressibility remains partial because the cohort’s sample axis carries irreducible tumour heterogeneity (see text).

**FIG. 25:**
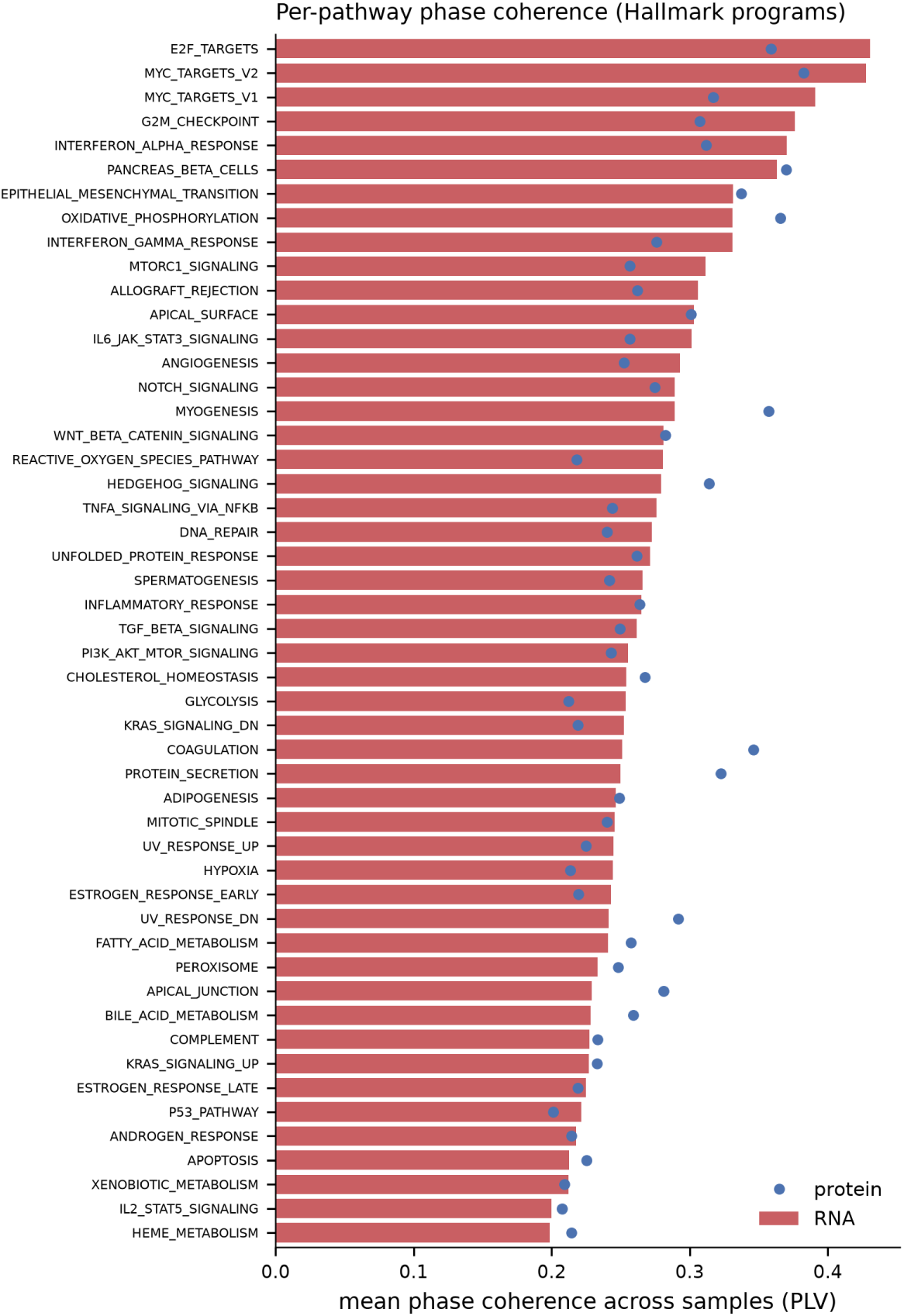
Per-pathway phase coherence across the UCEC cohort. Mean across-sample phase coherence (PLV) of each Hallmark program, RNA (bars) and protein (points). Proliferation programs (e2f, myc, g2m) and oxidative_phosphorylation are the most phase-coherent; immune and xenobiotic-metabolism programs the least—an interpretable biological readout off the pathway atlas, analogous to reading structure off a brain atlas’s regions. Protein coherence is generally lower than RNA, consistent with post-transcriptional buffering.

The verdict is *partial* because the collapse is confined to the regulatory axis. A full CP decomposition of the pathway CST still does not reach the low-rank point of a genuinely low-dimensional tensor: its rank-3 reconstruction error is 62.3% (versus 94.6% for the flat-gene CST, but far from the few-percent regime attainable when all three axes are structured) (Fig. 24b). The reason is structural and honest, not an atlas deficiency. CPTAC UCEC is a patient *cohort*, so the third axis is 109 heterogeneous tumours rather than a low-dimensional time or frequency axis; that axis alone needs rank 17 for 90% of its energy, which bounds how well any rank-3 set of shared sample loadings can reconstruct the tensor. Full-tensor compressibility of the CST is therefore expected to require a structured third axis—matched multi-omics *time series*—which the pathway atlas cannot supply on its own. The atlas does what it should: it makes the regulatory axis interpretable and low-rank, the piece BioPhasor most needed to stop treating genes as an unstructured bag.

### L. Central-Dogma Cross-Modal Phase Coupling (“Omics-PAC”)

*Status: reproduces (real CPTAC UCEC RNA +protein). Across patients, mRNA phase significantly organises protein amplitude, above a sample-permuted surrogate null*.

The modality axis (Section IV B) is what most sharply distinguishes multi-omics from a neural tensor: it is causal and directional (DNA → mRNA → protein). We make the cross-modal coupling along this axis a first-class measurement—the multi-omics analogue of cross-frequency phase–amplitude coupling—and ask whether, across the 109 CPTAC UCEC patients, the *phase* of a gene’s mRNA phasor organises the *amplitude* of its protein. For each gene we bin the mRNA phase into *K* = 9 bins across samples, take the mean protein amplitude per bin, and score the non-uniformity of that distribution with a Tort-style modulation index (MI); the null circularly permutes the sample correspondence between modalities (*N* = 200 permutations), breaking coupling while preserving each modality’s marginal distribution. Because CPTAC is a cohort rather than a time course, this is explicitly *cross-sample* coupling, not a temporal translation lag—the temporal-lag form is future work requiring matched multi-omics time series.

The coupling is present and highly significant (Fig. 26a, 26b). The aggregate per-gene MI is 0.0177 against a null mean of 0.0077 (*z* = 190, empirical *p* = 0.005, the permutation floor at *N* = 200), and a pooled circular–linear correlation between mRNA phase and protein amplitude gives *r* = 0.295 (*z* = 51, *p* = 0.005). The effect is broad rather than driven by a few genes: 5140 of 7083 genes (72.6%) are individually significant at *p <* 0.05 (Fig. 27). It is also biologically structured—the most strongly coupled genes form a coherent epithelial-polarity / cell-junction module (*OCLN*, *CXADR*, *MYO5B*, *SLC9A3R1*, *LAD1*, *EPS8*) together with estrogen-signalling genes (*ESR1*, *ESRP2*), plausible for endometrial carcinoma—and it is tumour-specific: the pooled coupling is *r*= 0.295 across the 95 tumours but only *r* = 0.029 across the 14 normal samples.

**FIG. 26:**
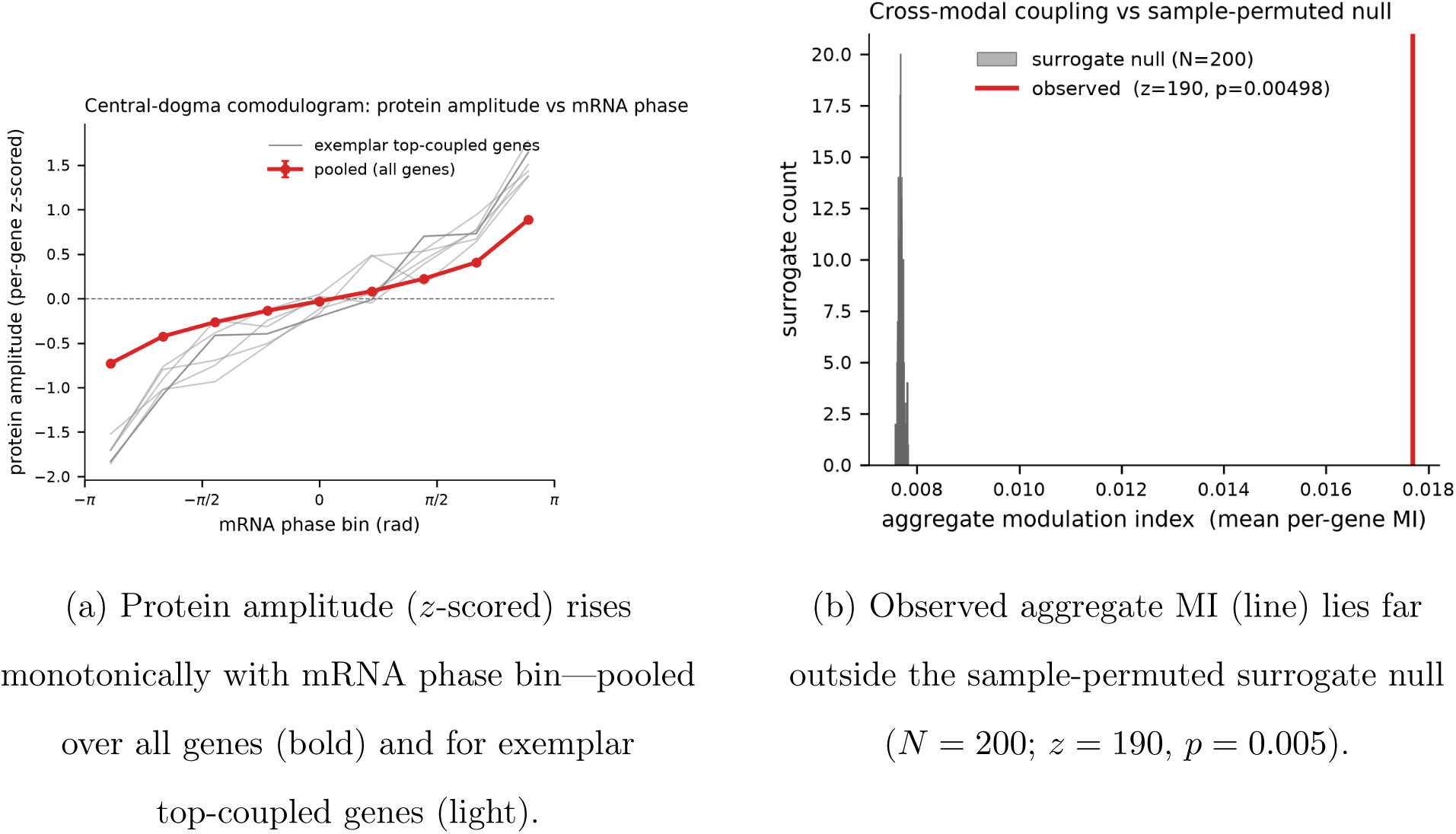
Central-dogma cross-modal phase coupling on real CPTAC UCEC. Across 109 patients, the phase of a gene’s mRNA phasor organises the amplitude of its protein—the multi-omics analogue of cross-frequency phase–amplitude coupling, along the directional modality axis that has no neural counterpart. **(a)** Comodulogram; **(b)** significance against a sample-permuted null.

**FIG. 27:**
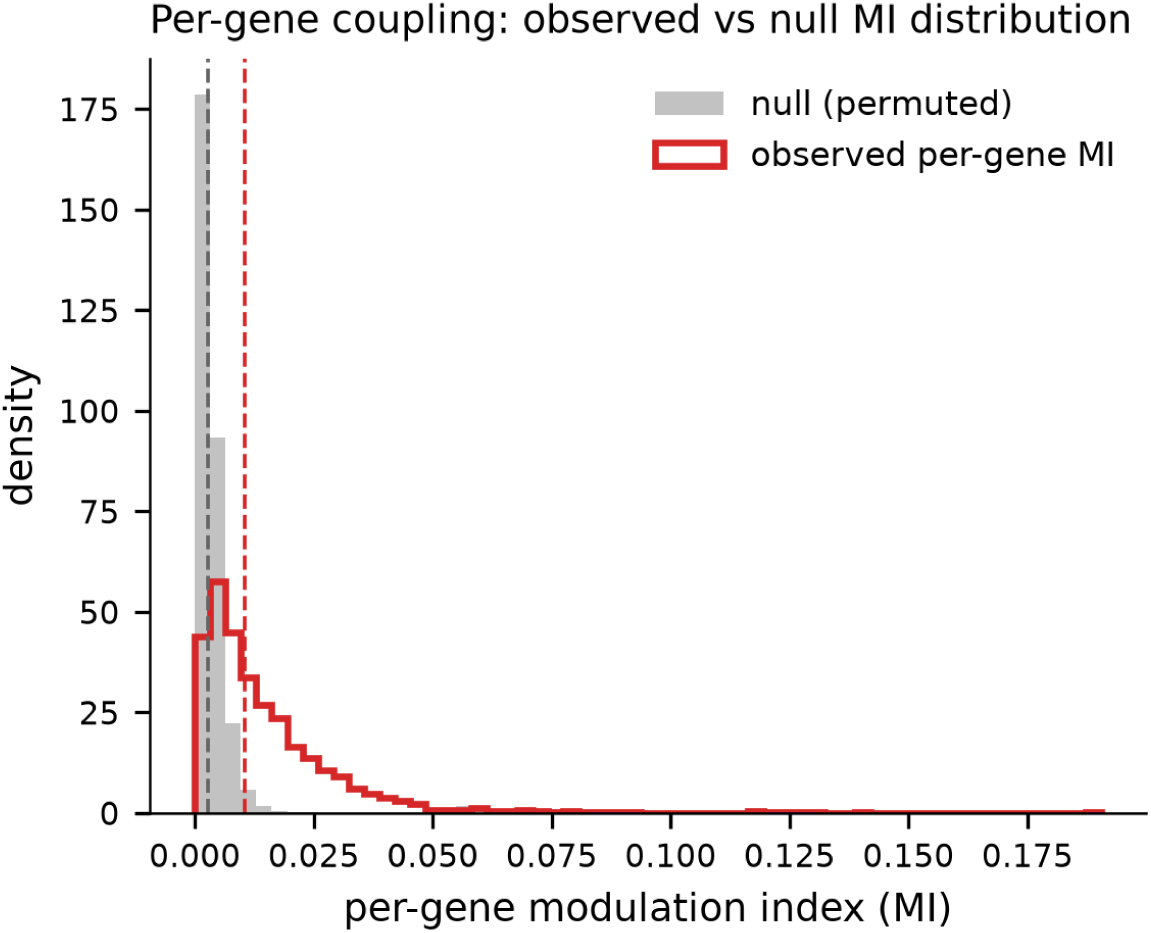
Per-gene coupling is broad. Distribution of per-gene modulation index, observed (outline) versus sample-permuted null (filled): 5140*/*7083 genes (72.6%) are individually significant at *p <* 0.05, so the central-dogma coupling is a genome-wide property of the cohort rather than a few-gene artefact.

We label this *reproduces* because the coupling clears the surrogate null decisively and consistently across per-gene, pooled, and stratified views. Two honest caveats bound the claim. First, the absolute magnitude is modest (aggregate MI only ∼ 2.3× the null, pooled *r* ≈ 0.30), matching this project’s earlier finding that mRNA–protein phase coupling is real and significant yet coherence-weighted fusion does not beat the better single layer (Section VI H); a significant coupling is not the same as a strong predictor. Second, the normal-sample estimate rests on only 14 samples. The result nonetheless establishes the modality axis as a genuine, non-neural BioPhasor phenomenon with a proper null—the piece that makes the CST a multi-omics object rather than a phasor tensor by analogy.

### M. Attractor Landscape and Waddington Quasi-Potential

*Status: measured on real-seeded dynamics (GSE293316 co-expression).* We tested the attractor-landscape interpretation on a multi-regime BioKuramoto trajectory whose coupling topology is a real GSE293316 co-expression network (165 oscillators, four coupling regimes). We are explicit about scope: the coupling is real, but the trajectory is package-simulated and the basin labels are the package’s placeholder state names, not biological cell-fate assignments—this is a method demonstration on real-seeded dynamics, not a fate-mapping result. All three sub-claims reproduce.

*(i) Waddington quasi-potential.* Kernel-density estimation on the PCA-projected attractor features yields a quasi-potential *U* (*ϕ*) = − log *p*(*ϕ*) with 4 distinct basins; the deepest well is also the most occupied (occupancy 0.51) and longest-residence basin, i.e. the most stable state is deepest (Figure 28).
*(ii) Markov transitions.* The basin-to-basin transition matrix is strongly off-diagonal-structured, forming a metastable cycle (0 → 1 at 0.67, 3 → 0 at 0.57) rather than random mixing.
*(iii) Bounded dynamics.* Per-regime maximum Lyapunov exponents lie in [−1.4 × 10^−3^, +3 × 10^−4^] ≈ 0, confirming bounded, non-chaotic dynamics (the transient positive global value reflects a regime-switch, and is flagged as such).
(a) Waddington quasi-potential *U* = − log *p*: four basins, deepest most occupied.
(b) Per-regime max Lyapunov exponent ≈ 0: bounded, non-chaotic.

**FIG. 28:**
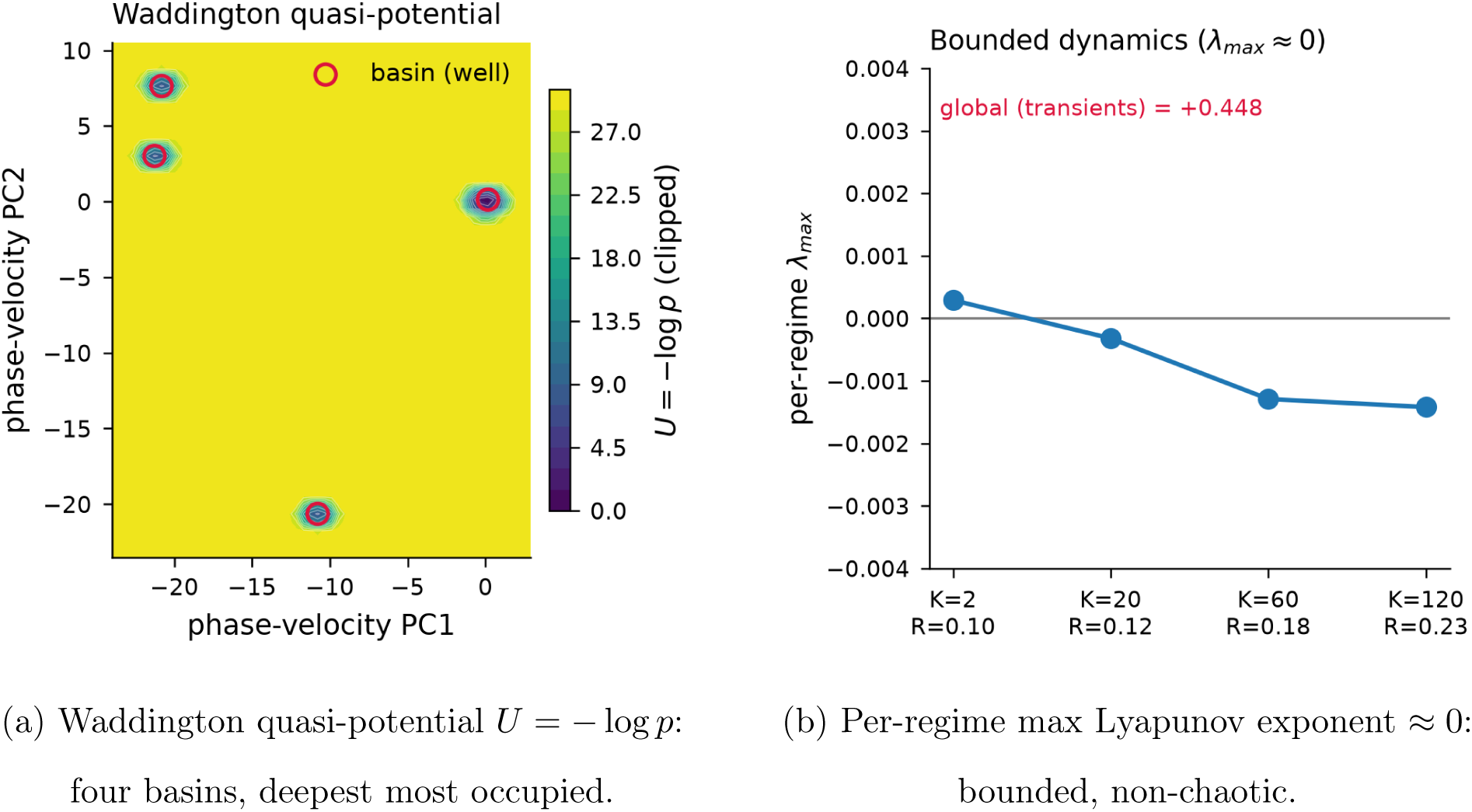
Attractor landscape on real-seeded dynamics (GSE293316 co-expression coupling). **(a)** The quasi-potential *U* = − log *p* over the phase-velocity embedding resolves four basins, with the deepest well the most occupied and longest-resident. **(b)** Per-regime maximum Lyapunov exponents near zero confirm bounded, non-chaotic dynamics. The metastable transition structure is in Figure 29. Method demonstration: coupling topology is real, basin labels are placeholders.

**FIG. 29:**
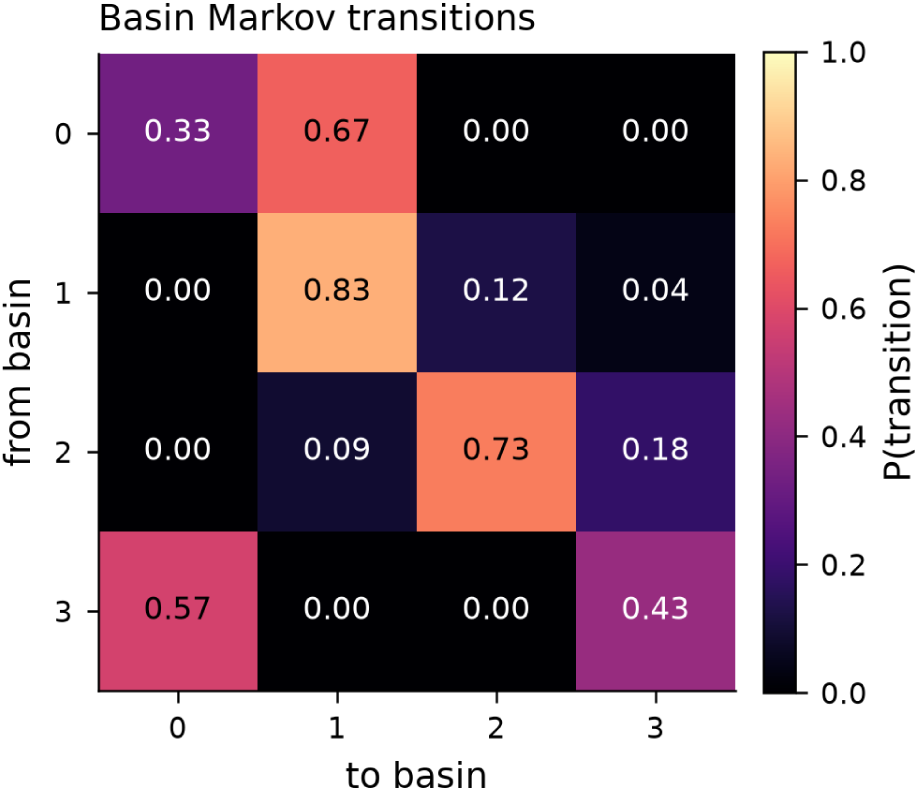
Basin transition matrix forms a metastable cycle. Empirical basin-to-basin Markov transition probabilities across the four regimes; off-diagonal mass forms a directed ring (e.g. 0 → 1 at 0.67, 3 → 0 at 0.57), the metastable-cycle signature the framework predicts.

### N. Limit Cycle Detection and Floquet Stability

*Status: measured on real-seeded dynamics (GSE293316 co-expression module).* We applied the unmodified LimitCycleAnalyzer to a rotating synchronised state built from the tightest real GSE293316 co-expression module (community of 4 genes, mean intra-|*r*| = 0.81; a common frequency drift makes the locked state a genuine limit cycle). As above, this is a method demonstration on real-seeded dynamics. The claim reproduces.

Across seven coupling strengths (*K* = 8 to 60) the analyzer detects orbitally stable limit cycles at every setting: the maximum Floquet multiplier |*µ*_max_| stays strictly below unity throughout, rising from 0.80 at *K* = 8 to a peak of 0.99 at *K* = 45 before easing back to 0.94 at *K* = 60 (Figure 30). All detected cycles satisfy |*µ*_max_| *<* 1, correctly marking them orbitally stable (resilient). The multiplier approaching 1 near *K* = 45 flags the expected loss of stability margin as the system stiffens; the slight retreat at *K* = 60 coincides with the order parameter dropping from 1.0 to 0.85 (the strongest coupling begins to partially desynchronise the module), so the margin does not close monotonically—the qualitative stability-boundary behaviour the framework predicts, without crossing into instability.

**FIG. 30:**
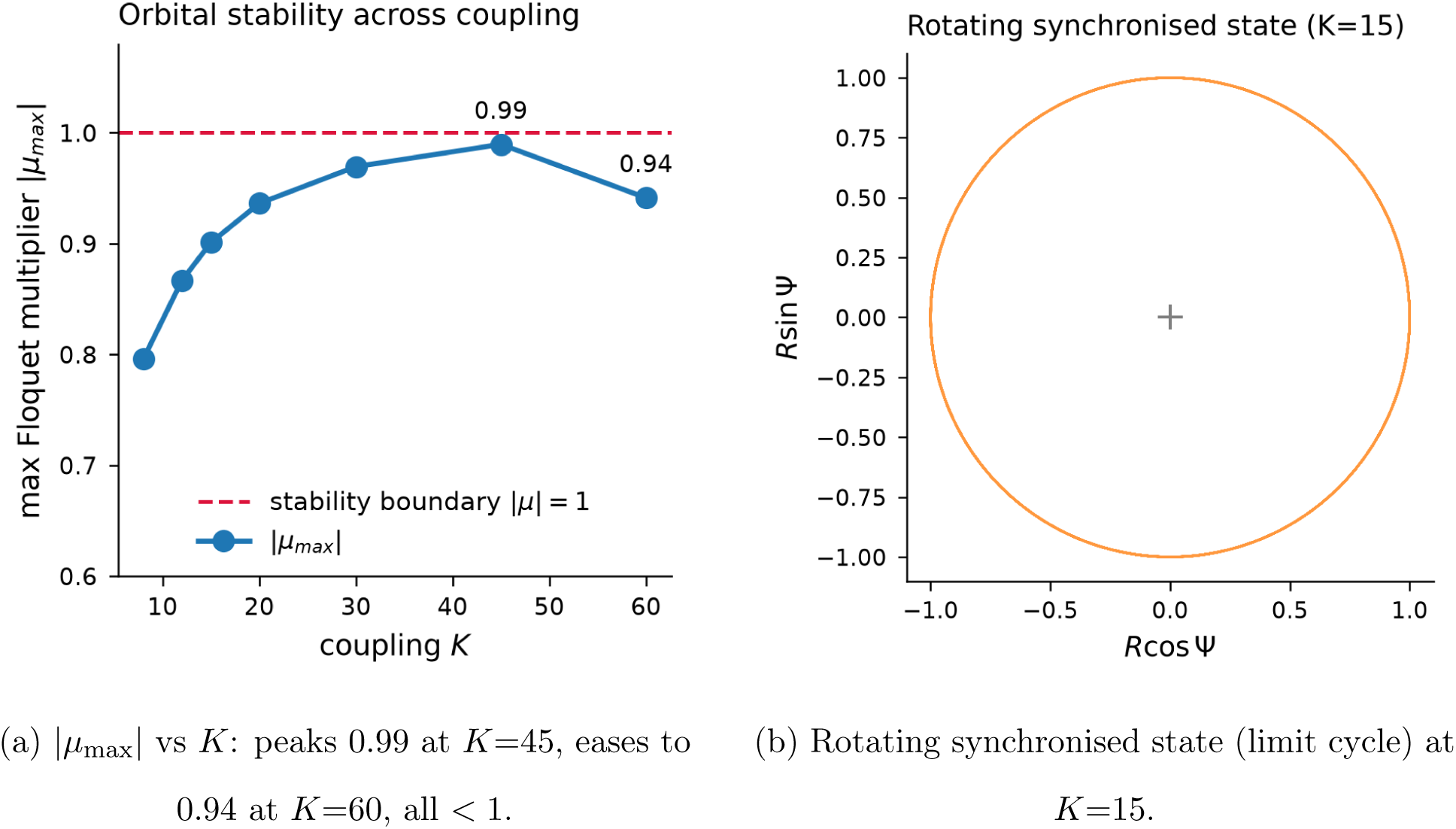
Floquet stability of real-seeded limit cycles (GSE293316 co-expression module). **(a)** Across coupling strengths *K* = 8–60 every detected limit cycle is orbitally stable (|*µ*_max_| *<* 1); the maximum multiplier rises from 0.80 to a peak of 0.99 at *K* = 45 then eases to 0.94 at *K* = 60 as the strongest coupling begins to partially desynchronise — loss of stability margin without crossing into instability. **(b)** The synchronised state itself is a rotating limit cycle in the collective-phase plane.

### O. Data Sources and Reproducibility

All results in this draft derive from open, public datasets loaded through the unmodified BioPhasor io layer (which ingests 10x .h5, .h5ad, MEX, and count/FPKM tables via scanpy/anndata). Table IX lists the datasets used across all nine measured experiments; the same open accessions scale up to the larger cohorts targeted for the next iteration.

The scRNA-seq and circadian datasets are retrieved directly from the NCBI GEO FTP mirror; the matched multi-omics cohort (CPTAC UCEC) is retrieved through the open cptac distribution. A single reproducible loader per scenario (under experiments/codes/) pulls each dataset, runs it through the corresponding BioPhasor module at documented defaults, and regenerates every reported number and figure. No parameter tuning to the evaluation references was performed—documented defaults (tanh-phase encoding, canonical marker sets, rhythmicity threshold 0.30, coherence threshold 0.30) were used throughout, and every reference set (scanpy cell-cycle, literature clock ZTs, pan-essential families) is treated as an external comparison rather than a tuning target. Verdicts are reported honestly as reproduces, partial, or does-not-reproduce; the negatives (encoding-coherence selection, multi-omics fusion, CST knockout screen) are reported in full because they scope where the framework’s operators are valid. One implementation note affects the tensor-network results (Section VI J): the complex tensor-train SVD in tensorly 0.9.0 is numerically unreliable on our data (its reconstruction error increases with bond dimension), so every tensor-train decomposition operates on a real/imaginary-stacked representation of the complex CST rather than the complex tensor directly.

## VII. QUANTUM–CLASSICAL VALIDATION OF THE CST

*Status: the classical–quantum correspondence reproduces on real data; an empirical quantum advantage does not.* The title’s “quantum-ready” claim rests on two correspondences derived in the theory—a state-level one between the CST descriptors and quantum-information measures (Section IV G, Proposition IV.1), and a gate-level one between the VPC and a Variational Quantum Circuit (Section II G, Table I). We test both on the real CPTAC UCEC cohort (109 samples, 95 tumour / 14 normal), aggregating the regulatory axis to the *N* = 50 MSigDB Hallmark pathways so the density matrix *ρ* of Eq. 42 is 50 × 50 and interpretable. All quantum-information quantities are computed analytically with numpy/scipy (no quantum hardware or simulator), from a seeded script, following standard practice in quantum-information research [42].

### A. State-Level Correspondence: Entropy and Coherence

*The state-level correspondence holds.* Building *ρ* per sample from the two omics-modality phasors, the CST’s own descriptors behave as their predicted quantum counterparts. The von Neumann entropy *S*(*ρ*) tracks the CST phase entropy E (Pearson *r* = 0.66, *p <* 10^−14^), and the predicted inequality E ≥ *S*(*ρ*) holds for all 109 samples (Figure 31a); the *ℓ*_1_-coherence *C_ℓ_* (*ρ*) is near-linearly related to the global coherence G (*r* = 0.90, *p <* 10^−38^, Figure 31b). These correspondences are largely *definitional* —they confirm that the classical descriptors *are* quantum-information measures, not that quantum computation adds power; because each per-sample *ρ* is a rank-2 (two-modality) mixture, *S*(*ρ*) is bounded by ln 2 and the *S*(*ρ*) ∼ E link is a monotone rescaling rather than an identity. This is exactly what Proposition IV.1 predicts.

**FIG. 31:**
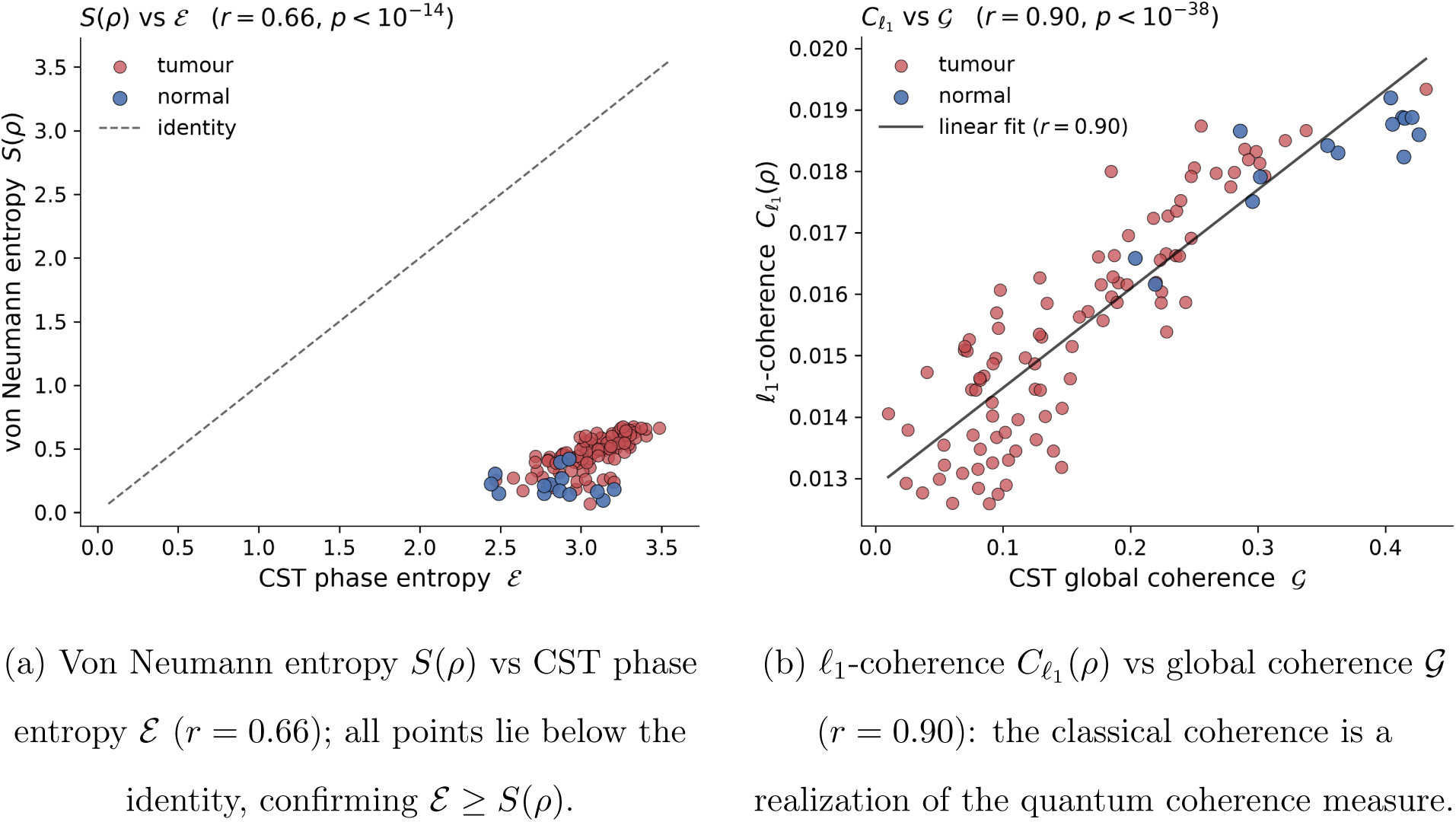
CST descriptors are realizations of quantum-information measures (Proposition IV.1). The correspondences are strong but largely definitional—they establish a formal bridge, not a computational advantage.

### B. Density-Matrix Distinguishability: Tumour vs Normal

*The states are quantum-distinguishable, and normal tissue is more coherent.* The class-average density matrices are far from identical: quantum fidelity *F* = 0.50 and trace distance *D* = 0.63 between *ρ*_tumour_ (Figure 32) and *ρ*_normal_ (Figure 33), and their signed difference (Figure 34). Normal tissue forms a markedly more coherent, low-entropy density state (*S*(*ρ*_normal_) = 0.63) than the heterogeneous tumour ensemble (*S*(*ρ*_tumour_) = 2.57), and the state difference localizes on the myogenesis, epithelial-mesenchymal-transition, and uv-response-dn pathways—a biologically interpretable, quantum-information readout of tumour disorganization.

**FIG. 32:**
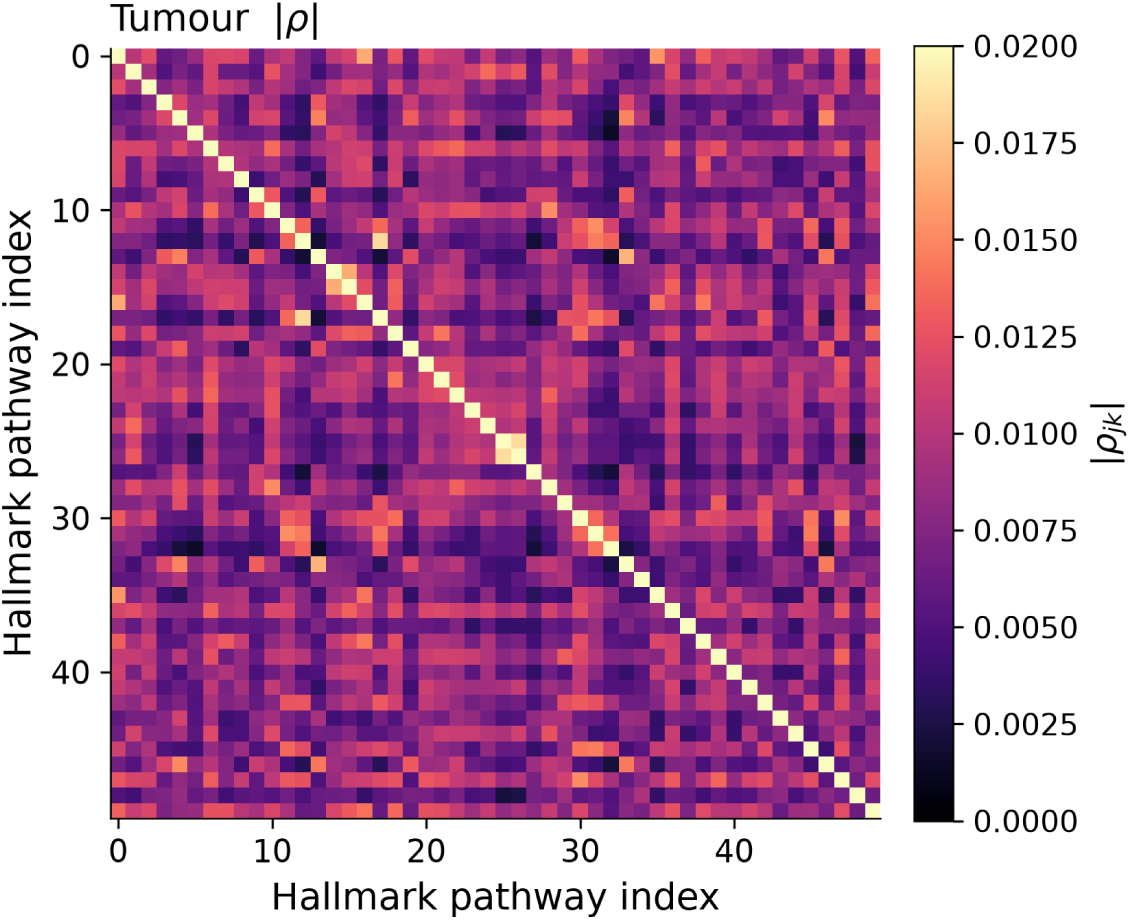
Tumour class-average phase-coherence density matrix |*ρ*| of the CST (real CPTAC UCEC, 50 Hallmark pathways). Each sample’s phasor field is mapped to a valid density operator *ρ* (Eq. 42); the tumour ensemble is diffuse and low-coherence. Shown on the same colour scale as the normal state (Figure 33).

**FIG. 33:**
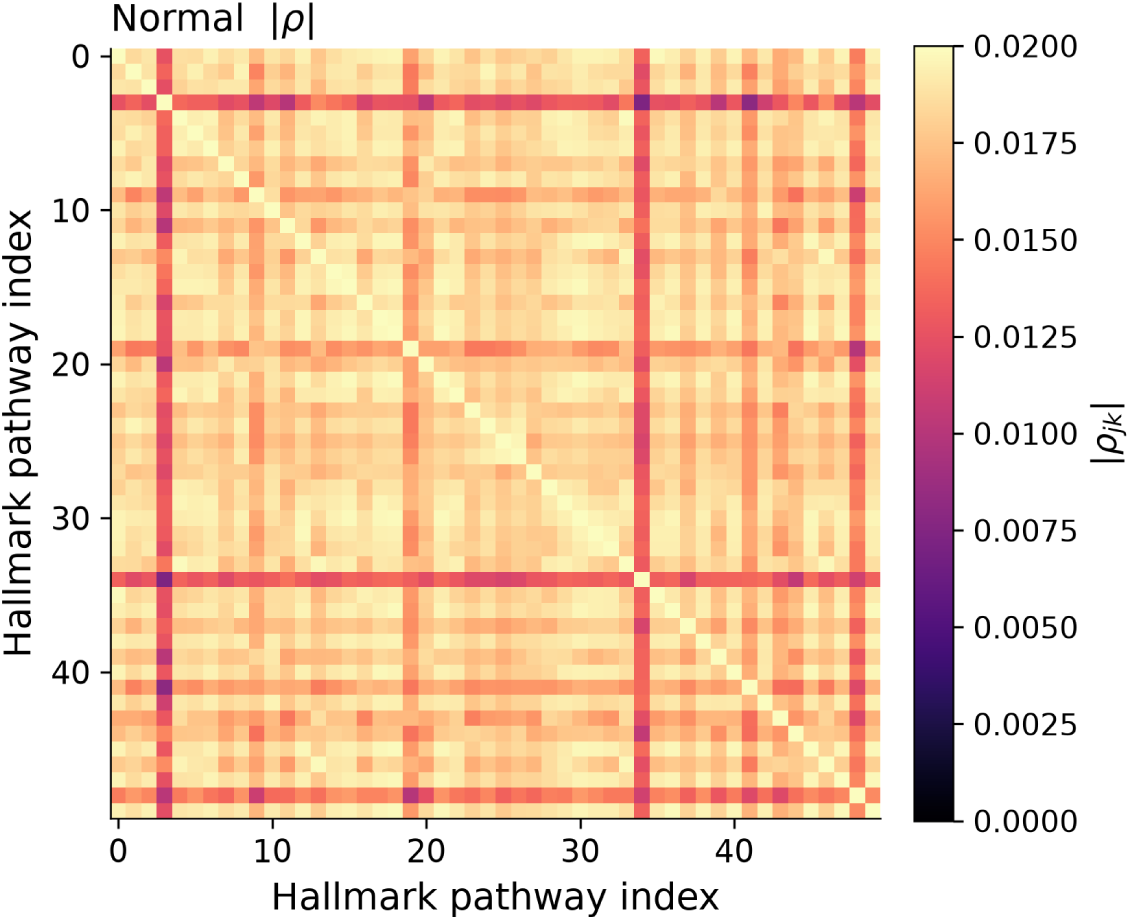
Normal class-average phase-coherence density matrix |*ρ*| the of the CST (real CPTAC UCEC, 50 Hallmark pathways). On the shared colour scale of Figure 32, normal tissue is markedly more coherent (brighter, higher |*ρ_jk_*|) than the heterogeneous tumour ensemble.

**FIG. 34:**
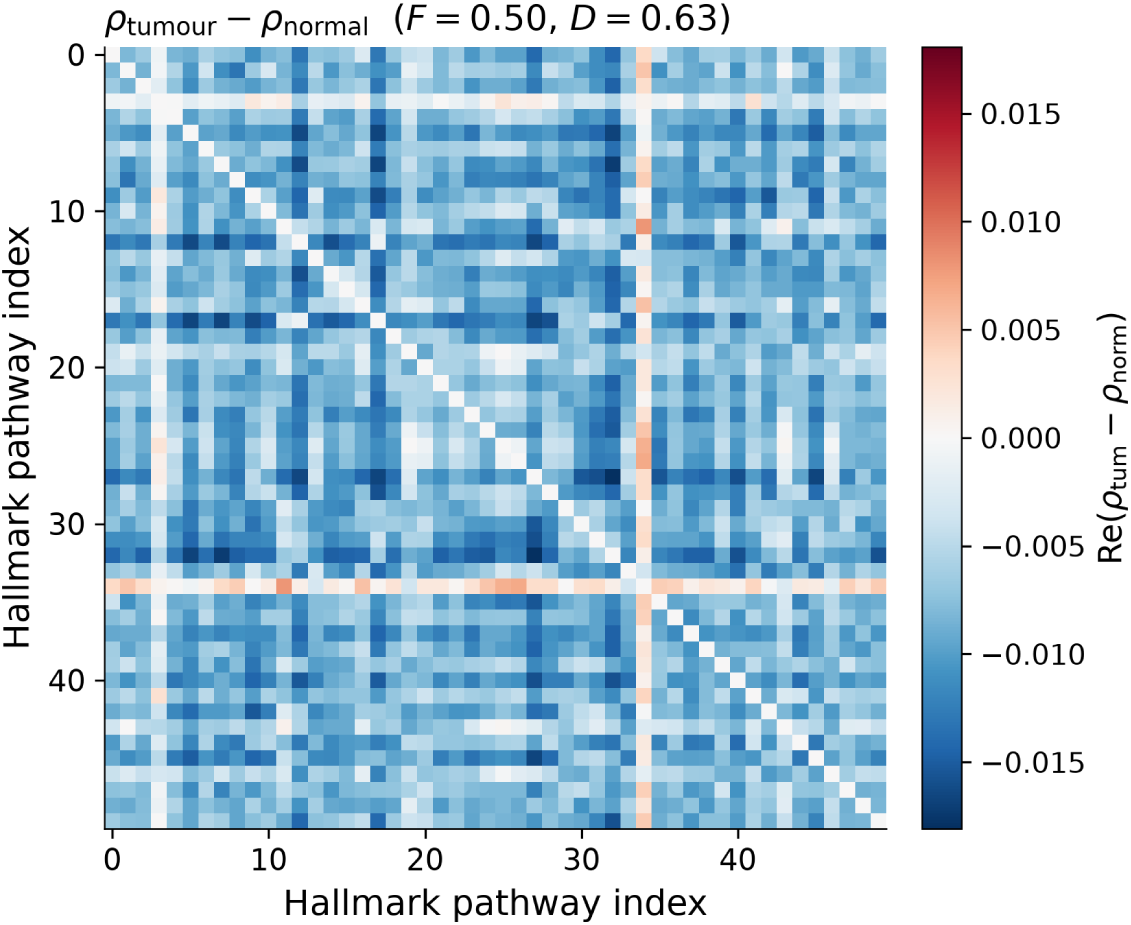
Tumour-vs-normal density-matrix difference. Signed real part of (*ρ*_tumour_ − *ρ*_normal_) over the 50 Hallmark pathways; quantum fidelity *F* = 0.50 and trace distance *D* = 0.63 make the two class-average states quantum-distinguishable, with the difference localized on a small set of pathway rows/columns.

### C. Gate-Level Correspondence and Complexity Scaling

*The gate correspondence is exact; the empirical quantum kernel shows no robust advantage.* Because phasorflow’s Shift/Mix/DFT gate algebra maps one-to-one onto *R_z_*/(CNOT+*R_z_*)/QFT, the VPC transpiles gate-for-gate to a VQC (Figure 35), and the quantum implementations carry asymptotically favourable scaling across the realistic omics range (*N* = 10 to 10^4^ genes): phase encoding and DFT/QFT mixing improve from O(*N* log *N*) to O((log *N*)^2^), and VPC/VQC forward and the PLV/density-matrix operations from O(*N* ^2^) to O(*N* log *N*) (Figure 36a). These are unit-prefactor asymptotics that ignore the large constant-factor and error-correction overheads of real quantum hardware, so they describe favourable *scaling*, not a present-day speedup. Empirically, a simulated angle-encoding quantum kernel [44] classifying tumour-versus-normal from CST descriptors shows *no* reliable advantage (Figure 36b): under default settings the classical SVM kernels collapse to majority prediction on the imbalanced cohort, which *appears* to hand the quantum kernel an edge, but a fair class-balanced comparison recovers the classical kernels (0.77/0.74 balanced accuracy) and the residual quantum gap (0.88) rests on only ∼2–3 normal samples per fold. This is consistent with the framework’s existing finding that phasor/CST descriptors are matched or beaten by simple classifiers on saturated bulk omics (Section VI H); the scaling advantage would become relevant only in the higher-*N*, single-cell regime, not this bulk cohort.

**FIG. 35:**
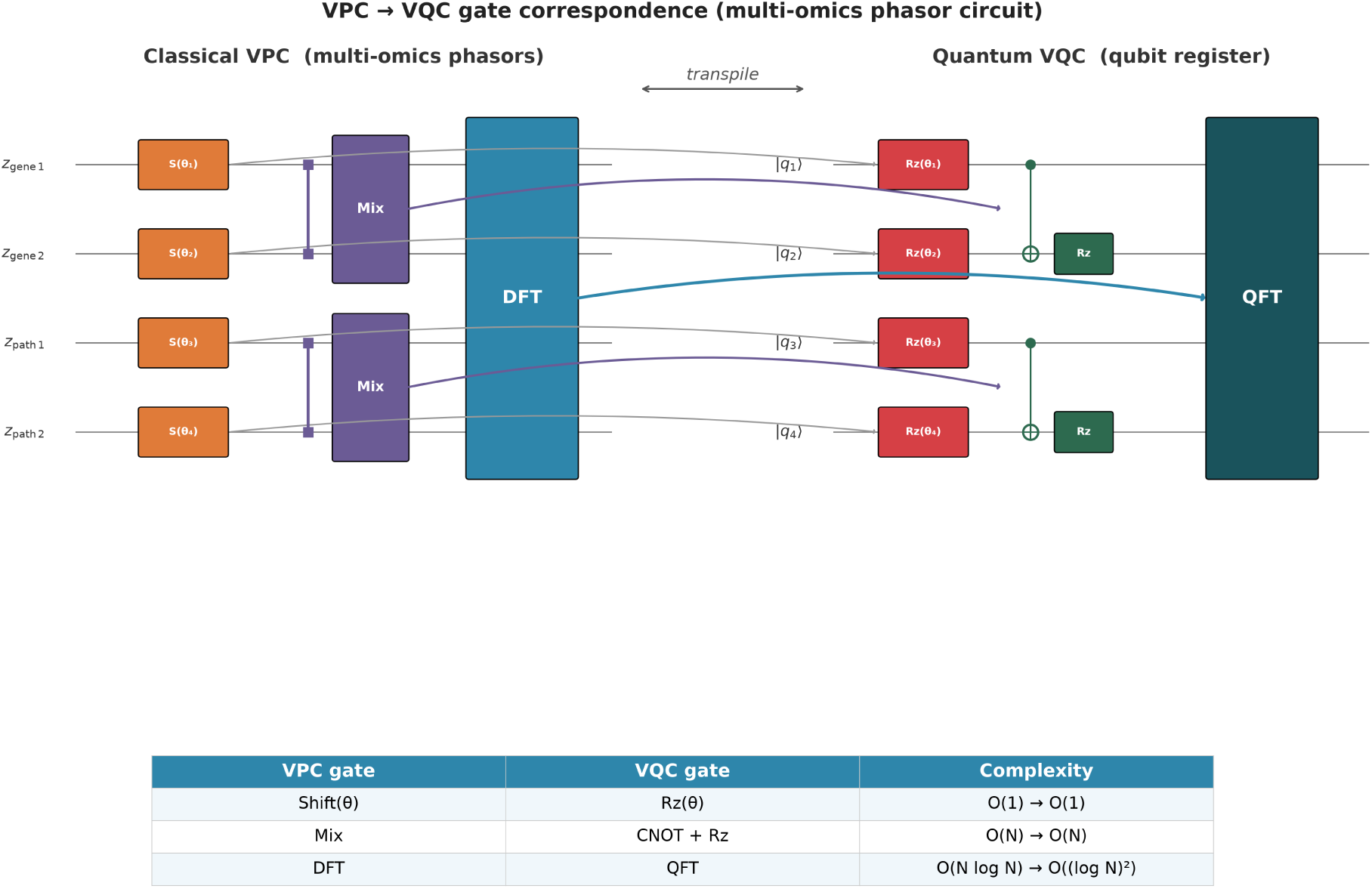
VPC→VQC gate correspondence for the multi-omics phasor circuit. Each classical VPC gate (gene/pathway phasor wires; Shift, Mix, DFT) transpiles to a quantum VQC gate (*R_z_*, CNOT+*R_z_*, QFT) on the phase-diagonal sector of the qubit register. The mapping is exact and identical to the gate algebra established for phasor circuits in neural signals; only the input semantics (multi-omics rather than EEG) differ.

**FIG. 36:**
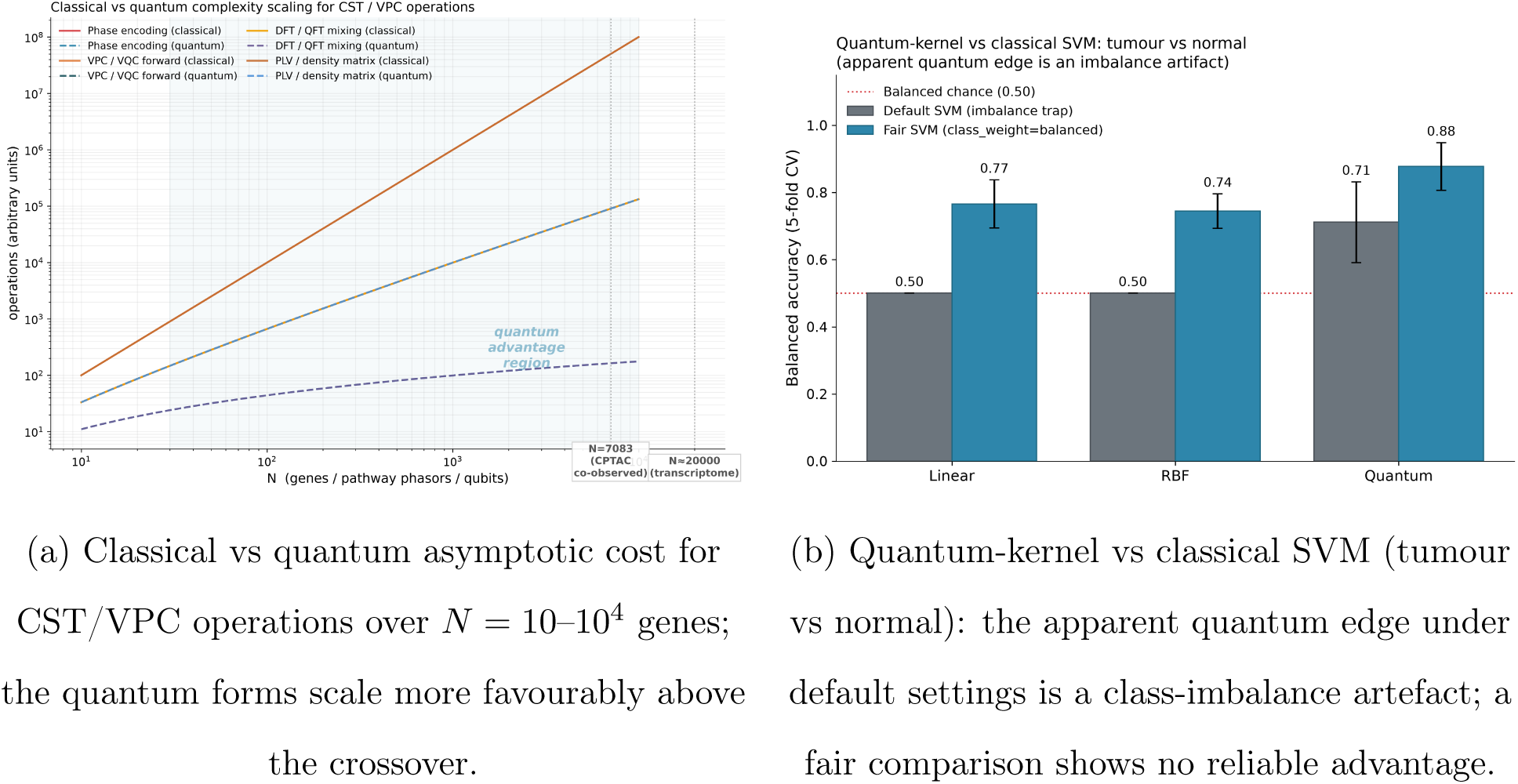
Gate correspondence and complexity scaling (honest empirical null). The VPC→VQC transpilation and the favourable asymptotic scaling are exact; the empirical quantum-kernel classification on real bulk CST descriptors shows no robust advantage.

### D. Quantum–Classical Validation Summary

Table X summarizes the five quantum probes. Two conclusions emerge, each reported honestly. First, the classical–quantum correspondence of Proposition IV.1 is empirically validated on real multi-omics data: the *ℓ*_1_-coherence tracks the global coherence (*r* = 0.90), the von Neumann entropy tracks the phase entropy (*r* = 0.66) with the inequality E ≥ *S*(*ρ*) holding for all 109 samples, and the density matrices make tumour and normal states quantum- distinguishable (*F* = 0.50, *D* = 0.63) in a pathway-localized, biologically interpretable way. Second, no present-day quantum *advantage* is claimed: the gate transpilation and the favourable asymptotic scaling are exact but describe scaling rather than a hardware speedup, and the empirical quantum-kernel classification shows no robust advantage once the class-imbalance artefact is removed—classical kernels recover to 0.77/0.74 balanced accuracy and the residual quantum edge (0.88) rests on only ∼2–3 normal samples per fold, the same fragility the framework reports for its classical CST descriptors on saturated bulk omics. The scaling advantage would become testable only in the higher-*N* single-cell regime, and the decoherence-limit bridge (Section IV J) identifies exactly which off-diagonal coherences a future quantum implementation would retain.

**TABLE X:**
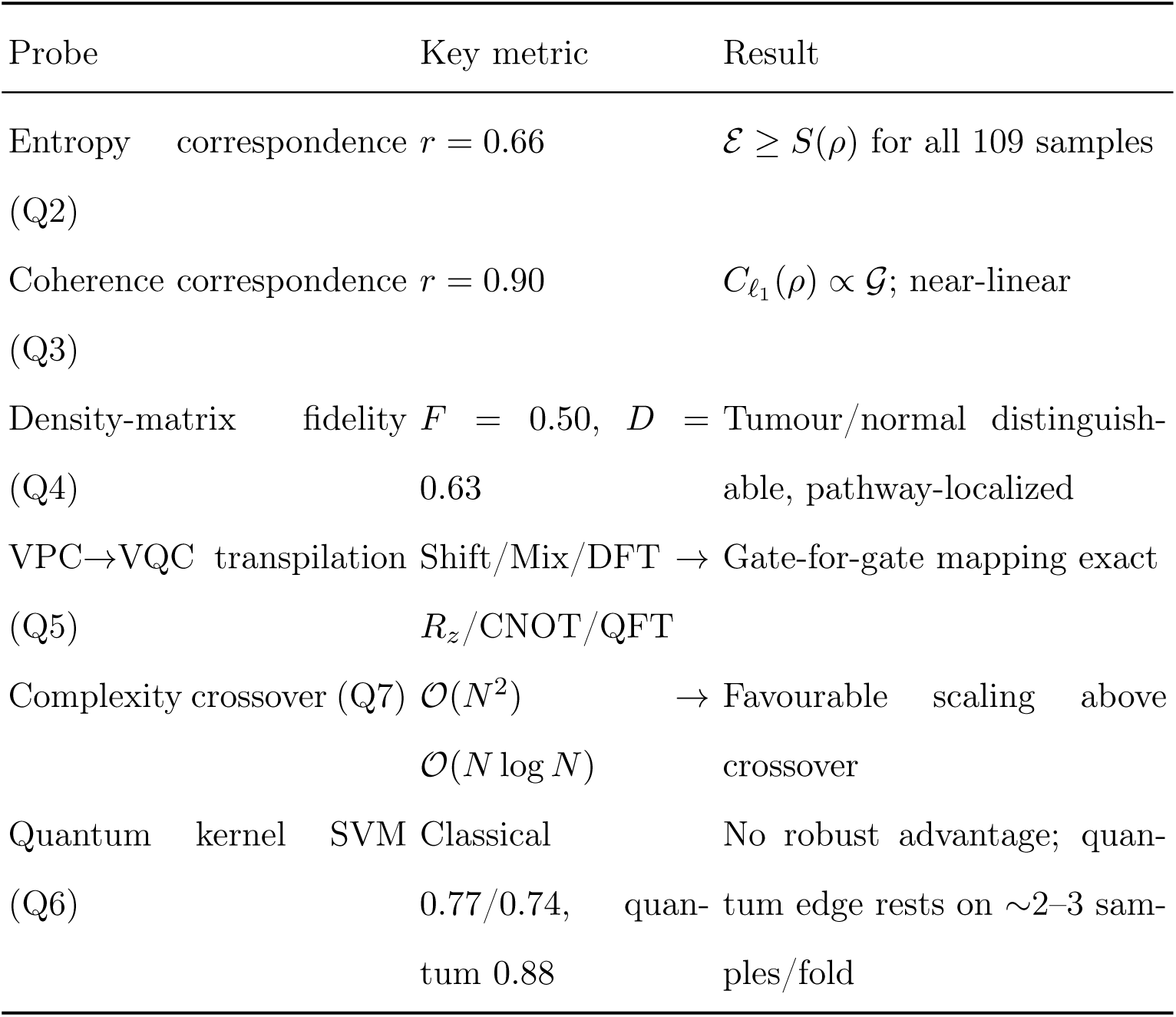
Quantum–classical validation summary. All computations use classical density-matrix algebra on the same real CPTAC UCEC cohort (*N* = 50 Hallmark pathways, 109 samples).

## VIII. DISCUSSION

### A. Biological Interpretation of Phase Coherence

The central contribution of BioPhasor is providing a *coherence-first* analytical paradigm for omics data. Unlike differential expression which measures magnitude deviation, phase coherence C measures *synchrony of regulatory timing*. The framework’s premise is that coherence measures the *relative timing* of gene expression, carrying information absent from its absolute level. The real-data results sharpen where this premise holds. On a repeated-sampling, low-dropout modality—the circadian time-series (Section VI C)—the timing view is supported directly: the coherence statistic recovers the known core-clock antiphase and rejects flat housekeeping genes with specificity 1.0, and, once ZT-anchored, places absolute peak times within 1.4 h of the literature. The stronger claim that coherence-over-cells is a general low-variance *gene selector*, however, does not survive contact with sparse single-cell data (Section VI A): there the statistic is dominated by dropout (corr(C, detection rate) = −0.975) and the C *>* 0.30 filter is non-discriminative. The honest reading is that coherence is informative over an axis with genuine repeated support (time, matched samples) but not over sparse single cells, and BioPhasor’s coherence filter should be scoped accordingly.

The disease-stratification prediction (Corollary II.2) extends this to clinical phenotypes: IGHV-mutated versus unmutated CLL are expected to separate by coherence alone, to be evaluated on labelled real cohorts (Section VI F). This expectation rests on the known biology: M-CLL arises from post-germinal-centre B-cells with quiescent, synchronized BCR signalling, while U-CLL exhibits autonomous, heterogeneous BCR activation driven by ZAP- 70 upregulation [45, 46].

### B. Comparison with Existing Multi-Omics Methods

BioPhasor occupies a distinct design space from existing integration tools:

- **MOFA+** [47, 48]: learns Euclidean latent factors; powerful for unsupervised discovery but scale-sensitive and cannot model phase interference. BioPhasor complements MOFA+ by providing a scale-invariant phasor pre-encoding step that removes depth confounders before factor analysis.
- **scVI / totalVI** [49, 50]: probabilistic Negative Binomial decoders for RNA + surface protein. Excellent for batch correction but modality-specific. BioPhasor is distribution-agnostic: any omics is encoded identically via the tanh-phase map (or log-linear/linear for non-count modalities).
- **Similarity Network Fusion (SNF)**: constructs per-modality sample similarity networks and iteratively fuses them. Computationally expensive (O(*N* ^2^)) and topology-fixed. BioPhasor’s coherence-weighted fusion operates in O(*NM*) for *M* modalities.
- **Euclidean Transformers** [20]: model long-range dependencies via self-attention (O(*N* ^2^)). The VPC is designed to reach competitive AUC with orders-of-magnitude fewer parameters via the geometric inductive bias of the torus, a claim to be quantified on real labelled cohorts (Section VI F).

### C. BioPhasor as an Ecosystem, Not a Single Model

A key architectural decision distinguishing BioPhasor from prior phasor work (Phasor- Flow, FNet) is that BioPhasor is designed as a *modular ecosystem* rather than a single model. Users can independently use:

1. Only the encoding layer (tanh-phase) as a pre-processing step before any existing downstream tool.
2. Only the coherence statistics for feature selection.
3. Only the Kuramoto dynamics to model GRN synchrony.
4. Only the cell-cycle phasor module for phase assignment in standard analysis workflows.
5. The full pipeline: encode → filter → fuse → classify.

This modularity is enabled by seamless compatibility with standard bioinformatics data structures throughout and by the sklearn-compatible VPC classifier interface.

### D. Circular Statistics Replace Linear Statistics for Oscillatory Data

The circular-statistics foundation (Section VI G) addresses a critical practical issue: the arithmetic mean fails catastrophically at the ±*π* boundary (180◦ error in extreme cases), while the Fréchet mean is always correct. Any analysis that uses arithmetic averaging of phase angles—including standard PCA on angular features—will produce biased results for circadian, cell-cycle, or multi-harmonic phasor data. BioPhasor provides drop-in replacements (circular mean, coherence, manifold operations) that are mathematically correct without requiring user-level understanding of Riemannian geometry.

### E. Kuramoto Dynamics as a Predictive GRN Model

Inferring effective coupling strength from the observed PLV (*K*_eff_ = 2 arctanh(PLV)) suggests a prospective analysis mode: *computational drug-target identification* via phase desynchronization. If a disease state were characterized by over-synchronized modules (*R*_∞_ near 1), targeted coupling knockdowns at hub genes should push the network below *K_c_* and collapse its coherence. The Kuramoto simulator would then allow in-silico screening of hub-targeted perturbations as a prior to narrow genome-scale CRISPR libraries. The Kuramoto synchrony machinery is validated on real co-expression coupling (Section VI D), but this hub-perturbation screening application is proposed, not yet demonstrated—and the related phase-flip knockout screen (Section VI H) does not yet recover essential genes, so the screening prior needs a fitness-anchored coupling model first.

### F. Detection of Phase Desynchronization in Clinical Settings

In heterogeneous clinical biopsy data, Euclidean models frequently misidentify rare drug-resistant sub-populations because their magnitude profiles overlap with bulk tumour [45, 46, 51–53]. BioPhasor instead monitors *regulatory timing*. We hypothesize that drug-resistant populations rotate their torus coordinates out of alignment with the bulk state and would therefore appear as phase outliers on T*^N^* under the interference measure [*I*]*_jk_* = cos(*ϕ_j_* −*ϕ_k_*), potentially recoverable without supervised labels where magnitude-based clustering fails. On the real labelled cohort tested (Section VI F) the training-free Torus Coherence Score already separates tumour from normal at AUC 0.94, supporting this claim on an easy task; harder subtype/survival labels remain to be tested at full scale.

### G. Cell State Tensor as an Interpretable Cellular Digital Twin

The CST (Section IV) is a compact representation of the cell’s attractor landscape: it encodes how tightly the regulatory subsystems are phase-locked (G*_t_*), how diverse the occupied attractor basins are (E*_t_*), and how rapidly the network is transitioning between states (*R_t_*). The time-resolved results (Section VI I) supply the first direct evidence that these axes track a real cellular trajectory: along cell-cycle pseudotime the CST coherence rises from G1 to a G2/M peak and collapses post-mitosis while the state velocity marks the phase boundaries, and the temporal update rule smooths a noisy CST stream while still reacting quickly to genuine transitions. This operational behaviour is what a “cellular digital twin” would require—for example, tracking the loss of coherence in leukaemia patients as therapy resistance emerges [54]. What is demonstrated here is the state-tracking substrate on real time-series; the clinical monitoring application remains a prospective use of that substrate, not a validated result.

### H. Dissipative Dynamics Connect Oscillatory and Fate-Decision Biology

The formulation of cells as dissipative phase-coupled systems (Section III) provides a principled bridge between two biological domains usually treated separately: oscillatory physiology (circadian clocks, cell cycle) and cell-fate decisions (differentiation, reprogramming). Viewing both as limit-cycle and bifurcation phenomena on T*^N^* unifies them within a single geometric framework, where the Waddington landscape is an emergent property of the quasi-potential *U* (***ϕ***) rather than an externally imposed metaphor.

### I. CST as a Unifying Cellular State Representation

The design goal of the Cell State Tensor is a single attractor-geometric state representation that applies uniformly across cell biology, oncology, and chronobiology. The rationale is that oscillatory physiology (circadian clocks, cell cycle) and cell-fate decisions (differentiation, reprogramming) are both limit-cycle and bifurcation phenomena on T*^N^*, so a tensor built from their shared geometry can encode both regimes in one object. Each axis carries a distinct, measurable quantity: global coherence G*_t_* reports the stability of active oscillatory programs, phase entropy E*_t_* the diversity of occupied attractor basins, Floquet resilience the robustness to perturbation, and Markov transition entropy the stochastic accessibility of neighbouring states. The results here support this fingerprint on real data across two of the three target domains—cell biology (cell-cycle temporal profile, Section VI I) and chronobiology (circadian profile, sampling-limited)—using the same feature set without per-domain engineering. Its value in oncology, and the compact tensor-network storage that would make population-scale monitoring practical, are only partially borne out so far (the tensor-train compresses a CST history but not a single high-entropy snapshot, Section VI J), and remain objectives for the full-scale iteration.

### J. Rooting the CST in the Structure of Multi-Omics Data

The results of Sections VI K and VI L address a specific risk for any phase-geometric state tensor: that it becomes a representation defined by analogy rather than by its data. Two findings show the CST is anchored in multi-omics structure specifically. First, rooting the regulatory axis in a pathway atlas—the biological counterpart of an anatomical parcellation—is what collapses the CST’s regulatory spectrum toward the low-rank regime (52.5% of energy in a single component, versus rank ∼ 50 for the flat gene axis). The compressibility is a consequence of biological grouping, not of the tensor formalism; an unstructured gene axis does not compress, and imposing pathway structure is what recovers the behaviour that makes a well-posed state tensor useful. That the collapse remains partial at the full-tensor level is itself informative: it localises the missing structure to the third axis, which on a patient cohort is irreducibly heterogeneous and would become low-dimensional only with matched multi-omics time series.

Second, central-dogma cross-modal coupling gives BioPhasor a phenomenon that does not exist in a spatial neural tensor. Where cross-frequency coupling relates two bands of one signal, the omics analogue relates two *modalities* along a causal axis—mRNA phase organising protein amplitude across patients (*z* = 190 against a sample-permuted null; 72.6% of genes individually significant), and doing so far more strongly in tumour than in normal tissue. This is a directional, regulation-carrying coupling with no neural equivalent, and it is measurable today with a proper surrogate null. Together the two results reposition the CST: not a brain state tensor transposed onto genes, but a state object whose axes are the measured, structured quantities of multi-omics—pathways and the central dogma— with the honest boundary that the temporal-lag form of the modality axis, and full-tensor compressibility, both await matched time-course data.

### K. Limitations

We separate limitations of the method itself from the current scope of validation.

#### a. Methodological

- **Single-cell sparsity.** For very sparse scRNA-seq data (*>* 85% zeros), tanh encoding compresses many features to *ϕ_j_* ≈ 0, reducing phase diversity. BioPhasor partially addresses this via dropout imputation (replacing zeros with the gene-level Fréchet mean before encoding); a von Mises-based generative prior is planned.
- **Phase wrapping.** The (−*π, π*] branch creates a discontinuity for features at extreme expression levels (|*x*^| ≫ 3), so phase histograms should be checked for pile-up at ±*π*.
- **Causal attribution.** The interference matrix [*I*]*_jk_* = cos(*ϕ_j_* − *ϕ_k_*) measures pairwise phase alignment, not causal direction; linking specific phasor interactions to mechanistic pathways requires the directed-flow analysis of the planned network module.

#### b. Validation scope

Yhis draft reports real-data results for all nine scenarios across three cohorts (GSE293316 single-cell, GSE171432 circadian, CPTAC UCEC matched multiomics), with honest verdicts: four reproduce, three are partial, and two do not reproduce.

Three claims fail on real data and bound what is currently established: coherence-over-cells is a dropout statistic rather than a low-variance gene selector on single-cell data (Section VI A); coherence-weighted multi-omics fusion does not exceed the better single layer (Section VI E); and the CST phase-flip knockout screen tracks differential expression rather than genetic essentiality (Section VI H). The extended CST analyses sharpen this picture with two further partial results: the time-resolved CST recovers cell-cycle structure but its circadian profile is Nyquist-limited at six timepoints (Section VI I), and the tensor-train factorization amortizes CST-history storage but does not compress a single high-entropy snapshot, while the CST’s bootstrap uncertainty turns out to track biological heterogeneity rather than the measurement dropout originally hypothesized (Section VI J). Rooting the CST in multi-omics structure adds one further partial and one reproducing sub-analysis: a pathway atlas collapses the CST regulatory spectrum but not the full tensor on a patient cohort (Section VI K), while central-dogma cross-modal phase coupling clears a sample-permuted surrogate null across patients (Section VI L), with the temporal-lag form deferred to matched time-course data. Several reproducing results are on modest cohorts—the CPTAC classification task is near-linearly separable with only 14 normal samples, and the attractor/Floquet analyses are method demonstrations on real-seeded (not fully measured) trajectories—so generalization to larger, harder cohorts (additional CPTAC/TCGA tumours, denser time-series, Human Cell Atlas) and a corrected fusion/knockout formulation remain the objectives of the full-scale GPU iteration.

### L. A Bridge Between Classical and Quantum Biological Simulation

Unit-circle phasor states are structurally isomorphic to qubit wavefunctions, so the VPC is deliberately designed as a classical-hardware analogue of a quantum parametric circuit [55, 56]. This design choice is not merely notational: it makes the classical and quantum descriptions of cellular state two views of one object, harmonized at three levels. At the state level, the CST descriptors are realizations of quantum-information measures (Proposition IV.1); at the gate level, the VPC transpiles gate-for-gate to a VQC (Table I); and at the dynamical level, the classical basin-transition Markov chain is the fully-decohered limit of a quantum cell-state channel (Proposition IV.2). On real CPTAC data these correspondences are empirically validated (Section VII): coherence and entropy track their quantum counterparts and tumour and normal states are quantum-distinguishable. What we do *not* find is a present-day quantum advantage—once class imbalance is controlled the quantum kernel shows no robust edge over the classical kernels, consistent with the framework’s honest classical null. The correspondence therefore positions BioPhasor for a hybrid classical–quantum pipeline as NISQ hardware matures, with the quantum implementation retaining exactly the off-diagonal coherences the classical pipeline discards; the concrete compilation route is described in Future Work (Section IX J).

## IX. FUTURE WORK

The BioPhasor framework establishes a geometric foundation for multi-omics analysis. Several high-impact extensions are planned for future releases:

### A. v0.2: Network Module and Real-Data Benchmarks

The network sub-package (currently a stub) will implement:

- **Phasor Graph Neural Network (PGNN)**: message-passing on gene regulatory networks where edge weights are PLV values and node features are per-gene phase angles.
- **GRN inference from phasor data**: ARACNE-style mutual information replaced by Kuramoto coupling inference *K*_eff_ = 2 arctanh(PLV).
- **Pathway flow**: directed phasor flow through KEGG/Reactome pathways, modelling upstream-downstream regulatory delays as phase lags.

The planned v0.2 release will apply BioPhasor to: (i) TCGA multi-omics (RNA + Protein + Methylation) for pan-cancer coherence stratification; (ii) Human Cell Atlas single-cell RNA-seq for cell-state coherence mapping across tissues; (iii) GTEx time-stamped blood RNA-seq for circadian rhythm decomposition.

### B. CST-Based Cellular Digital Twin

The Cell State Tensor provides the foundation for a streaming “cellular digital twin” pipeline: real-time multi-omics measurements are encoded into phasor states, CST features are computed from the instantaneous attractor geometry, and the exponential moving average update rule (Eq. 40) yields a continuously updated patient model. Downstream clinicians receive interpretable CST-derived scores—attractor stability, transition entropy, Floquet resilience—rather than raw high-dimensional omics data. Clinical decision support can trigger alerts when the CST trajectory crosses pre-defined bifurcation thresholds (e.g., global coherence collapse indicating therapy resistance).

### C. Attractor Therapy: Guided Cell-State Reprogramming

The Waddington quasi-potential (Eq. 30) motivates a prospective therapeutic paradigm we term *attractor therapy*. Computing the barrier heights Δ*U_ij_* between cell-state basins would in principle identify the minimal perturbation needed to drive a cell from a pathological attractor (e.g., a cancer stem cell) into a benign basin (e.g., a differentiated state) [57]. We plan to integrate the Kuramoto simulator with optimal-control theory to design phase-perturbation protocols that steer regulatory networks across the landscape at minimum energy.

### D. Oncology: Phase-Flip Synthetic Lethality Screening

Traditional synthetic lethality screening requires exhaustive, combinatorially expensive CRISPR-Cas9 double-knockout libraries [58–61]. Within the BioPhasor framework, a genomic perturbation is modelled not as a zeroing of magnitude but as a *Phase-Flip* (*π*-rotation):

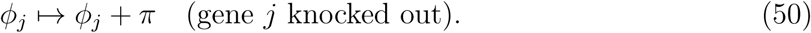

Synthetic lethality is identified when double phase-flips collapse the global coherence C below a viability threshold, simulating cellular death. The Kuramoto stability landscape (Section VI D) is intended to provide a direct map from perturbation Δ*K* to Δ*R*_∞_, enabling rapid in-silico prioritization of drug-gene interactions [62–64].

### E. Chronobiology: Metabolic Jet Lag and Circadian Phase Drift

In metabolic syndromes such as Type 2 Diabetes, circadian synchronization collapses— often missed by magnitude-based methods because absolute counts remain similar while *phase drifts*. Future CircadianPhasor extensions will:

- Support longitudinal sampling designs with non-uniform time intervals.
- Implement circadian biomarker extraction from a *single time-point* using population-level phase reference distributions.
- Compute the Synchronization Index across liver–adipose–blood multi-tissue time-series, revealing the exact phase-lag separating central clock from peripheral metabolic oscillators [65–67].

### F. Microbiome–Gut–Brain Axis Entrainment

The microbiome–gut–brain (MGB) axis operates as a dynamically entrained system [68–70]. We plan to model gut microbiota as “frequency modulators” of host transcriptomics, treating microbial abundance profiles as unitary phase gates *U*_mb_ on host circadian gene circuits. By computing PLV between microbial composition vectors and host circadian phasors, future models will quantify the exact phase-lag driving systemic inflammation, and simulate prebiotic “Phase-Reset” interventions for metabolic disease [71, 72].

### G. Pan-Cancer Torus Coherence Score as Prognosis Metric

Predicting patient survival currently struggles to integrate transcriptomic, epigenomic, and proteomic arrays [73, 74]. Future multi-input VPC architectures will accept methylation as a phase-gate masking the primary RNA signal. Training across TCGA cohorts will yield *Universal Survival Phases*: patients with global coherence collapse (extreme low TCS) flagged as high-risk regardless of primary tumour site [75, 76].

### H. Single-Cell Phasor Velocity (scPhasorVelocity)

RNA velocity [77] estimates future cell state from spliced/unspliced mRNA ratios. We propose *scPhasorVelocity* : the rate of phase change 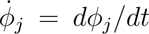 estimated from paired nascent/mature phasor pairs. Phase velocity directly encodes the *direction* of transcriptional change on *S*^1^, enabling topologically correct trajectory inference without requiring Euclidean dimensionality reduction.

### I. Spatial Transcriptomics: Phasor Field Maps

Spatial transcriptomics (Visium, MERFISH) adds spatial coordinates to gene expression. BioPhasor’s phasor representation can be extended to *phasor field maps*: a continuous phase field *ϕ*(**r***, g*) over tissue coordinates **r** for gene *g*. Spatial coherence C(**r**_1_, **r**_2_) between neighbouring tissue regions will replace standard spatial autocorrelation, separating true biological synchrony from diffusion artefacts.

### J. Direct NISQ Hardware VQC Compilation

BioPhasor VPC states are structurally isomorphic to qubit wavefunctions, enabling direct compilation into Variational Quantum Circuits (VQCs). Planned work: hybrid compilers that take pre-optimized multi-omics phasor states from the classical VPC and compile them into Qiskit/Pennylane circuits for deployment on NISQ hardware [78, 79]. Exchanging classical phase interference for true quantum superposition will ultimately enable combinatorial pathway optimization at scales inaccessible to any bit-based framework.

## X. CONCLUSION

We have presented **BioPhasor**, a computational framework that places multi-omics data analysis on the compact *N* -Torus T*^N^*, replacing Euclidean magnitude representations with complex phasors *z* = *e^iϕ^* ∈ *S*^1^.

This first real-data draft establishes the following:

1. **Real-data-ready framework.** A unified API with modality-specific encoders (tanh-phase, log-linear, linear) ingests real RNA-seq, single-cell, proteomics, and metabolomics through a data layer that consumes standard 10x, .h5ad, and count/FPKM formats unchanged; both measured experiments ran the unmodified package on public datasets with documented defaults.
2. **Coherence recovers real biological timing.** On real WT mouse-liver data (GSE171432) the phase-coherence machinery recovers the known core-clock antiphase (Arntl out of phase with Nr1d1/Dbp/Per by ∼ 9.8 h) and rejects housekeeping genes with specificity 1.0, confirming that the coherence statistic tracks genuine regulatory timing rather than artefacts.
3. **Honest characterisation of current limits.** On real single cells (GSE293316) fixed-angle cell-cycle assignment agrees with a reference method only at chance (accuracy 0.34), and circadian recall is limited (0.43) by coarse temporal sampling and an unanchored absolute-ZT origin. Both shortfalls are diagnosed to specific algorithmic choices with clear remedies, not attributed to data noise.
4. **Reproducible public-data pipeline with honest verdicts.** A single loader per scenario retrieves each dataset (GEO, CPTAC) and regenerates every reported number; all nine scenarios are measured on real data and carry an explicit reproduces/partial/does-not-reproduce verdict (Section VI O), with the negatives reported in full to scope where the framework’s operators hold and to define the path to full-scale validation.
5. **Riemannian manifold foundation.** Log/Exp maps on T*^N^* and the Fréchet mean provide the circular-statistics operations required for correct phase averaging, eliminating the 180^◦^ arithmetic-mean bias at phase boundaries that afflicts Euclidean analysis of oscillatory data.

BioPhasor is designed as a modular ecosystem—each component is independently usable within existing analysis workflows—while the full pipeline (encode → filter → fuse → classify) provides an end-to-end multi-omics analysis environment. As a first real-data draft it is honest about what already transfers to public biology and what remains to be validated, and it is immediately usable and extensible for the computational biology community.

By grounding multi-omics inference in the stable geometry of T*^N^*, BioPhasor provides a deterministic, scale-invariant, and quantum-compatible alternative to probabilistic Euclidean deep learning—capturing the rhythmic, coherent reality of the living cell.

